# Effects of neural noise on predictive model updating across the adult lifespan

**DOI:** 10.1101/2022.12.14.520501

**Authors:** Ina Bornkessel-Schlesewsky, Phillip M. Alday, Andrew W. Corcoran, Erica M. Wilkinson, Isabella Sharrad, Reinhold Kliegl, Richard L. Lewis, Steven L. Small, Matthias Schlesewsky

## Abstract

In the perceptual and sensorimotor domains, ageing is accompanied by a stronger reliance on top-down predictive model information and reduced sensory learning, thus promoting simpler, more efficient internal models in older adults. Here, we demonstrate analogous effects in higher-order language processing. One-hundred and twenty adults ranging in age from 18 to 83 years listened to short auditory passages containing manipulations of adjective order, with order probabilities varying between two speakers. As a measure of model adaptation, we examined attunement of the N400 event-related potential, a measure of precision-weighted prediction errors in language, to a trial-by-trial measure of speaker-based adjective order expectedness (“speaker-based surprisal”) across the course of the experiment. Adaptation was strongest for young adults, weaker for middle-aged adults, and absent for older adults. Over and above age-related differences, we observed individual differences in model adaptation, with aperiodic (1/f) slope and intercept metrics derived from resting-state EEG showing the most pronounced modulations. We suggest that age-related changes in aperiodic slope, which have been linked to neural noise, may be associated with individual differences in the magnitude of stimulus-related prediction error signals. By contrast, changes in aperiodic intercept, which reflects aggregate population spiking, may relate to an individual’s updating of inferences regarding stimulus precision. These two mechanisms jointly contribute to age-related changes in the precision-weighting of prediction errors and the degree of sensory learning.

## 1 Introduction

Ageing is associated with increasingly noisy information processing in the human brain. In addition to being reflected in higher intra- and inter-individual variability of behavioural responses, this has been linked to less effective neuromodulation (e.g. Li et al., 2001) and a concomitant attenuation of neural gain (Aston-Jones & Cohen, 2005; Eldar et al., 2013; Li et al., 2000; Li & Sikström, 2002). Gain modulation shapes the activation function of a neuron, with higher gain increasing the unit’s response to a relevant stimulus and decreasing its response to an irrelevant stimulus, thus resulting in a more binary response function and stronger stimulus discriminability (Aston-Jones & Cohen, 2005). Increases in gain result from the release of catecholamines (dopamine, noradrenaline) (Servan-Schreiber et al., 1990), neuromodulatory systems which decline with age and which have been linked to age-related changes in information processing (see Li & Rieckmann, 2014, for a review). Computational modelling has demonstrated that neuromodulation-induced changes in neural gain can account for a wide range of age-related changes in cognition, including in learning rate, interference susceptibility and complexity cost (Li et al., 2000), as well as decreases in processing fidelity / increases in processing variability (Li & Rieckmann, 2014).

Somewhat more recent approaches have linked neural noise in ageing to changes in aperiodic (1/f) activity of the human electroencephalogram (EEG; Voytek et al., 2015). In contrast to neural oscillations in frequency bands such as alpha (*∼* 8-12 Hz), theta (*∼* 4-7 Hz) and beta (*∼* 15-30 Hz), aperiodic activity defines the overall shape of the EEG’s power spectral density (PSD). Logtransformed aperiodic activity approximates a 1/f power law (He, 2014), the functional form of which can be summarised with a linear model defined by two parameters: the aperiodic *offset* or *intercept*, which reflects the amount of power predicted for the lowest frequency bin included in the fitted model; and the aperiodic *slope* or (negative) *exponent*, which reflects the steepness of the power decay from lower to higher frequencies (Donoghue et al., 2020; Voytek & Knight, 2015).

Flatter aperiodic slopes are thought to reflect less synchronous (“noisier”) neural activity, due to increased levels of aberrant or background firing not tied to an oscillatory carrier frequency (Voytek et al., 2015; Voytek & Knight, 2015). As argued by Voytek and Knight (2015), a certain level of neural noise is beneficial, serving as a control mechanism to protect against hyper- synchonisation (Radman et al., 2007), which can result when lower frequency oscillations bias spiking activity through phase-amplitude coupling (Canolty & Knight, 2010), thus leading to increases in local population firing rate and elevated local field potentials (LFPs). From this perspective, neural noise is defined as temporally decorrelated spiking during non-preferred oscillatory phases. While very low levels of neural noise can result in an exaggerated state of overcoupling in pathological cases (e.g. in Parkinson’s disease), too much neural noise is also detrimental to information processing (e.g. in schizophrenia or autism) (see Voytek & Knight, 2015, for further details). Changes in aperiodic slope may also reflect a shift in the balance between excitatory and inhibitory activity (Gao et al., 2017), which is in line with the proposal that individuals with autism and schizophrenia show altered excitation-to-inhibition ratios (Rubenstein & Merzenich, 2003) and aperiodic slope characteristics (Manyukhina et al., 2022; Molina et al., 2020) in comparison to neurotypical individuals. In contrast to the aperiodic slope, the aperiodic intercept has been shown to reflect aggregate population spiking activity across widespread brain regions (Manning et al., 2009) and correlates with the fMRI BOLD response (Winawer et al., 2013).

Returning to the relationship between age and aperiodic activity, ageing is associated with a flattening of the aperiodic slope and a downshifting of the aperiodic offset (Donoghue et al., 2020; Merkin et al., 2023; Voytek et al., 2015; Waschke et al., 2017). Flatter aperiodic slopes in older adults correlate with decreased behavioural performance in visual working memory tasks (Voytek et al., 2015), less consistent neurophysiological responses (peak alpha inter-trial coherence, ITC) to visual stimuli (Tran et al., 2020) and higher irregularity (entropy) of EEG activity in an auditory discrimination task (Waschke et al., 2017). Thus, as for the gain-related perspective on neural noise, higher levels of neural noise as measured by an age-related flattening of the aperiodic slope are accompanied by increasing deficits in perceptual and cognitive processing.

Recent research in fact points to a compelling link between the gain-related and aperiodicactivity-related perspectives on neural noise and ageing. Pertermann et al. (2019) demonstrated a strong correlation between on-task changes in aperiodic slope and pupil dilation, an established marker of noradrenergic activity, when response inhibition was required during a Go/NoGo task. This could be taken to suggest that the lower neural gain associated with higher neural noise leads to a less effective discrimination of relevant versus irrelevant information, with potential implications for a wide range of behavioural tasks (cf. Bornkessel-Schlesewsky et al., 2022).

A possible implication of these neurophysiological correlates of ageing that has hitherto not been examined is that they may be accompanied by systematic age-related changes in predictive model updating. Recent work on individual differences in predictive processing in young adults suggests that individuals with steeper resting aperiodic slopes adapt their predictive models more rapidly to novel input during language comprehension (Bornkessel-Schlesewsky et al., 2022). Steeper on-task aperiodic slopes have likewise been argued to accompany stronger predictive language processing in both younger and older adults (Dave et al., 2018) and to be conducive for language learning in adulthood (Cross et al., 2022). These findings provide an initial indication of how the results on neural noise, aperiodic slope and gain control could be linked to the prominent literature on predictive coding and active inference as a possible unified theory of sentient behaviour (Friston, 2005, 2009; Parr et al., 2022). On this view, perception is underwritten by unconscious inference, whereby the brain inverts an internal (generative) model to infer the hidden causes of its sensory states. The goal is to maximise model evidence (i.e. the marginal likelihood of sensory data) by minimising the discrepancy between predicted and observed sensory input (i.e. prediction error) in an approximately Bayes-optimal fashion. This is accomplished by propagating prediction errors from lower to higher levels of a hierarchically-organised cortical architecture and using these error signals to continuously update beliefs at higher levels.

Crucially, the propagation of prediction errors up the cortical hierarchy is determined by prior beliefs about the *precision* (inverse variance) of sensory input. Informative sources of sensory data are accorded greater precision than ambiguous or unreliable sources of input, thus compelling stronger updates of prior beliefs. Notably, evidence suggests that older adults give greater weight to top-down model predictions vis-à-vis incoming sensory evidence (Chan et al., 2021; Moran et al., 2014; Wolpe et al., 2016), thus reducing the extent to which new information is assimilated within the model. Moran et al. (2014) suggest that this age-related shift away from sensory learning reflects an optimisation of the neural architecture for predictive modelling: by relying more strongly on established internal models of the world that have been refined over the course of a lifetime, older adults avoid excessive model complexity and overfitting. This may be beneficial for dealing with input that is noisier in older adulthood due to sensory degradation, and for avoiding the complexity cost that is associated with model updating (Zénon et al., 2019).

We propose that the flattening of aperiodic slope may provide a neurophysiological mechanism to underpin this age-related change in cognitive processing. The reduction of neural gain that accompanies a flattened slope prevents overly rapid model updating, thereby serving a protective function against exaggerated model complexity and overfitting in old age. This proposal thus posits a cognitive counterpart to the potential protective physiological function of neural noise as discussed by Voytek and Knight (2015).

Here, we test the hypothesis that older adults adapt their predictive models more slowly to novel input environments and examine how this relates to aperiodic activity. We employed the same design as in Bornkessel-Schlesewsky et al. (2022, Experiment 2) with a sample of 120 participants spanning the adult lifespan (18 to 83 years of age). Participants listened to 150 short (4-5 sentence) passages recorded by two male speakers. Embedded in each passage were two two-adjective noun phrases (e.g. “huge grey elephant”), with one speaker producing a higher (70:30%) ratio of expected-to-unexpected adjective orders (“huge grey” vs. “grey huge”) and the ratio being reversed for the other speaker. Using a novel measure of “speaker-based surprisal” first introduced in Bornkessel-Schlesewsky et al. (2022), we tracked the expectedness of adjective orders within the experiment as they unfolded on a trial-by-trial basis. As in our previous study, our primary outcome measure was the N400 event-related potential (ERP) as a presumed index of precision-weighted prediction error in language processing (Bornkessel- Schlesewsky & Schlesewsky, 2019). We interpret the extent to which N400 amplitude at the position of the critical second adjective attuned to speaker-based surprisal across the experiment (i.e. the strength of the relationship between N400 amplitude and speaker-based surprisal) as an indicator of predictive model adaptation to the novel linguistic environment provided by the experimental context.

We predicted that the attunement of N400 amplitude to speaker-based surprisal over the course of the experiment would be reduced with increasing age. We further expected that the relationship between the N400 and speaker-based surprisal would be additionally modulated by aperiodic activity, such that older adults with a steeper aperiodic slope would show stronger model adaptation effects than those with a flatter slope.

## 2 Results

In addition to the main language processing task, participants were exposed to a passive auditory oddball paradigm. The mismatch negativity (MMN) elicited in such paradigms has been discussed extensively in relation to predictive model updating (e.g. Garrido et al., 2009) and associated changes across the adult lifespan (Moran et al., 2014). We thus utilised ERP estimates derived from the oddball paradigm as individual differences measures for the language processing task, supplementing resting-state aperiodic slope and intercept as our main individual differences measures of interest. We also examined peak alpha frequency (Individual Alpha Frequency, IAF) and Idea Density (ID), to allow for comparisons with our previous work on young adults (Bornkessel-Schlesewsky et al., 2022), as well as measures of cognitive ability and language proficiency.

In the following, we first discuss the individual differences measures, before turning to the results for the main language processing task.

### 2.1 Individual differences measures

#### 2.1.1 Auditory oddball paradigm

In a first step, we analysed the MMN and P3 ERP components elicited within the passive auditory oddball paradigm. The main rationale for doing so was to use these as individual differences measures for the magnitude of predictive auditory model updating (MMN) and neural gain control (P3), respectively. The MMN has been linked to precision-weighted prediction error signals in auditory processing (Todd et al., 2014; Todd et al., 2011; Todd et al., 2013) and was used by Moran et al. (2014) in their analysis of changes in predictive model updating across the adult lifespan. By contrast, the P3 has been associated with activation of the noradrenergic system and can thus serve as a proxy for phasic changes in gain control engendered by the release of noradrenaline from the brainstem locus coeruleus (Nieuwenhuis et al., 2005). Note that, according to Nieuwenhuis et al. (2005), this is the case for both the target-related, posteriorly distributed P3b and the more anterior P3a elicited in a passive oddball paradigm such as the one employed here.

Grand average ERPs for the oddball paradigm are shown in Figure 1. The figure shows the expected biphasic MMN-P3 pattern for oddball versus standard tones across all three age groups.

**Figure 1:**
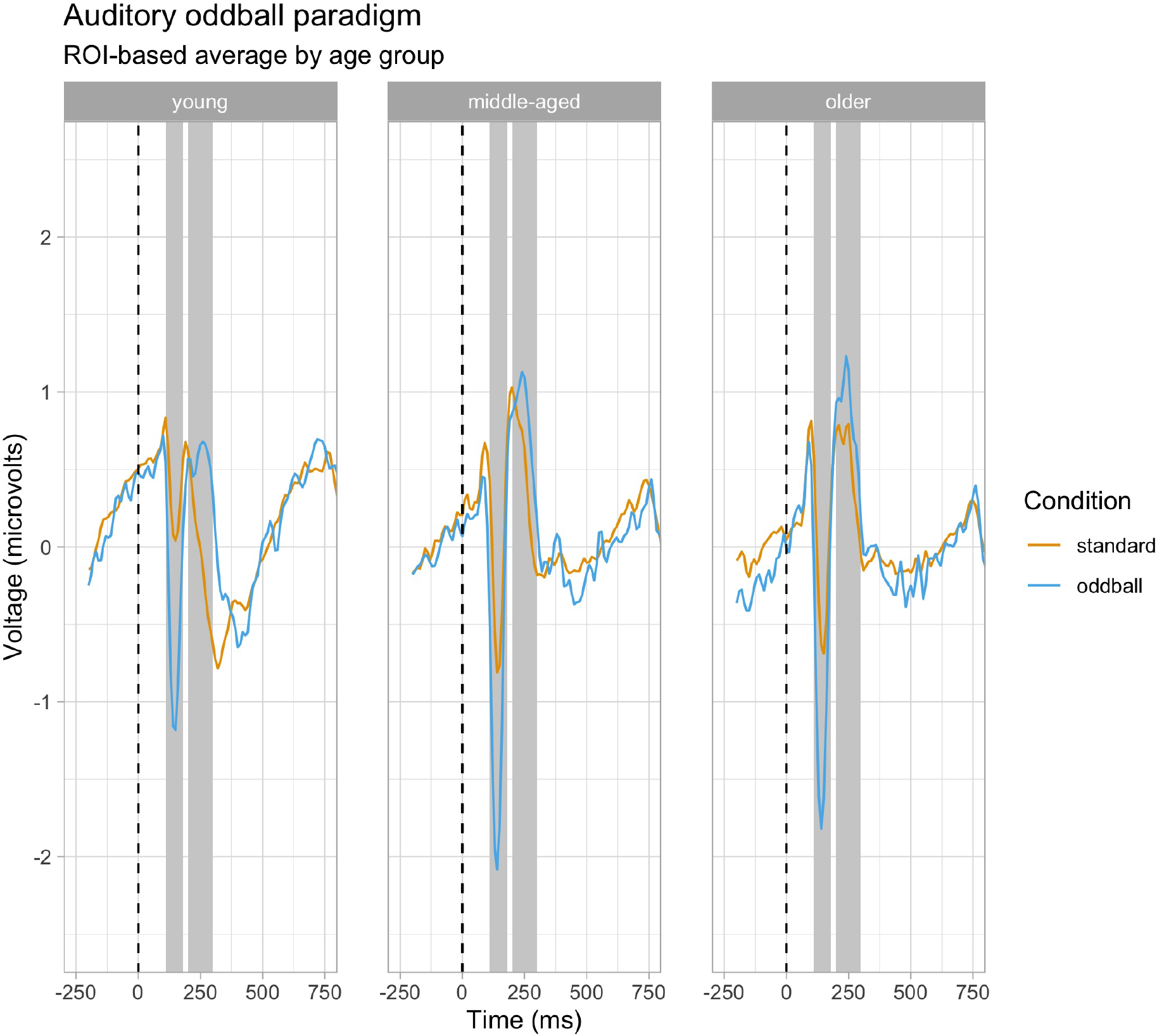
Grand average ERPs elicited by the passive auditory oddball tasks. ERP traces are averaged within the fronto-central region of interest (ROI) used for analysis. Tone onset occurred at the dashed vertical line and positivity is plotted upwards. Facets show the ERP responses by age group. Grey shaded areas indicate the time windows selected for analysis of the MMN and the P3, respectively.

For the analysis of the oddball data, we first examined the behaviour of the MMN and the P3 by investigating effects of Age and Epoch (as a proxy for time on task). To this end, we calculated linear mixed-effects models (LMMs) with fixed effects of Prestimulus amplitude, Condition (standard vs. oddball), Age, Epoch, and their interaction. Random effects structures were determined using the parsimonious model selection procedure described in section 4.8.1.

For the MMN, the final model included random intercepts by Participant and Channel as well as by-participant random slopes of Prestimulus Amplitude and Condition. This model revealed an interaction of Condition x Epoch x Age (Estimate = 0.0452, Std. Error = 0.0150, *z* = 3.03, *p* = 0.0025), which is visualised in the left-hand panel of Figure 2.

**Figure 2:**
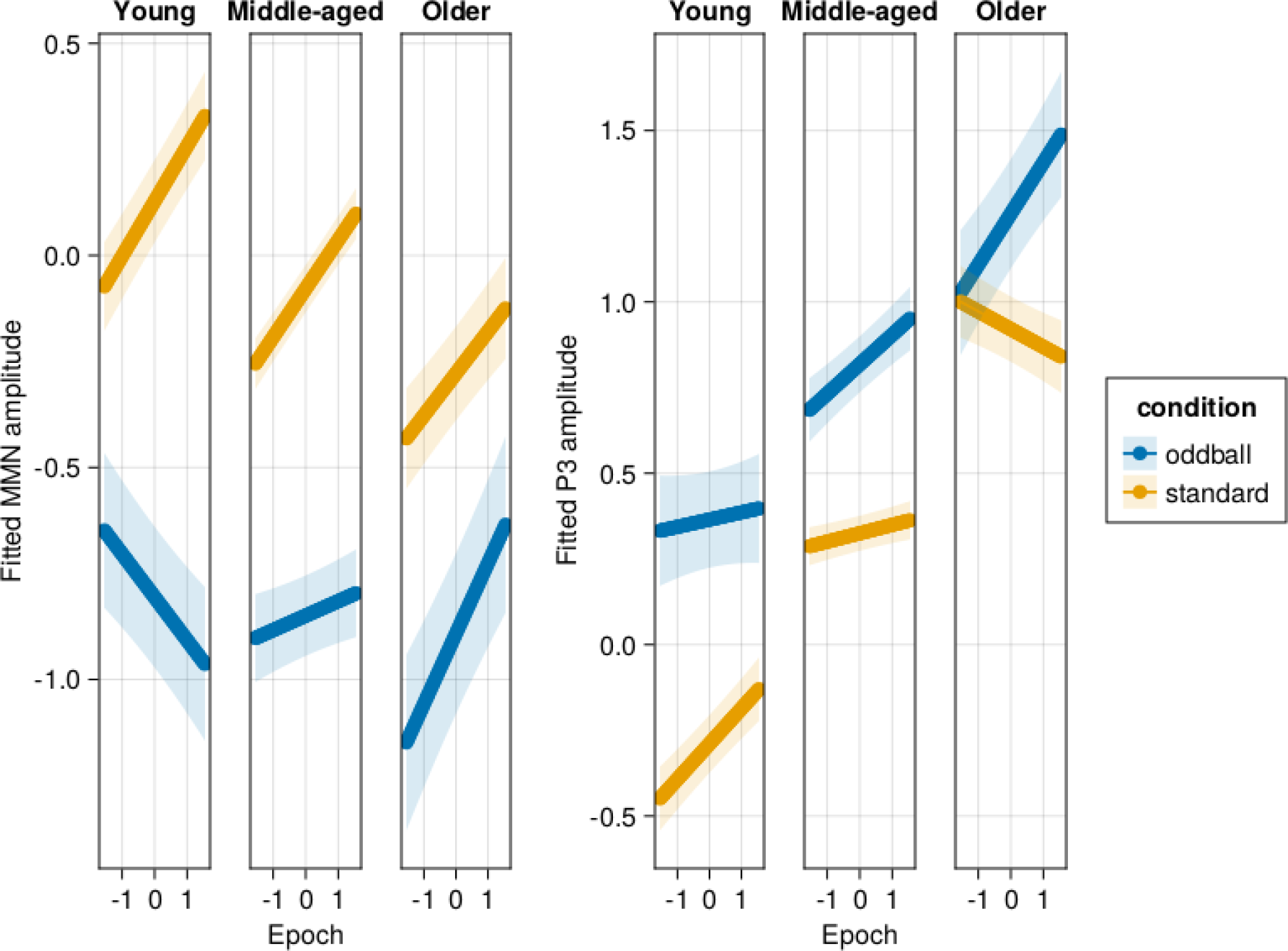
Effects of age and epoch on MMN (left panel) and P3 (right panel) amplitude in the auditory oddball task as modelled. The facets show fitted effects for young adults (estimated for age 18), middle-aged adults (estimated for age 51) and older adults (estimated for age 83), respectively. Note that the trichotomisation of age is for visualisation purposes only, with age included in the statistical models as a (standardised) continuous predictor. Shaded ribbons signify standard errors.

For the P3, the final model included random intercepts by Participant and Channel as well as by-participant random slopes of Prestimulus Amplitude and Condition and by-channel random slopes of Prestimulus Amplitude. Again, there was a significant interaction of Condition x Epoch x Age (Estimate = 0.0429, Std. Error = 0.0155, *z* = 2.76, *p* = 0.0058). This interaction is visualised in the right-hand panel of Figure 2.

To extract by-participant MMN and P3 effect estimates as individual differences predictors, we ran mixed models on amplitudes averaged over the electrodes in the ROI of interest and over epoch for the MMN and P3 time windows, respectively. Prestimulus amplitude, Condition and their interaction were included as fixed effects. The final, parsimoniously selected random effects structure included a random intercept of participant and by-participant random slopes of Prestimulus amplitude and Condition. We extracted individual fitted MMN and P3 effects from this analysis for inclusion in the main mixed model analysis.

#### 2.1.2 Distribution of and correlations between individual differences measures

The distribution of (z-transformed) individual differences variables is visualised in Figure 3.

**Figure 3:**
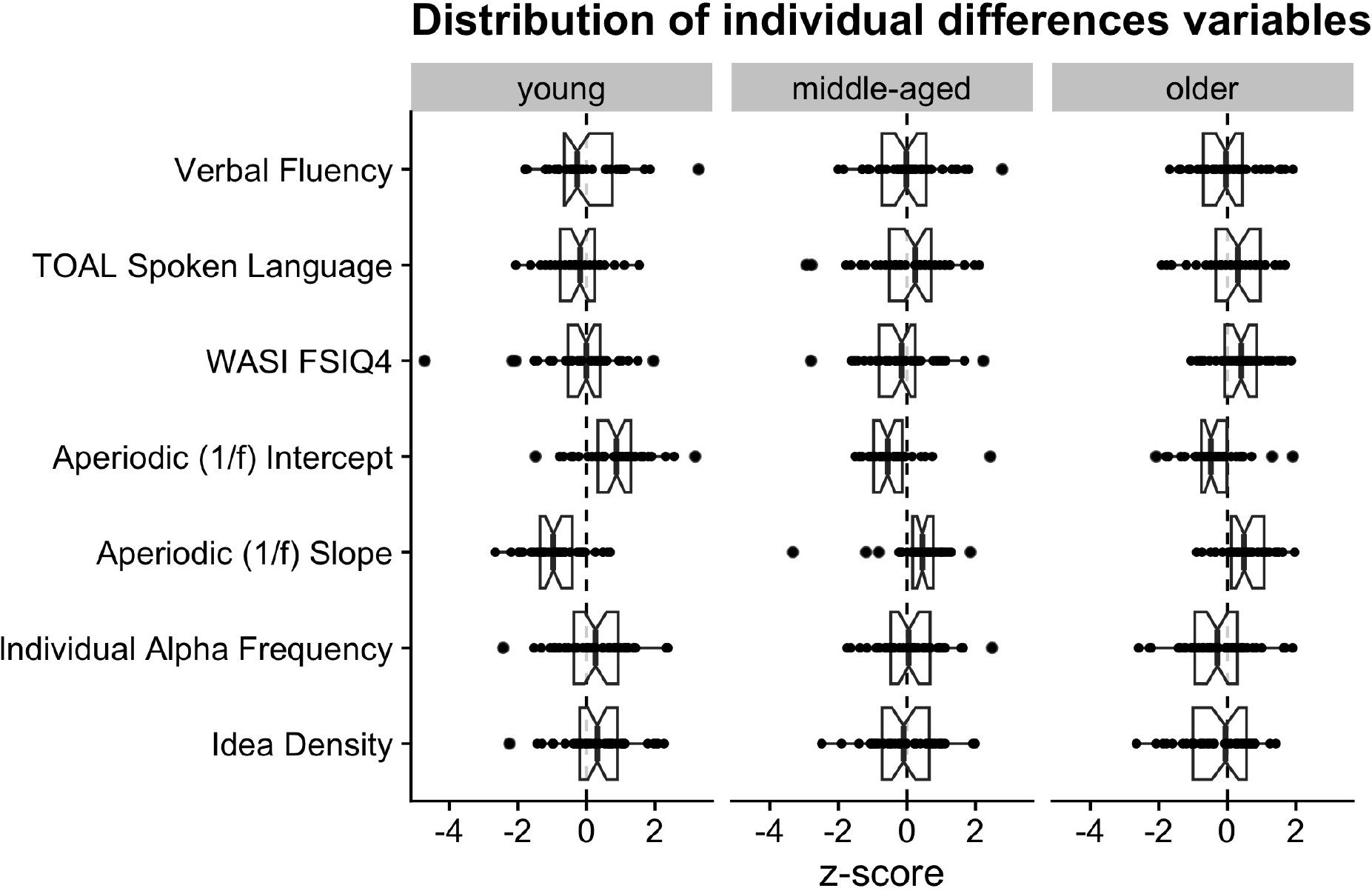
Distribution of the (z-transformed) resting-state EEG-based and behavioural individual differences metrics by age group.

Correlations between the individual differences measures collected as part of this experiment are visualised in Figure 4.

**Figure 4:**
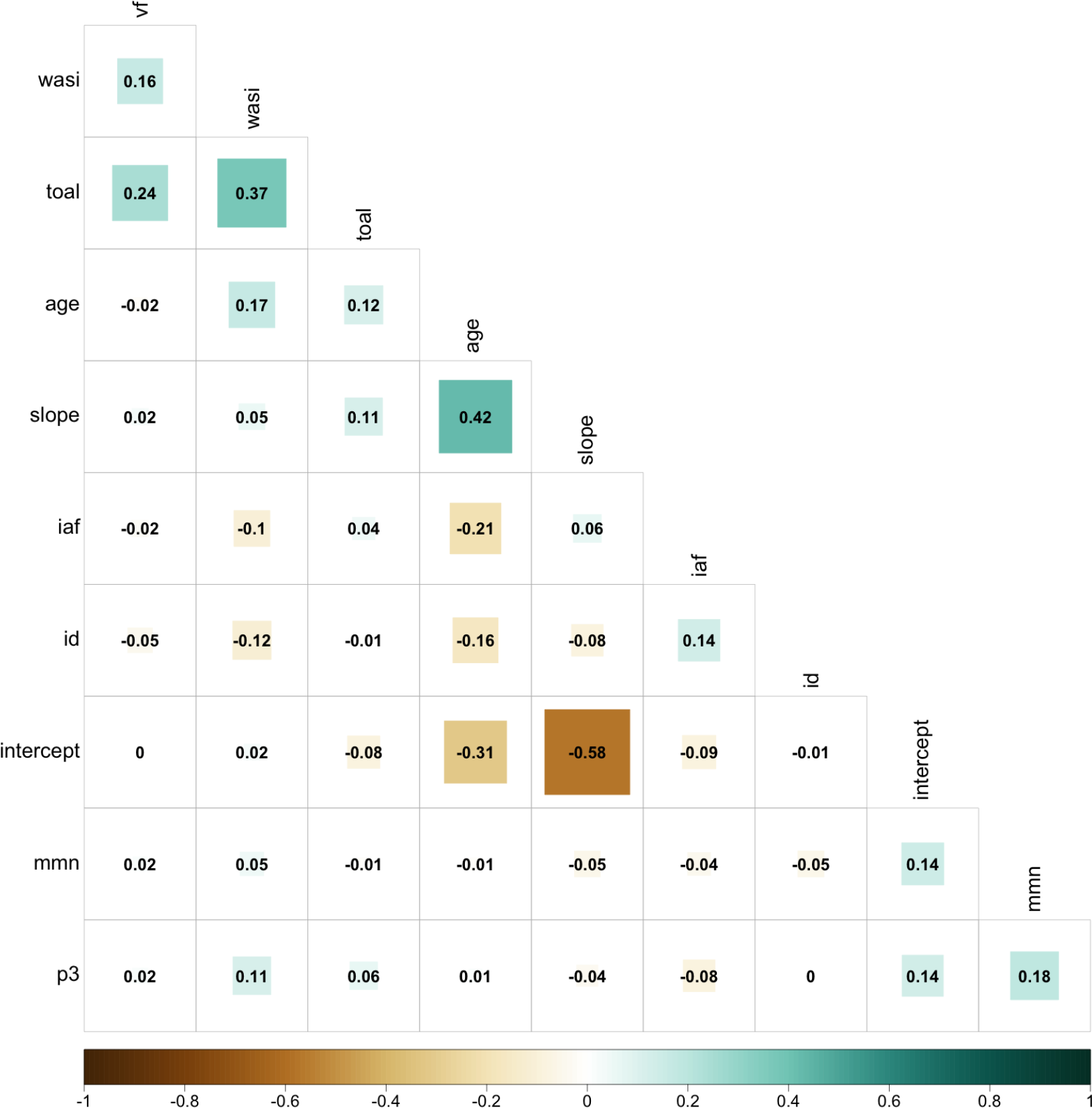
Visualisation of the correlation coefficients (Kendall’s *τ* ) between individual differences predictors for the present study.

As expected, age showed a correlation with a number of the metrics of interest. Replicating previous results, older age was associated with a downshifting of the aperiodic intercept, a shallower aperiodic slope (note that steeper slopes are denoted by more negative values, hence the negative correlation here), a lower individual alpha peak frequency (IAF) and lower Idea Density (ID). The relationship between age and the aperiodic intercept and slope parameters was more substantial than that with the other predictors. Note that aperiodic slope and intercept also showed a moderate correlation such that higher intercepts were associated with steeper slopes.

The three cognitive metrics, WASI FSIQ-4 as a measure of cognitive ability, TOAL-4 as a measure of spoken language proficiency, and verbal fluency, were positively correlated with one another. This is perhaps not surprising given that all involve tests of language abilities.

The two metrics derived from the auditory oddball paradigm, MMN and P3 amplitude, showed a small positive correlation with one another and with aperiodic intercept.

### 2.2 Main language processing task

In the following, we report the analysis of the main language processing task. Here, we examined the extent to which the amplitude of the N400 ERP attuned to speaker-based surprisal over the course of the experiment and how this attunement was modulated by age and individual differences. We begin by focusing on effects of age only, before examining whether individual differences metrics derived from neurophysiological and behavioural measures can explain variance in the N400 response over and above that already accounted for by age differences.

#### 2.2.1 More conservative model adaptation to novel language input in older adults

In addition to Age, Speaker-based Surprisal, Epoch (as a proxy for time on task), Canonicity, Word frequency and Prestimulus Amplitude (Alday, 2019), we included the follwoing control variables as fixed effects: Quadratic prestimulus amplitude, Quadratic speaker-based surprisal and Cubic speaker-based surprisal (see section 4.8.1 for details). The random effects structure was determined using a parsimonious model selection procedure, as described in section 4.8.1.

Multiple five-way interactions involving our three predictors of main interest, Speaker-based Surprisal, Epoch and Age, reached significance (see the model summary in the Appendix for details).

However, as we consider Prestimulus amplitude, Word frequency and Canonicity covariates of no interest for the present analysis, we focus on the interaction of Age x Speaker-based surprisal x Epoch (Estimate = 0.0484, Std. Error = 0.0115, *z* = 4.20, *p <* 0.0001), which is visualised in Figure 5

**Figure 5:**
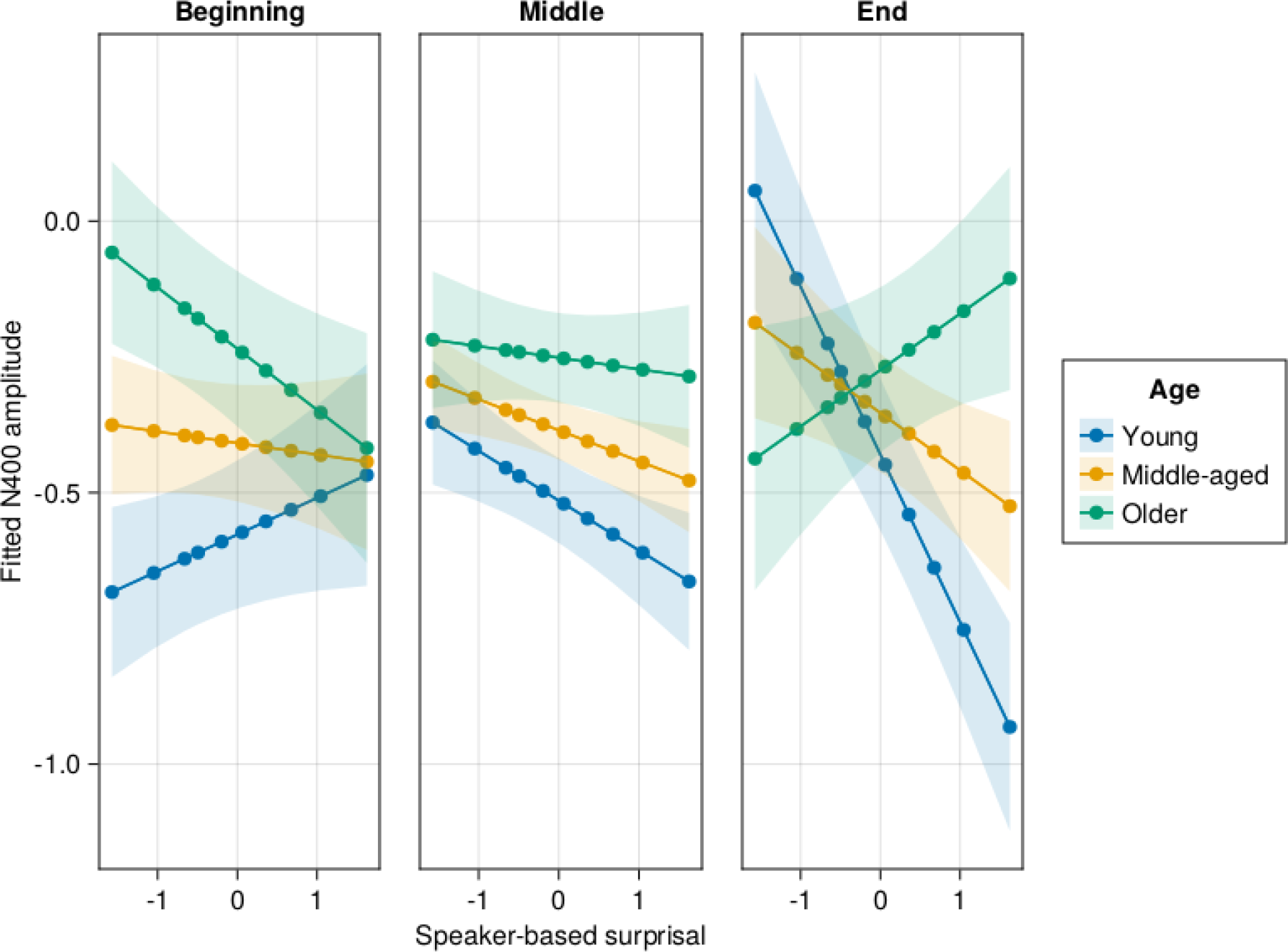
Effects of age on changes in the relationship between speaker-based surprisal and N400 amplitude over the course of the experiment. Note that position in the experiment (operationalised via epoch in the statistical model) is trichotomised into beginning (estimated for the first quantile), middle (median) and end (last quantile) for visualisation purposes only; epoch was included in the model as a continuous predictor. The same holds for age, which is trichotomised for visualisation purposes (estimated for the minimum, median and maximum ages in our sample) but was entered into the statistical model as a continuous predictor. Speaker-based surprisal is visualised from the 5th to the 95th quantile to remove extreme values. Shaded ribbons signify standard errors.

The figure indicates that younger adults show a strong attunement to speaker-based surprisal over the course of the experiment: note the strong relationship between N400 amplitude and speaker-based surprisal in the right-hand facet of the figure. This attunement is reduced in middle-aged adults and absent in older adults, who in fact even show a slight tendency towards an inverted effect at the end of the experiment. These results support our hypothesis that ageing is associated with a slowed rate of sensory learning, in this case through the novel characteristics of the speakers’ adjective order choices within the context of the experiment, and increased reliance on established top-down predictive models.

#### 2.2.2 Individual differences in predictive language model updating beyond those of age

On account of the relationship between age and most of the individual differences predictors of interest, the individual differences predictors were residualised on age prior to being included in any LMMs of interest. To this end, linear regressions were run with the individual differences predictor as the outcome variable and linear, quadratic and cubic effects of age included in successive models and compared to a base model including only an intercept term. Likeli-hood ratio tests were used in a forward model selection procedure to determine the best-fitting model. For a visualisation of the results, see Figure 12 in the Appendix.

Goodness-of-fit metrics for the individual differences models in comparison to the model without individual differences (including only age) are shown in Table 1. All individual differences measures improved model fit in terms of AIC over the model including age alone. The slope, intercept and ID models also showed a better fit to the data than the base model in terms of BIC, with the IAF and WASI models showing a very slight improvement in terms of BIC.

**Table 1:**
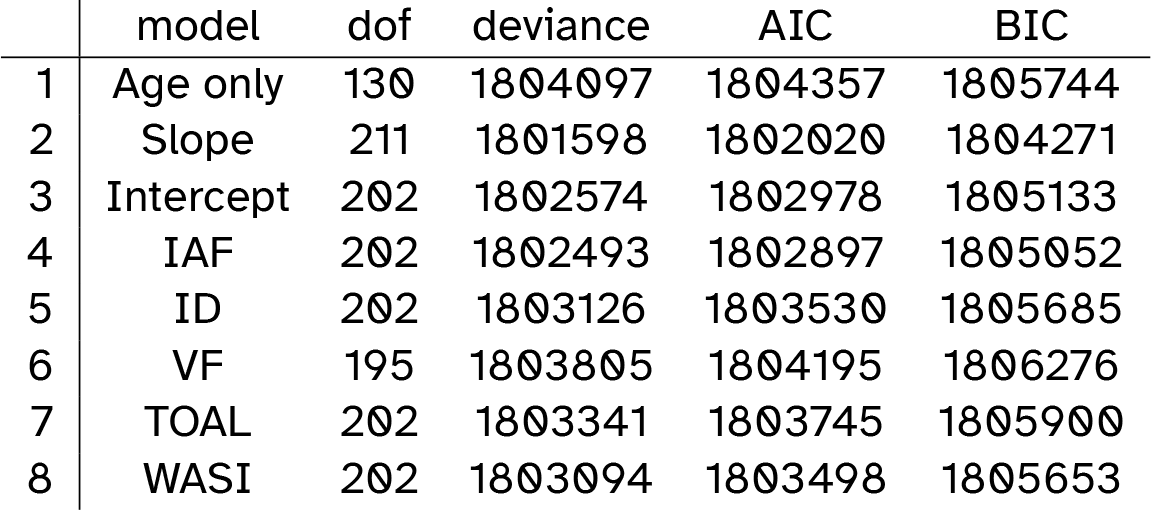
Comparison of goodness-of-fit metrics for the LMM involving only age and the individual differences models.

Full model summaries for all of the individual differences models are included in the Appendix.

##### Aperiodic (1/f) slope

For aperiodic slope, the highest-order significant interaction was that of Prestimulus Amplitude x Age x Frequency x Speaker-based Surprisal x Canonicity x Epoch x Slope (Estimate = -0.1372, Std. Error = 0.0558, *z* = -2.46, *p <* 0.0139). However, in view of the fact that, as noted above, Prestimulus Amplitude, Canonicity and frequency are covariates of no interest in the present analysis, the interaction of main interest was Age x Speaker-based Surprisal x Epoch x Slope (Estimate = -0.3335, Std. Error = 0.0746, *z* = –4.47, *p <* 0.0001). This interaction is visualised in Figure 6.

**Figure 6:**
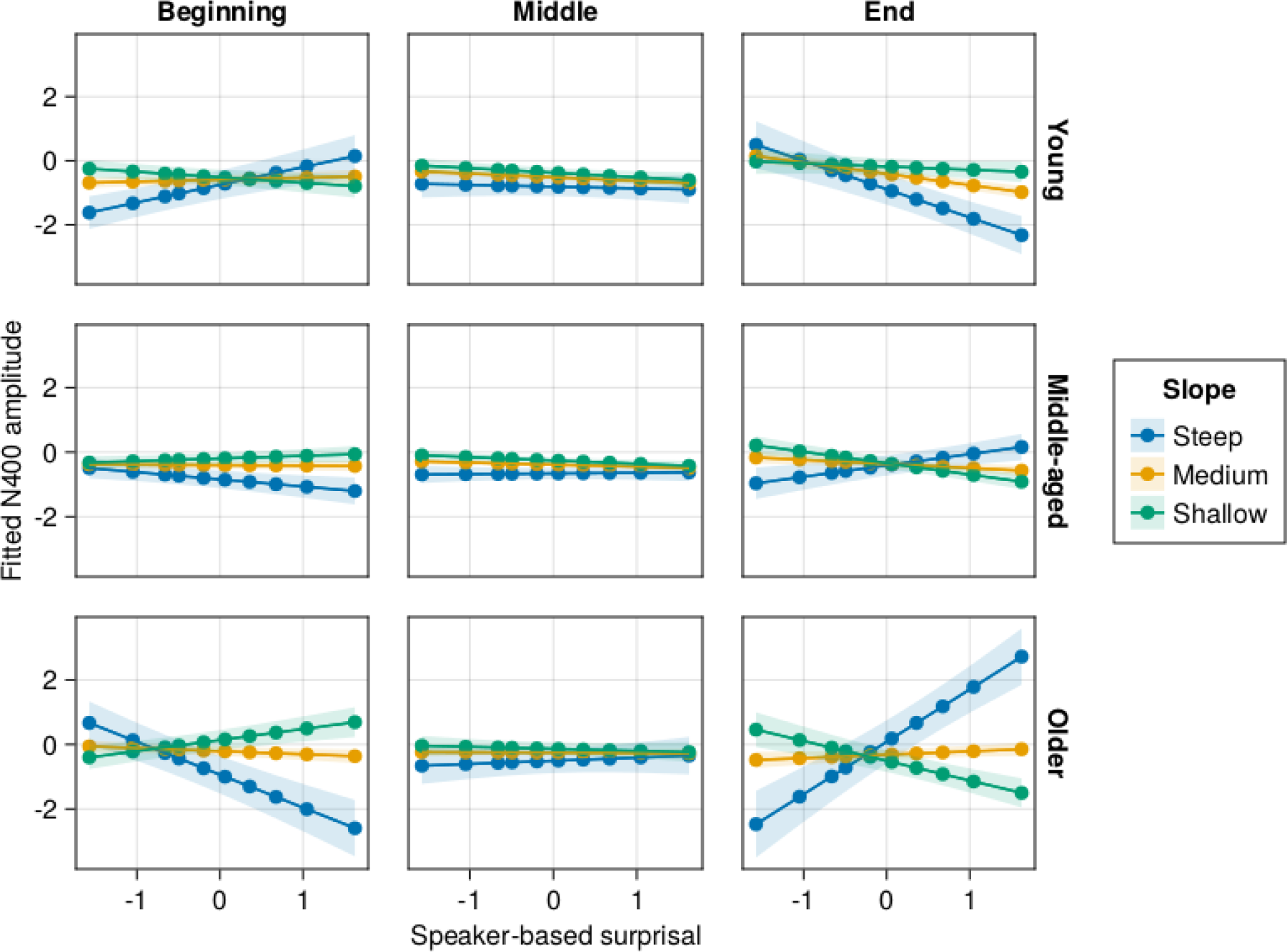
Effects of age and (age-residualised) aperiodic slope on changes in the relationship between speaker-based surprisal and N400 amplitude over the course of the experiment. Note that position in the experiment (operationalised via epoch in the statistical model) is trichotomised into beginning (estimated for the first quantile), middle (median) and end (last quantile) for visualisation purposes only; epoch was included in the model as a continuous predictor. The same holds for age, which is trichotomised for visualisation purposes (estimated for ages 18, 50 and 83 as the minimum, median and maximum of ages in our sample) but was entered into the statistical model as a continuous predictor. Speaker-based surprisal is visualised from the 5th to the 95th quantile to remove extreme values. Shaded ribbons signify standard errors.

##### Aperiodic Intercept

The analysis of the aperiodic intercept data revealed several 6-way interactions and a five-way interaction of Age x Speaker-based Surprisal x Canonicity x Epoch x Intercept (Estimate = -0.07977, Std. Error = 0.01667, *z* = -4.79, *p <* 0.0001). However, the main interaction not including the covariates of no interest did not reach significance: Age x Speaker-based Surprisal x Epoch x Intercept (Estimate = 0.01166, Std. Error = 0.0170, *z* = 0.69, *p* = 0.4930).

Effects are visualised in Figure 7.

**Figure 7:**
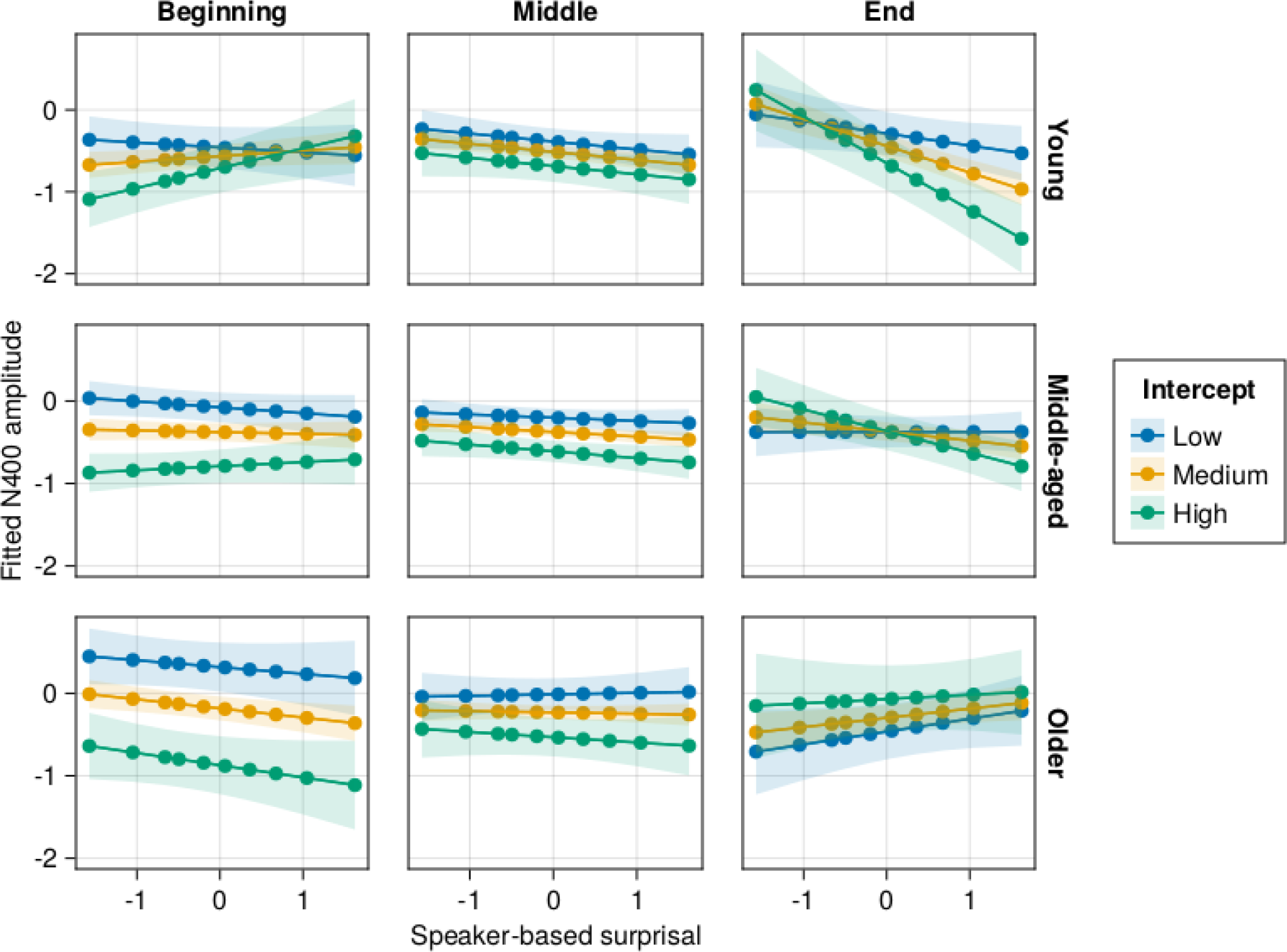
Effects of age and (age-residualised) aperiodic intercept on changes in the relationship between speaker-based surprisal and N400 amplitude over the course of the experiment. Note that position in the experiment (operationalised via epoch in the statistical model) is trichotomised into beginning (estimated for the first quantile), middle (median) and end (last quantile) for visualisation purposes only; epoch was included in the model as a continuous predictor. The same holds for age, which is trichotomised for visualisation purposes (estimated for ages 18, 50 and 83 as the minimum, median and maximum of ages in our sample) but was entered into the statistical model as a continuous predictor. Speaker-based surprisal is visualised from the 5th to the 95th quantile to remove extreme values. Shaded ribbons signify standard errors.

##### Individual Alpha Frequency (IAF)

As for the aperiodic intercept, the analysis of IAF revealed several 6-way interactions, but the main interaction of interest without the covariates of no interest did not reach significance: Age x Speaker-based Surprisal x Epoch x IAF (Estimate = -0.01080, Std. Error = 0.0179, *z* = -0.60, *p* = 0.5469). This is reflected in the visualisation in Figure 8.

**Figure 8:**
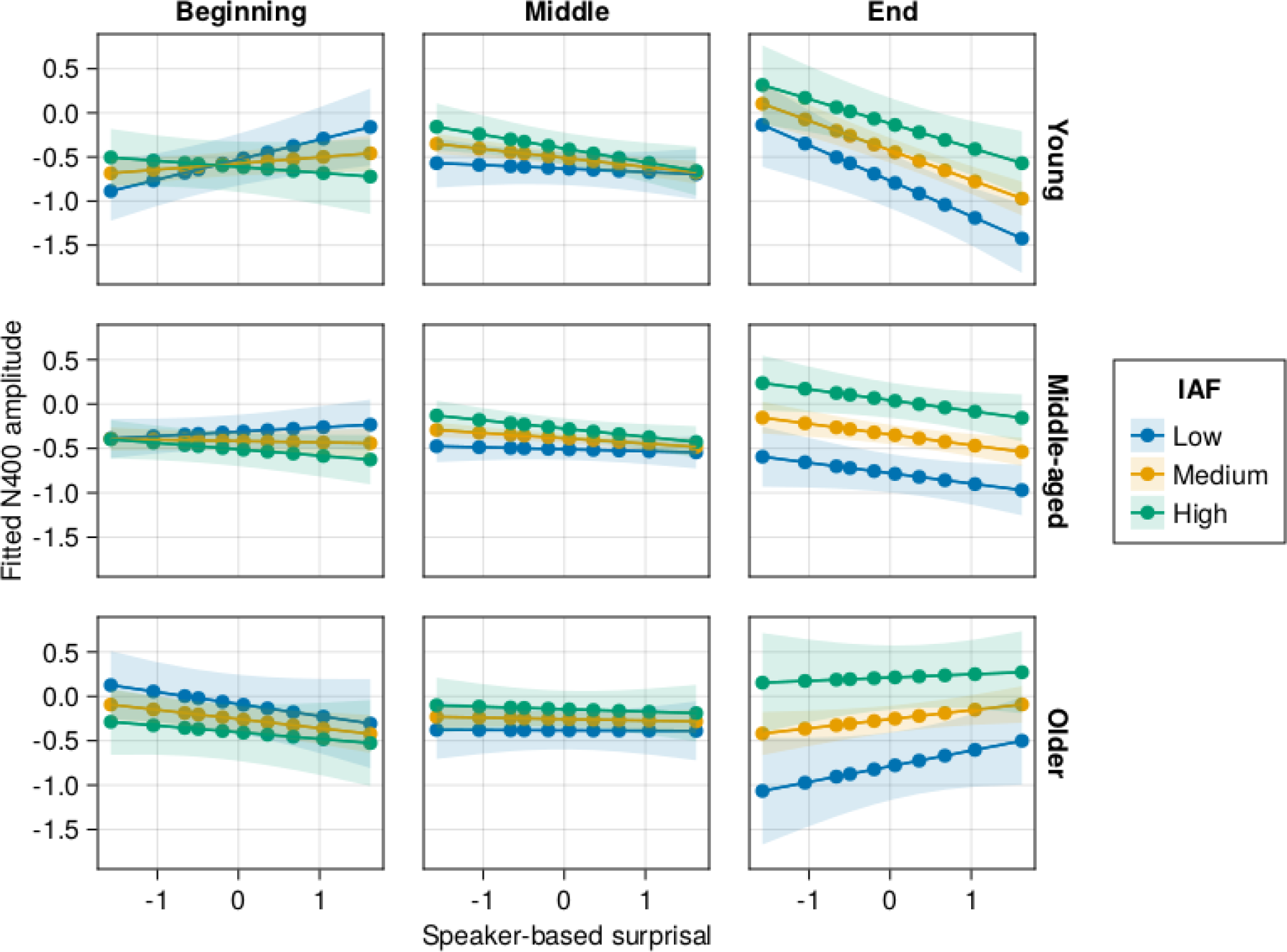
Effects of age and (age-residualised) individual alpha frequency (IAF) on changes in the relationship between speaker-based surprisal and N400 amplitude over the course of the experiment. Note that position in the experiment (operationalised via epoch in the statistical model) is trichotomised into beginning (estimated for the first quantile), middle (median) and end (last quantile) for visualisation purposes only; epoch was included in the model as a continuous predictor. The same holds for age, which is trichotomised for visualisation purposes (estimated for ages 18, 50 and 83 as the minimum, median and maximum of ages in our sample) but was entered into the statistical model as a continuous predictor. Speaker-based surprisal is visualised from the 5th to the 95th quantile to remove extreme values. Shaded ribbons signify standard errors.

##### Idea Density

For Idea Density, the analysis showed a significant top-level interaction of Prestimulus Amplitude x Age x Frequency x Speaker-based Surprisal x Epoch x Idea Density (Estimate = 1.4510, Std. Error = 0.3534, *z* = 4.11, *p <* 0.0001). The relevant lower-order interaction was, however, once again not significant: Age x Speaker-based Surprisal x Epoch x Idea Density (Estimate = 0.0142, Std. Error = 0.3445, *z* = 0.04, *p* = 0.9672). This is again reflected in Figure 9.

**Figure 9:**
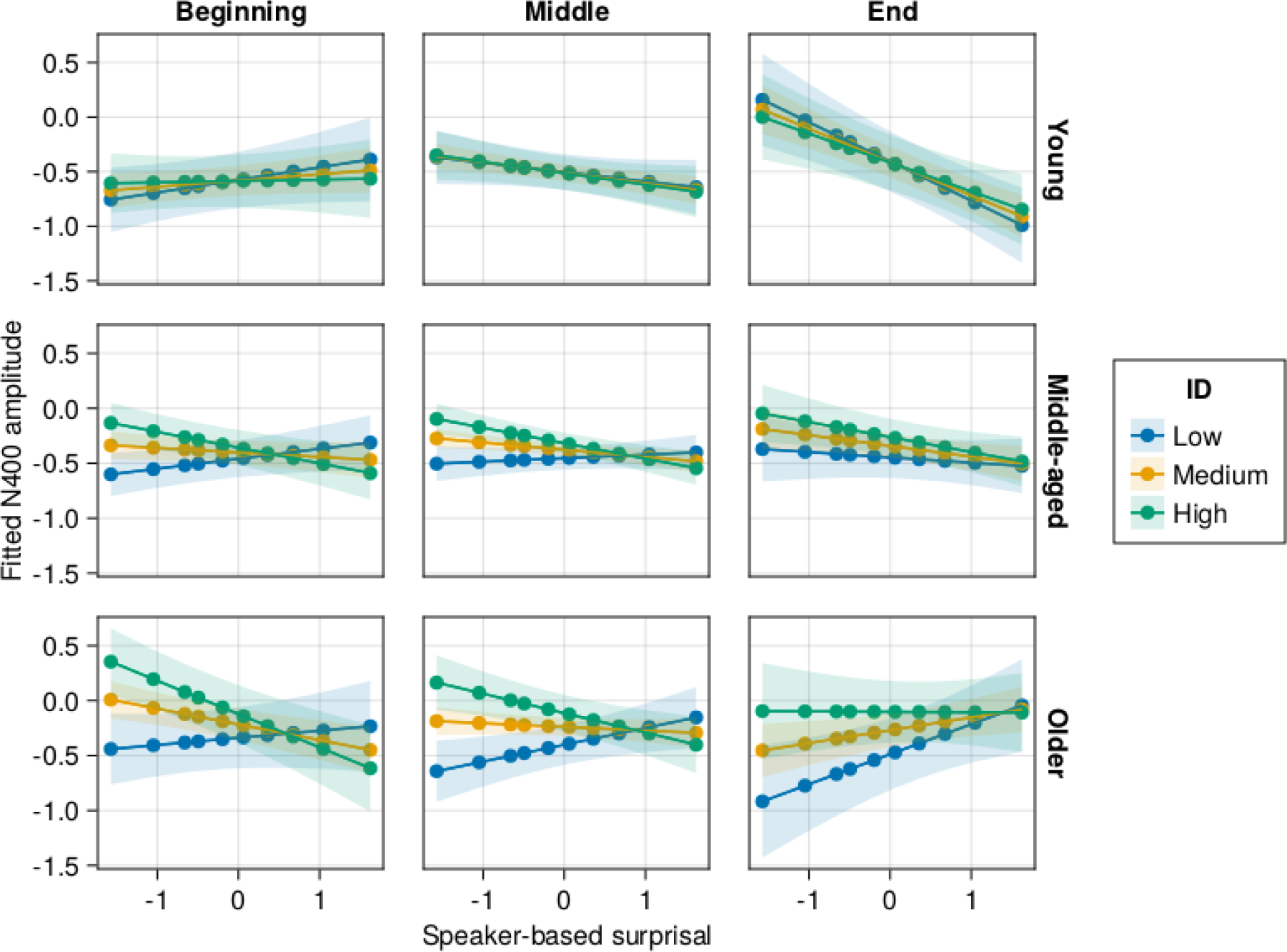
Effects of age and (age-residualised) Idea Density (ID) on changes in the relationship between speaker-based surprisal and N400 amplitude over the course of the experiment. Note that position in the experiment (operationalised via epoch in the statistical model) is trichotomised into beginning (estimated for the 5th quantile), middle (50th quantile) and end (95th quantile) for visualisation purposes only; epoch was included in the model as a continuous predictor. The same holds for age, which is trichotomised for visualisation purposes (estimated for ages 18, 50 and 83 as the minimum, median and maximum of ages in our sample) but was entered into the statistical model as a continuous predictor. Shaded ribbons signify standard errors.

##### Cognitive Measures

As is apparent from Table 1, the model including verbal fluency showed only very little improvement over the model only including age. AIC values for the TOAL and WASI models also indicated a worse fit for these models in comparison to the other individual differences predictors, with the exception of the IAF model, which showed similar AIC and BIC values to the WASI model.

Analyses involving verbal fluency, TOAL and WASI scores as predictors did not yield very compelling results regarding individual differences beyond those of age. These analyses are thus reported in the Appendix (cf. Figures 13 – 15), which show extremely broad confidence intervals and do not allow for any conclusions to be drawn for the influence of these individual differences predictors beyond the effect of age alone.

##### MMN and P3 estimates from the auditory oddball task

By-participant fitted MMN and P3 values from the auditory oddball, again residualised against age, were used as two final individual differences predictors for the N400 analysis. These models needed to be fit to a slightly reduced sample of participants (n=114) due to missing values. Hence, to allow for a comparison with a base model only including age and no other individual-differences measures, we refit the best-fitting model without individual differences from the previous analysis to this reduced data set and verified that the random effects structure was motivated by the data. We then proceeded to add the residualised, fitted MMN and P3 values (in separate models) as for the other individual differences predictors above. Goodness-of-fit metrics for the model comparison are shown in Table 2. As is apparent from the table, the MMN and P3 models both improve model fit over and above the model only including age in regard to both AIC and BIC.

**Table 2:** Comparison of goodness-of-fit metrics for the LMM involving only age and the individual differences models involving MMN and P3 estimates derived from the auditory oddball task.

|  | model | dof | deviance | AIC | BIC |
| --- | --- | --- | --- | --- | --- |
| 1 | Age only | 130 | 1756128 | 1756388 | 1757771 |
| 2 | MMN | 202 | 1754502 | 1754906 | 1757055 |
| 3 | P3 | 202 | 1754346 | 1754750 | 1756900 |

For both the MMN and the P3, the analysis revealed several five-way interactions and the pertinent interaction without any covariates of no interest: Age x Speaker-based Surprisal x Epoch x MMN amplitude (Estimate = 0.0566, Std. Error = 0.01698, *z* = 3.33, *p* = 0.0009); Age x Speaker-based Surprisal x Epoch x P3 amplitude (Estimate = 0.0518, Std. Error = 0.0163, *z* = 3.18, *p* = 0.0015). These effects are visualised in Figures 10 and 11 for the MMN and the P3, respectively.

**Figure 10:**
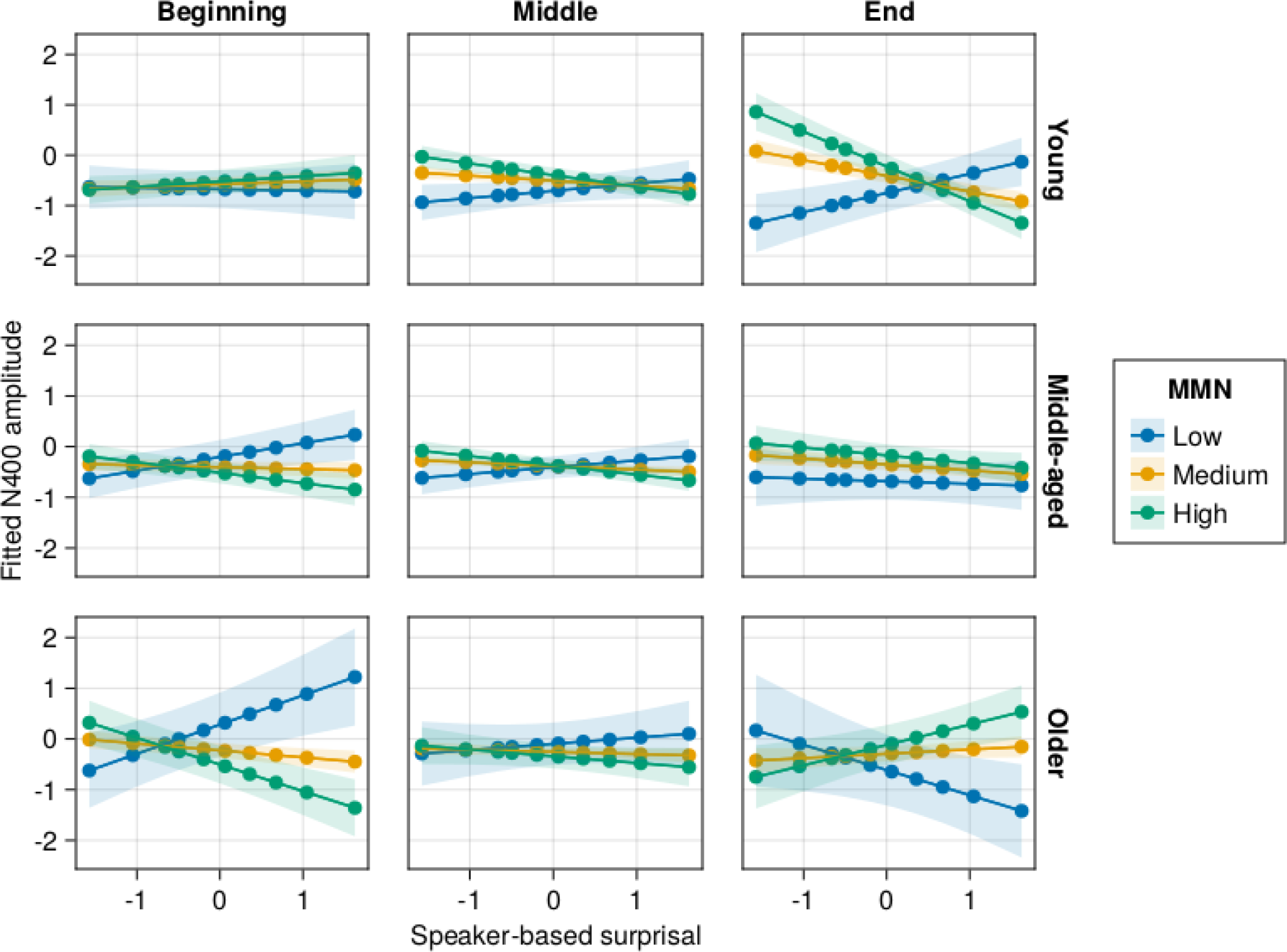
Effects of age and (age-residualised) fitted MMN amplitude (as derived from the auditory oddball task) on changes in the relationship between speaker-based surprisal and N400 amplitude over the course of the experiment. Note that position in the experiment (operationalised via epoch in the statistical model) is trichotomised into beginning (estimated for the 5th quantile), middle (50th quantile) and end (95th quantile) for visualisation purposes only; epoch was included in the model as a continuous predictor. The same holds for age, which is trichotomised for visualisation purposes (estimated for ages 18, 50 and 83 as the minimum, median and maximum of ages in our sample) but was entered into the statistical model as a continuous predictor. Shaded ribbons signify standard errors.

**Figure 11:**
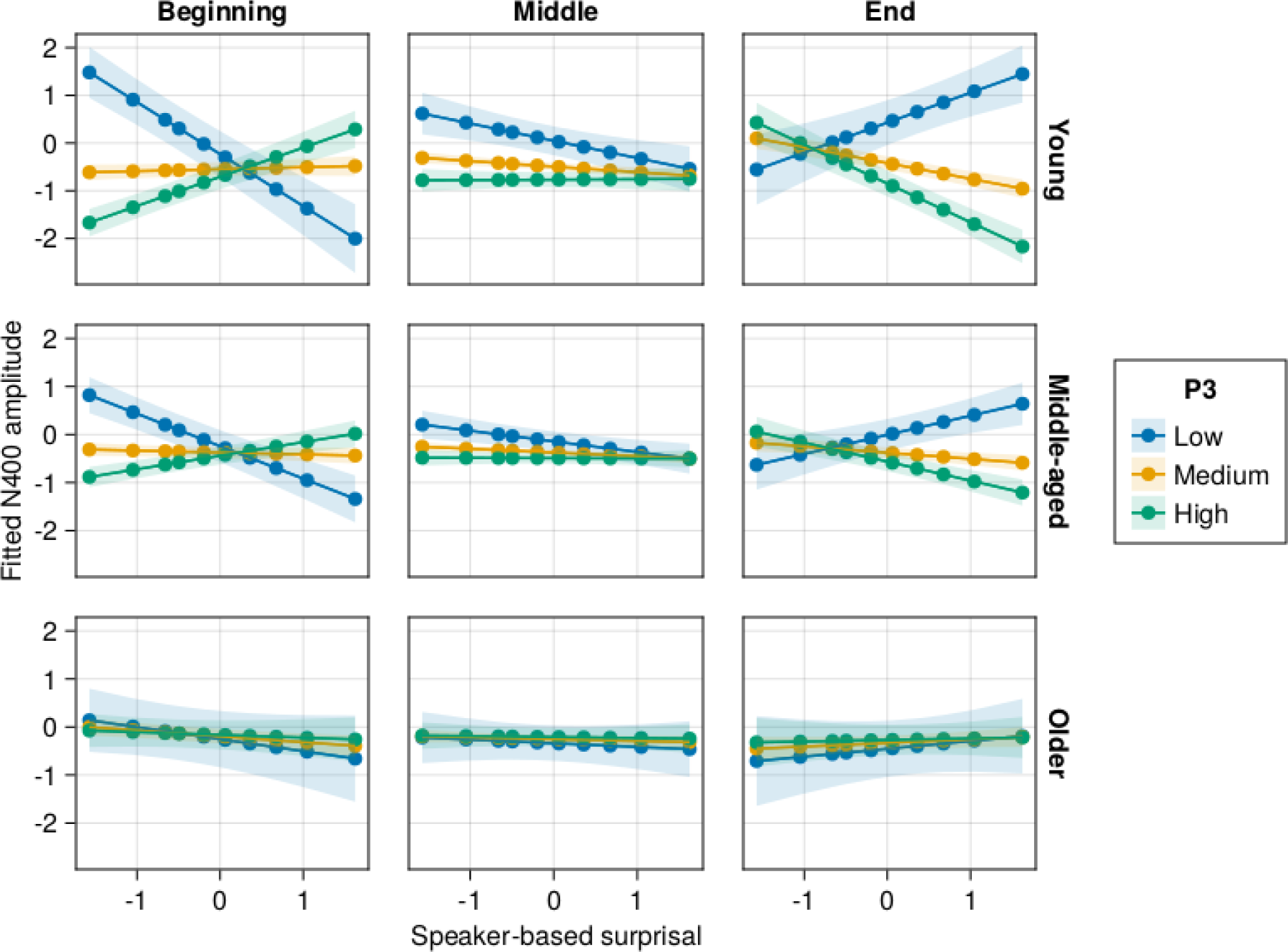
Effects of age and (age-residualised) fitted P3 amplitude (as derived from the auditory oddball task) on changes in the relationship between speaker-based surprisal and N400 amplitude over the course of the experiment. Note that position in the experiment (operationalised via epoch in the statistical model) is trichotomised into beginning (estimated for the 5th quantile), middle (50th quantile) and end (95th quantile) for visualisation purposes only; epoch was included in the model as a continuous predictor. The same holds for age, which is trichotomised for visualisation purposes (estimated for ages 18, 50 and 83 as the minimum, median and maximum of ages in our sample) but was entered into the statistical model as a continuous predictor. Shaded ribbons signify standard errors.

## 3 Discussion

The present study examined lifespan changes in predictive model adaptation and the way in which these are modulated by individual differences. Participants performed a naturalistic auditory language comprehension task, in which we measured the adaptation of individual brain responses to speakers’ word order idiosyncrasies using the trial-by-trial-based measure of speaker-based surprisal at the level of adjective clusters (cf. Bornkessel-Schlesewsky et al., 2022). As hypothesised, the N400 ERP response’s degree of attunement to speaker-based surprisal was reduced with increasing age: young adults showed a strong alignment between N400 amplitude and speaker-based surprisal by the end of the experimental session; this was reduced for middle-aged adults and absent for older adults. All individual differences measures explained additional variance in the N400 over and above that explained by age, suggesting that individual differences in information processing modulate the effects of ageing. In the following sections, we first discuss the overall effects of age, before turning to an interpretation of the individual differences.

### 3.1 Stronger reliance on top-down predictive models and reduced sensory learning in older adults

The observation that N400 amplitude attunement to speaker-based surprisal is reduced with increasing age supports the hypothesis that ageing leads to a stronger reliance on established predictive models and, accordingly, a lower tendency to update these models based on unexpected sensory input. While this has previously been reported for basic auditory processing (Moran et al., 2014), multisensory illusions (Chan et al., 2021) and sensorimotor attenuation (Wolpe et al., 2016), the present study is the first to demonstrate that it also holds in the domain of higher-order language processing.

As argued by Moran et al. (2014), the stronger weighting of top-down model predictions in older adults has the functional benefit of reducing model complexity and avoiding overfitting. This is considered to reflect an optimisation of information processing over the adult lifespan. We suggest that the lower propensity for model updating may be promoted by several age-related neurophysiological changes. Older adults show higher levels of neural noise, as reflected in shallower aperiodic slopes (Donoghue et al., 2020; Merkin et al., 2023; Voytek et al., 2015; Waschke et al., 2017), and an age-related reduction of neural gain (Li et al., 2000; Li et al., 2001; Li & Rieckmann, 2014; Li et al., 2006). Pertermann et al. (2019) recently provided empirical evidence for a possible link between aperiodic activity and neural gain control. In Bornkessel-Schlesewsky et al. (2022), we suggested that lower neural noise and increased neural gain may allow for a more flexible deployment of precision-weighting mechanisms in predictive model updating. Within such conditions, information that is more relevant in a given situation can be distinguished more readily from irrelevant information, possibly leading to an increased allocation of attention to high-precision sensory evidence (Parr & Friston, 2017). Age-related increases in neural noise and concomitant reductions in neural gain reduce this ability for flexible model adaptation, leading to decreased sensory learning and a stronger reliance on established predictive (language) models (Moran et al., 2014).

Further support for this perspective comes from the passive auditory oddball task that our participants completed in addition to the main language processing task. In contrast to younger and middle-aged adults, who show a P3 effect for oddball versus standard tones across the entire task, the P3 effect for older adults develops across the duration of the session. Older adults thus require more time on task to attribute a motivationally significant status to the oddball tone, as reflected by increased P3 amplitude (Nieuwenhuis et al., 2005). This is compatible with the view that older adults are less readily able to distinguish between relevant and irrelevant stimuli.^1^ Importantly, this result does not appear to be related to a reduced discriminability of standard versus oddball tones for the older adults, as they show an MMN effect for oddball versus standard tones throughout the experiment.^2^

### 3.2 Individual differences in predictive model adaptation across the lifespan are most pronounced for aperiodic slope and intercept

While all individual differences metrics improved LMM fit over and above the inclusion of age alone, the aperiodic metrics slope and intercept showed the most pronounced individual differences in relation to predictive model updating. Young adults with a steeper aperiodic slope and those with a higher aperiodic intercept showed stronger model adaptation to speaker-based surprisal in comparison to individuals with flatter slopes and lower intercepts (see Figures 6 and 7). While middle-aged adults with higher aperiodic intercepts still showed a slight tendency towards adaptation, the analysis including intercept revealed no adaptation effects for older adults. For aperiodic slope, there were no apparent effects for middle-aged adults, but a “reversed” effect for older adults, with those with a steeper slope showing an anti-adaptation effect of more pronounced N400 amplitudes for low vs. high speaker-based surprisal. We will return to this unexpected effect below.

In contrast to aperiodic slope and intercept, Individual Alpha Frequency (IAF) and the models based on behavioural metrics did not shed any additional light on age-related changes in model adaptation. IAF did not modulate the likelihood for adaptation, with low IAF individuals showing generally more negative N400 amplitudes across all levels of speaker-based surprisal and across all ages. In the Idea Density (ID) analysis, young adults showed a general adaptation effect regardless of ID, while there was no discernible adaptation effect for the middle-aged or older adults: confidence intervals for the effects at the end of the experiment were so broad that we cannot distinguish between ID-related groups and there appears to be a generally high degree of variability (cf. Figure 9). The behavioural metrics based on cognitive ability (WASI FSIQ-4), verbal fluency (VF) and spoken language proficiency (TOAL-4) did not yield any compelling results (see Figures 13–15 in the Appendix).

The overall pattern of results observed in the present study is thus driven most strongly by individual differences in aperiodic slope and intercept. Interestingly, these were the two measures that correlated most strongly with age (see Figures 3 and 4). It is striking, however, that individual differences in these measures not captured by age showed an accentuation of the pattern beyond the effects explained by age alone. As discussed above, we assume that the lower levels of neural noise and increased neural gain associated with steeper aperiodic slopes facilitate the ability to flexibly discriminate between relevant and irrelevant sensory information from the perspective of model updating (Bornkessel-Schlesewsky et al., 2022). This enables more rapid learning, possibly mediated by precision-related shift in attention (Parr & Friston, 2017), and a concomitant increase in the ability to adapt predictive models to the current sensory environment.

Less is known about the role of the aperiodic intercept in ageing, beyond the basic observation 23 that the age-related downshifting of the intercept likely reflects reduced aggregate population spiking (Donoghue et al., 2020; Voytek & Knight, 2015). This renders any interpretation of our intercept-related results somewhat more speculative. Recall that there is a moderate correlation between intercept and slope (*τ* = *−*0.58) in the present sample, with higher intercepts associated with steeper slopes. Thus, to a certain extent, the pattern observed here for aperiodic intercept may simply reflect this relationship. Note, however, that the pattern of results was not entirely mirrored between the two metrics, thus suggesting at least partially differentiable mechanisms.

We suggest that the results observed for the MMN and P3 when included as individual differences predictors within the N400 models could help to illuminate this putative functional dissociation between aperiodic slope and intercept. The pattern for the MMN (Figure 10) closely mirrored that observed for the slope: larger individual MMN effects were associated with stronger model adaptation to speaker-based surprisal for younger adults, while there was an inverse (“anti-adaptation”) effect for older adults with more pronounced MMN amplitudes. In contrast, the pattern for the P3 (Figure 11) more closely mirrored that for the intercept, with larger P3 effects associated with stronger model adaptation in young adults in particular. The MMN is viewed as an index of sensory learning in predictive coding accounts (Garrido et al., 2009; Moran et al., 2014), with the MMN effect (the amplitude difference between oddball and standard stimuli) thought to reflect the increasing precision of internal model predictions about the standard stimulus (Friston, 2005; Todd et al., 2014; Todd et al., 2011; Todd et al., 2013). The P3, by contrast, reflects the processing of motivationally significant stimuli and is associated with increased neural gain through the release of noradrenaline from the locus coeruleus (Nieuwenhuis et al., 2005). In the literature on the role of precision in predictive model updating, P3 effects have been linked to prediction errors on the probabilistic context in which stimuli appear – which defines stimulus precision – as opposed to prediction errors about the (content of) the stimulus itself (Feldman & Friston, 2010). This distinction between inferences about the content of a sensory signal and inferences about its precision has been described as akin to the dissociation between first-order statistical inferences (estimating the mean) and secondorder statistical inferences (estimating the variance) (Hohwy, 2012). Accordingly, MMN effects in the present study can be viewed as reflecting precision-weighted prediction errors about each tone, while P3 effects reflect prediction errors about the precision itself (cf. Schröger et al., 2015). Both aspects ultimately contribute to learning and the likelihood for model updating.

Building on these insights, the current results provide initial evidence to suggest that aperiodic slope / neural noise may be more strongly associated with individual differences in first-order inferences (content-based prediction errors), while aperiodic intercept may more closely reflect individual differences in second-order inferences (context-based prediction errors).

This account may also offer a possible explanation for the inverse (“anti-adaptation”) N400 effect observed for older adults with a steeper aperiodic slope. This effect was not expected and we can only speculate as to its interpretation. Assuming that individuals with steeper aperiodic slopes show an amplification of precision-weighted prediction error signals for sensory input, it may be the case that, rather than leading to more rapid model adaptation as in young adults, the competing tendency imposed by a strong, established predictive model in older adulthood leads to a change of processing strategy for these individuals. Recognising the uncertainty of the sensory input presented to them in the current experimental context, these individuals may have eventually refrained from updating their models in response to surprising input. As noted above, however, this explanation is a mere speculation at this point and will require further examination in future research.

### 3.3 Ageing effects on predictive language model updating

The present study is by no means the first to examine higher-order language comprehension in older adults. Intriguingly, a common conclusion from the relatively substantial existing literature on ageing and the electrophysiology of sentence comprehension has been that older adults do not draw on contextual information to the same extent as younger adults and, accordingly, that they rely less on predictive language processing than younger adults (e.g. DeLong et al., 2012; Federmeier & Kutas, 2005; Wlotko & Federmeier, 2012). For example, Wlotko and Federmeier (2012) parametrically varied word predictability inside sentence contexts presented to young and older adults, with predictability operationalised via a manipulation of cloze probability from 0 to 100. In contrast to the graded N400 response observed in young adults as a function of the degree of constraint / predictability, they argued that older adults show more of a binary response: they either derive some facilitation from the context for a predictable word (though this is reduced in magnitude in comparison to younger adults) or not. Likewise, DeLong et al. (2012) observed an N400 effect for unexpected versus expected articles in sentences such as “The day was breezy, so the boy went outside to fly an/a …” only for young but not for older adults.

This design has been argued to reflect prediction in sentence comprehension to a greater degree than typical N400 paradigms, as the (un-)expectedness of the article does not stem from its own meaning, which is the same for “a” and “an”, and rather depends on the following noun (expected in this example: “kite”). Hence, an effect at the position of the article was assumed to require a prediction-based rather than an integration-based interpretation (DeLong et al., 2005); however, see Nieuwland et al. (2018), who failed to observe this effect in a largescale replication study of this result for young adults as originally reported by DeLong et al. (2005).

These results initially seem incompatible with the current study in that they have collectively been taken to suggest a reduced reliance on predictive language processing by older adults. This is particularly interesting in view of the fact that, like the present results, the broader predictive coding literature outside of language suggests a higher reliance on top-down predictive models with increasing age rather than a shift away from predictive processing (Chan et al., 2021; Moran et al., 2014; Wolpe et al., 2016). Note, however, that the present design differs from existing ageing studies in the language literature in that prediction errors allow for learning across the experimental session based on the novel adjective order patterns produced by each speaker. By contrast, typical studies examining the interplay between ageing and sentence-internal constraint or predictability do not contain any information that allows for learning to occur across trials. Thus, an alternative interpretation of these existing results is that, in experimental settings involving a relatively large number of unexpected continuations (as is typical in traditional psycholinguistic experiments), older adults are more sensitive to the lower precision of the sensory input and thus tend to update their models less strongly than younger adults. Likewise, in the current experiment, older adults did not use the prediction errors elicited by unexpected adjective orders to learn / update their existing predictive models as strongly as younger and even middle-aged adults in spite of the fact that learning would have been possible across the course of the experiment here. Overall, we suggest that the data are compatible with the notion that older adults show a lower precision-weighting of prediction errors during language comprehension, with a concomitant reduction of internal model updating.

### 3.4 Summary and conclusions

By means of a moderately large lifespan sample (n=120; age-range: 18-83 years), the present study was the first to demonstrate that older adults show a stronger reliance on top-down predictive models and reduced learning based on sensory information during higher-order language comprehension. Similar findings have previously been reported for perceptual and sensorimotor processing. For older adults, our results showed a less pronounced adaptation of the N400 event-related potential to a trial-by-trial measure of adjective order expectedness within the experimental context given the idiosyncrasies of a particular speaker (“speaker-based surprisal”). This effect was graded across the adult lifespan, with younger adults showing the most pronounced degree of N400 amplitude adaptation, middle-aged adults showing a modest degree of adaptation and older adults, as a group, showing very little apparent adaptation.

Age-based differences were additionally modulated by individual differences measures, with resting-state derived aperiodic (1/f) slope and intercept metrics appearing to most clearly modulate individual model adaptability. Young participants with a steeper aperiodic slope or a higher aperiodic intercept showed the strongest model adaptation within their age group, with middle-aged adults with a high intercept also showing a more pronounced tendency towards model adaptation than those with a lower intercept. The patterns for aperiodic slope and intercept were mirrored by those using individual MMN and P3 amplitude estimates derived from a passive auditory oddball task as predictors, respectively. Through the association with the MMN and P3, we suggest that age-related changes in aperiodic slope / neural noise may be associated with changes in the magnitude of precision-weighted prediction errors signals to sensory input (first-order inferences), while changes in aperiodic intercept / aggregate population spiking may relate to the estimation of precision itself (second-order inferences) within the current environment. Together, these two mechanisms contribute to sensory learning by determining the precision-weighting of prediction errors, the main mechanism within the predictive coding framework that determines how an internal model will be updated in the face of unexpected sensory input. Our findings suggest that it may be fruitful to examine whether these two purported mechanisms are indeed separable, associated with distinct individual differences in information processing, and potentially contribute jointly but dissociably to the higher reliance on top-down predictive model information by older adults.

## 4 Materials and Methods

### 4.1 Participants

One hundred and twenty adults (88 identifying as female, 31 identifying as male, 1 identifying as other; mean age: 47.7 years, sd: 19.9, range: 18–83) participated in the experiment. In order to ensure approximately equal sampling across the adult lifespan, 40 were young adults (under 40 years of age), 40 were middle aged (between 40 and 60 years of age) and 40 were older adults (over 60 years of age). One hundred and nineteen participants were right-handed as assessed by the Edinburgh handedness inventory (Oldfield, 1971) and 1 was left-handed. All participants were native speakers of English who had not learned another language prior to starting school. They reported having no diagnosis of neurological or psychiatric conditions, normal hearing and normal or corrected-to-normal vision. The experimental protocol was approved by the University of South Australia’s Human Research Ethics Committee (protocol number 36348).

Three participants were excluded from the analysis of the main experimental task due to missing values in one of the key individual differences predictors.

The data for the young adults (main language processing task; individual differences measures of aperiodic slope, individual alpha frequency and idea density) were originally reported in Bornkessel-Schlesewsky et al. (2022, Experiment 2) and are reanalysed here for the purposes of the lifespan comparison.

### 4.2 Materials

The materials were identical to those in Bornkessel-Schlesewsky et al. (2022, Experiment 2). Participants were presented with 150 short passages (approx. 5 sentences in length), each of which contained two critical two-adjective noun phrases (NPs; e.g. “the huge grey elephant”). An example passage is provided below:

*Example of the passages presented to participants in the current study:*

Florence was enjoying her long-awaited holiday in Singapore with her close friends. One of the activities she was most looking forward to was visiting the zoo, where she had the opportunity to ride a **huge grey elephant**. Although standing in the **warm humid air** was dreadful, being waved to through the enclosure by the zookeeper brought a smile to her face.

The position of the critical NPs within each passage varied so as to not be predictable. The order of the prenominal adjectives either adhered to the expected sequence of “value *>* size *>* dimension *>* various physical properties *>* colour” (Kemmerer et al., 2007, p.240) or did not. Passages were recorded by two male speakers of Australian English with the probability of adjectives in the critical NPs occurring in an expected or unexpected order manipulated across speakers (*∼* 70%:30% vs. *∼* 30%:70%). The assignment of speakers to producing more expected or unexpected orders was counterbalanced across participants. To further accentuate the speaker-specific characteristics, presentation of the two speakers was alternated in a block-based manner: one block of the canonical speaker was followed by two blocks of the noncanonical speaker and two further blocks of the canonical speaker.

Comprehension questions were presented after approximately 1/3 of all passages (randomly distributed) to ensure attentive processing. An example comprehension question for the above example is: “Did the zookeeper wave at Florence?” (correct answer = yes).

### 4.3 Language Models

We employed the novel measure of speaker-based surprisal, as first introduced in Bornkessel-Schlesewsky et al. (2022), to examine how individuals differ in the adaptation of their predictive models to the current environment during language processing. Speaker-based surprisal was calculated as bigram-based surprisal for the second adjective (ADJ2) in the critical 2-adjective NPs embedded in the passages. To track predictability at the level of adjective within the experimental environment, we established adjective clusters using pre-derived word vectors from van Paridon and Thompson (2021) to determine similarities between adjectives, followed by a PCA-based dimensionality reduction and k-means clustering. For the 6 adjective clusters determined in this manner we calculated the NP-by-NP cumulative intra-experimental frequencies for the ADJ1-ADJ2 bigram cluster and the ADJ1 unigram cluster and then computed surprisal according as follows:

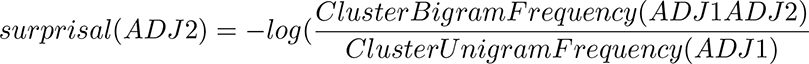

*surprisal*(*ADJ* 2) = *−log*(*ClusterBigramF requency*(*ADJ* 1*ADJ* 2) *ClusterUnigramF requency*(*ADJ* 1)

Here, *ClusterBigramFrequency(ADJ1ADJ2)* refers to the frequency with which two-adjective bigrams comprising a first adjective belonging to the cluster of ADJ1 and a second adjective belonging to the cluster of ADJ2 occured in the experiment, while *ClusterUnigramFrequency(ADJ1)* refers to the frequency with which adjectives belonging to the cluster of ADJ1 occurred in the experiment. To track surprisal fluctuations on an incremental, trial-by-trial basis, we calculated the NP-by-NP cumulative intra-experimental frequencies for the ADJ1-ADJ2 bigram cluster and the ADJ1 unigram cluster and then computed surprisal as described above. This was done separately for each speaker, thus allowing us to examine to what extent participants’ expectations adapted to the distributional properties of each of the two speakers within the experiment. Using speaker-based surprisal, we aimed to examine how participants’ N400 responses– as an assumed proxy for precision-weighted prediction error signals – were modulated by the exposure to adjective order variations throughout the course of the experiment and by each speaker.

For further details on the computation of speaker-based surprisal and examples of the adjective clusters derived using the procedure outlined above, see Bornkessel-Schlesewsky et al. (2022).

### 4.4 Behavioural individual differences measures

#### 4.4.1 Idea density (ID)

Participants were given 10 minutes to produce a written text sample of approximately 300 words in response to the prompt “Describe an unexpected event in your life”. From this text, we calculated ID using the automated Computerized Propositional Idea Density Rater (CPIDR; Brown et al., 2008).

#### 4.4.2 Cognitive tests

Participants completed an additional battery of cognitive tests. These included:

- The four-subtest version of the Wechsler Abbreviated Scale of Intelligence Second Edition (WASI-II; Pearson Clinical), comprising Block design, Vocabulary, Matrix reasoning and Similarities tasks
- Three subtests from the Test of Adolescent and Adult Language-Fourth Edition (TOAL-4), namely Word opposites, Derivations and Spoken analogies
- Semantic and phonological verbal fluency tasks
- A computer-based hearing test to measure pure-tone hearing thresholds (pure-tone audiometry)

### 4.5 Procedure

Participants completed two in-lab testing sessions: (1) a behavioural session comprising the cognitive tests/text sample production, and (2) an EEG session comprising the collection of resting-state EEG recordings as well as the main language comprehension task. Sessions were either completed on the same day, separated by a break (approx. 30 minutes), or on two days (with the second session completed within 7 days of the first session).

#### 4.5.1 Behavioural session

After the consent process, participants provided demographic, language and well-being details and subsequently completed the cognitive tests. The behavioural session took maximally 1.5 hours to complete.

#### 4.5.2 EEG session

In the EEG session, participants were fitted with an EEG cap and first underwent a 2-minute eyes-open and 2-minute eyes-closed resting state EEG recording. The resting-state recordings were repeated at the end of the experimental session. Note that a subset of participants completed two (rather than one) eyes-closed resting state EEG recording sessions both before and after the experiment: one in which they were instructed to relax and one in which they were asked to try to keep their mind blank. For the purposes of calculating resting-state individual difference metrics (IAF and 1/f slope), we used the eyes-closed recordings with the “relax” instructions, as these were comparable to the eyes-closed resting-state recordings with only a single session.

In addition, participants completed a short (approximately 3.5 minute) passive auditory oddball paradigm from the ERP Core package (Kappenman et al., 2020) prior to the main language processing task (following Kurthen et al., 2020). For this task, they were presented with 290 1,000 Hz sine wave tones (100 ms duration) with a volume modulation: 230 standards at a volume of 80 dB and 60 deviants at a volume of 70 dB. The inter-stimulus interval was jittered between 450 and 550 ms.

In the main language processing task, trials commenced with a fixation asterisk, presented for 500 ms in the centre of a computer screen, followed by the auditory presentation of a passage via loudspeakers. After audio offset, the fixation asterisk remained on screen for another 500 ms. A comprehension question followed in approximately 1/3 of all trials, with participants responding “yes” or “no” via a game controller (maximal response time: 4000 ms). Assignment of “yes” and “no” responses to the left and right controller buttons was counterbalanced across participants. When there was no comprehension question, participants were asked to “Press the YES key to proceed”. The next trial commenced after an inter-trial interval of 1500 ms. Participants were asked to avoid any movements or blinks during the presentation of the fixation asterisk if possible.

Comprehension data was not analysed in the present paper as the comprehension task was merely intended to ensure attentive processing.

The 150 passages were presented in 5 blocks, separated by short self-paced breaks. Prior to commencing the main task, participants completed a short practice session. After the main task, the resting state recordings were repeated. Overall, the EEG session took approximately 3 hours including electrode preparation and participant clean-up.

### 4.6 EEG recording and preprocessing

The EEG was recorded from 64 electrodes mounted inside an elastic cap (actiCAP) using a Brain Products actiCHamp amplifier (Brain Products GmbH, Gilching, Germany). The electrooculogram (EOG) was recorded via electrodes placed at the outer canthi of both eyes as well as above and below the left eye. The EEG recording was sampled at 500 Hz and referenced to FCz.

Data preprocessing was undertaken using MNE Python version 1.0.3 (Gramfort et al., 2013; Gramfort et al., 2014). All raw data files were first converted to the brain imaging data structure for electroencephalography (EEG-BIDS; Pernet et al., 2019) using the MNE-BIDS Python package version 0.10 (Appelhoff et al., 2019). Subsequently, EEG data were re-referenced to an average reference. EOG artefacts were corrected using an ICA-based correction procedure, with independent components (ICs) found to correlate most strongly with EOG events (via the `create_eog_epochs` function in MNE) excluded. Raw data were filtered using a 0.1 – 30 Hz bandpass filter to exclude slow signal drifts and high frequency noise.

For the main language processing task, epochs were extracted in a time window from -200 to 1000 ms relative to critical word (ADJ2) onset and mean single-trial amplitudes were extracted for the prestimulus (-200–0 ms) and N400 (300–500 ms) time windows using the retrieve function from the philistine Python package (Alday, 2018).

For the auditory oddball task, epochs were extracted in a time window from -200 to 1000 ms relative to the onset of each tone and mean single-trial amplitudes were extracted for the prestimulus (-200–0 ms), MMN (110–180 ms) and P3 (200–300 ms) time windows in a frontocentral region of interest comprising electrodes Fz, FC1, FC2 and Cz. The ROI specification and MMN time window were adopted from Kurthen et al. (2020), while the P3 time window was selected via visual inspection of the grand average ERPs.

### 4.7 Resting-state EEG-based individual differences measures: Individual alpha frequency (IAF) and aperiodic (1/f) activity

IAF and aperiodic slope estimates were calculated from participants’ eyes-closed resting-state recordings.

IAF (peak alpha frequency) was estimated from electrodes P1, Pz, P2, PO3, POz, PO4, O1 and O2 using a Python-based implementation (Alday, 2018) of the procedure described in Corcoran et al. (2018) We calculated the mean of pre and post estimates for use as an individual differences metric.

Aperiodic (1/f) intercept and slope estimates were calculated in Python using the YASA toolbox (Vallat & Walker, 2021). YASA implements the irregular-resampling auto-spectral analysis (IRASA) method for separating oscillatory and aperiodic activity (Wen & Liu, 2016). As for IAF, by-participant intercept and slope estimates were computed as means of pre and post restingstate recordings from electrodes: F7, F3, Fz, F4, F8, FC5, FC1, FC2, FC6, T7, C3, Cz, C4, T8, CP5, CP1, CP2, CP6, P7, P3, Pz, P4, P8, PO9, O1, O2, PO10, AF7, AF8, F5, F1, F2, F6, FT7, FC3, FC4, FT8, C5, C1, C2, C6, TP7, CP3, CPz, CP4, TP8, P5, P1, P2, P6, PO7, PO3, POz, PO4, PO8.

### 4.8 Data analysis

Data analysis was undertaken using R (R Core Team, 2021) and Julia (Bezanson et al., 2017). For data import and manipulation, we used the `tidyverse` collection of packages (Wickham et al., 2019) as well as the `vroom` package (Hester & Wickham, 2021). Figures displaying the distribution and correlation of individual differences predictors were created in R using `ggplot2`. Other packages used include `corrr` (Kuhn et al., 2020), `kableExtra` (Zhu, 2021) and `here` (Müller, 2020). For package version numbers, please see the analysis scripts provided with the raw data (see Data Availability Statement). For R, see the html outputs in the src/ subdirectory; for Julia see the Manifest.toml file.

EEG data were analysed using linear mixed effects models (LMMs) with the `MixedModels.jl` package in Julia (Bates et al., 2021). Effects plots were created in Julia using the `Effects.jl` (Alday et al., 2022) and `AlgebraOfGraphics.jl` packages.

For the ERP data in the main language processing task, we examined single-trial N400 amplitude as our outcome variable of interest. As in Bornkessel-Schlesewsky et al. (2022), we analysed mean EEG voltage in a time window from 300 to 500 ms post onset of the critical second adjective (ADJ2) in a centro-parietal region of interest (C3, C1, Cz, C2, C4, P3, P1, Pz, P2, P4, CP3, CP1, CPz, CP2, CP4).

#### 4.8.1 Linear mixed modelling (LMM) approach

The modelling approach was analogous to that adopted in Bornkessel-Schlesewsky et al. (2022). We used a parsimonious LMM selection approach (Bates et al., 2015; Matuschek et al., 2017) to identify LMMs that are supported by the data and not overparameterised.

Fixed effects initially included log-transformed unigram frequency, speaker-based surprisal, adjective order canonicity, epoch (as a proxy for time-on-task), mean prestimulus amplitude and their interactions. Prestimulus amplitude (-200-0 ms) was included as a predictor in the model as an alternative to traditional EEG baselining (see Alday, 2019). The categorical factor canonicity was encoded using sum contrasts (cf. Brehm & Alday, 2022; Schad et al., 2020) such that model intercepts represent the grand mean. All continuous predictors were z-transformed prior to being included in the models.

Although not of interest within the scope of the current paper, we modelled the main effect of prestimulus amplitude with a second-order and the main effect of speaker-based surprisal with a third-order polynomial trend. The inclusion of these higher-order trends was supported by the data and significantly improved model fit. It further guarded against the interpretation of spurious interactions of their linear trends with other fixed effects (Matuschek & Kliegl, 2018).

The random-effect (RE) structure was selected in two steps, using likelihood-ratio tests (LRTs) to check improvement in goodness of fit and random-effects PCA (rePCA) to guard against overparameterisation during model selection.

For the N400 model including only age and no additional individual differences predictors, this led to a RE structure with variance components for grand means, prestimulus amplitude and prestimulus amplitude (2nd order) by subject, item and channel. In a second step, we added by-subject variance components for effects of canonicity, epoch, unigram frequency and speaker-based surprisal and by-item variance components for effects of epoch, unigram frequency, speaker-based surprisal and age.

Using the age-only LMM (as described above) as a reference, we added, in turn, fixed-effect covariates for individual differences in (1) 1/f slope, (2) 1/f intercept, (3) IAF (peak alpha frequency), (4) Idea Density, (5) Verbal Fluency, (6) General Cognitive Ability (WASI FSIQ-4) and (7) Language Proficiency (TOAL-4 composite score) to the model to check the extent to which they moderate / modulate adaptation to speaker-based surprisal. In each of these additional LMMs, adding the respective individual differences covariate as a by-item variance component significantly improved the goodness of model fit.

The model selection procedure is transparently documented in Julia scripts in the Open Science Framework repository for this paper (see Data Availability Statement).

## Acknowledgements

The authors would like to thank John Ellett for help with preparing the experimental materials, Alin Grecu and Casey Tonkin for recording the experimental stimuli and Nicole Vass for help with data collection. Thanks are also due to Zachariah Cross for helpful comments on a previous version of the manuscript.

## Data Availability Statement

Raw data and analysis code will be made available on a public repository upon publication of the manuscript.

## Appendix

### Appendix A Visualisation of residualisation models

**Figure 12:**
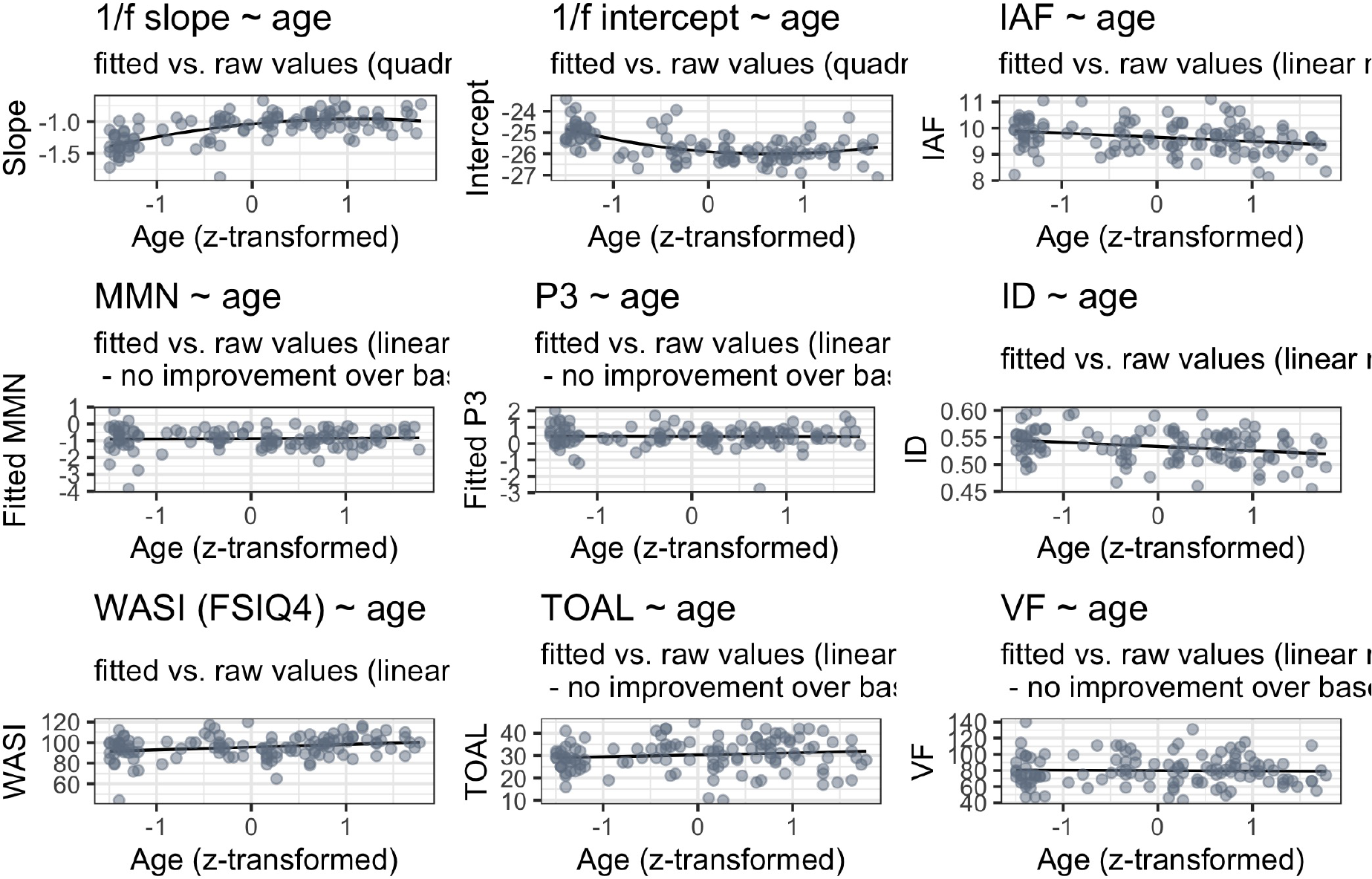
Visualisation of the results of the residualisation analysis for the individual differences predictors.

### Appendix B Supplementary figures

**Figure 13:**
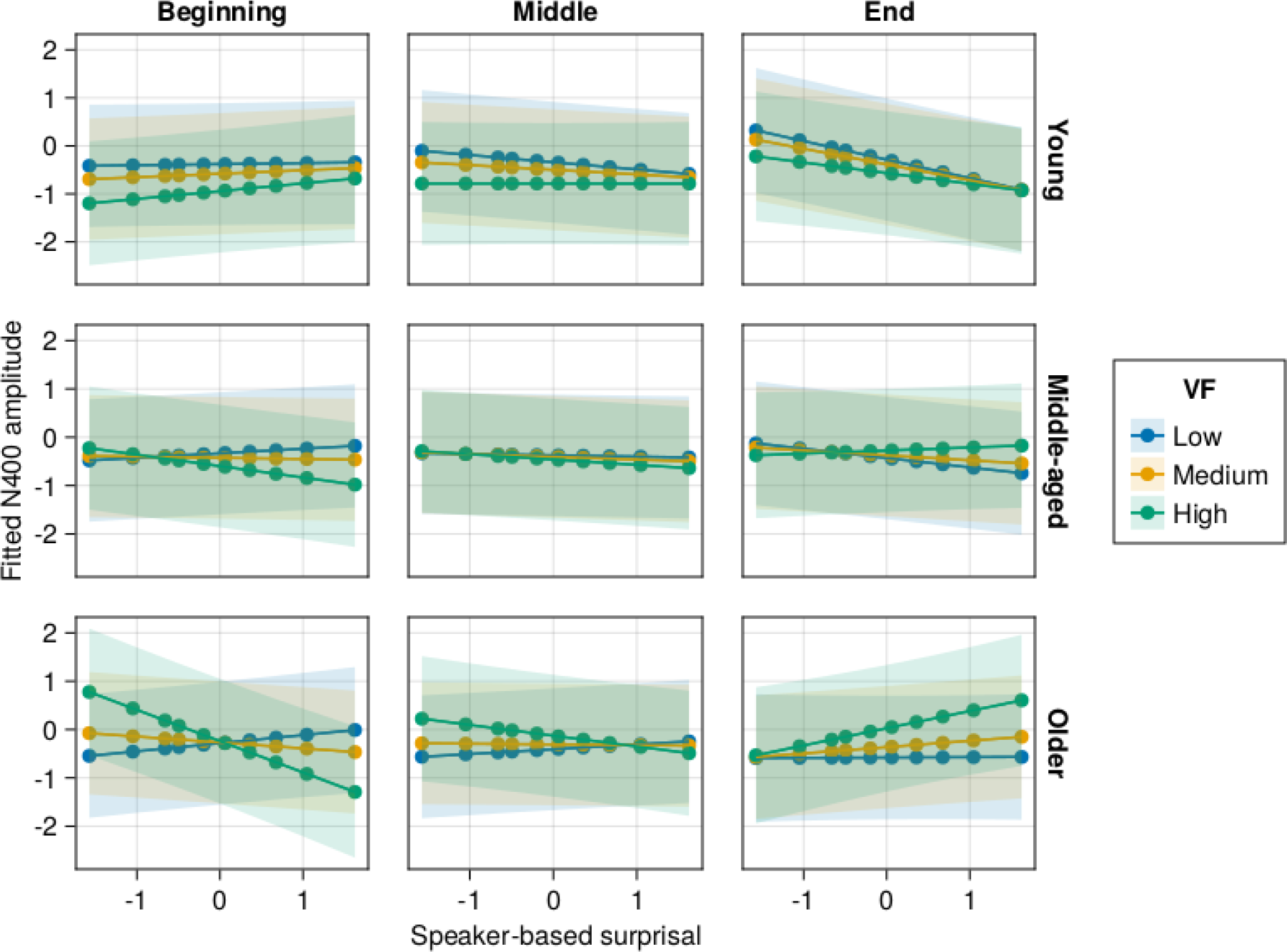
Effects of age and (age-residualised) Verbal Fluency (VF) on changes in the relationship between speaker-based surprisal and N400 amplitude over the course of the experiment. Note that position in the experiment (operationalised via epoch in the statistical model) is trichotomised into beginning (estimated for the 5th quantile), middle (50th quantile) and end (95th quantile) for visualisation purposes only; epoch was included in the model as a continuous predictor. The same holds for age, which is trichotomised for visualisation purposes (estimated for ages 18, 50 and 83 as the minimum, median and maximum of ages in our sample) but was entered into the statistical model as a continuous predictor. Shaded ribbons signify standard errors.

**Figure 14:**
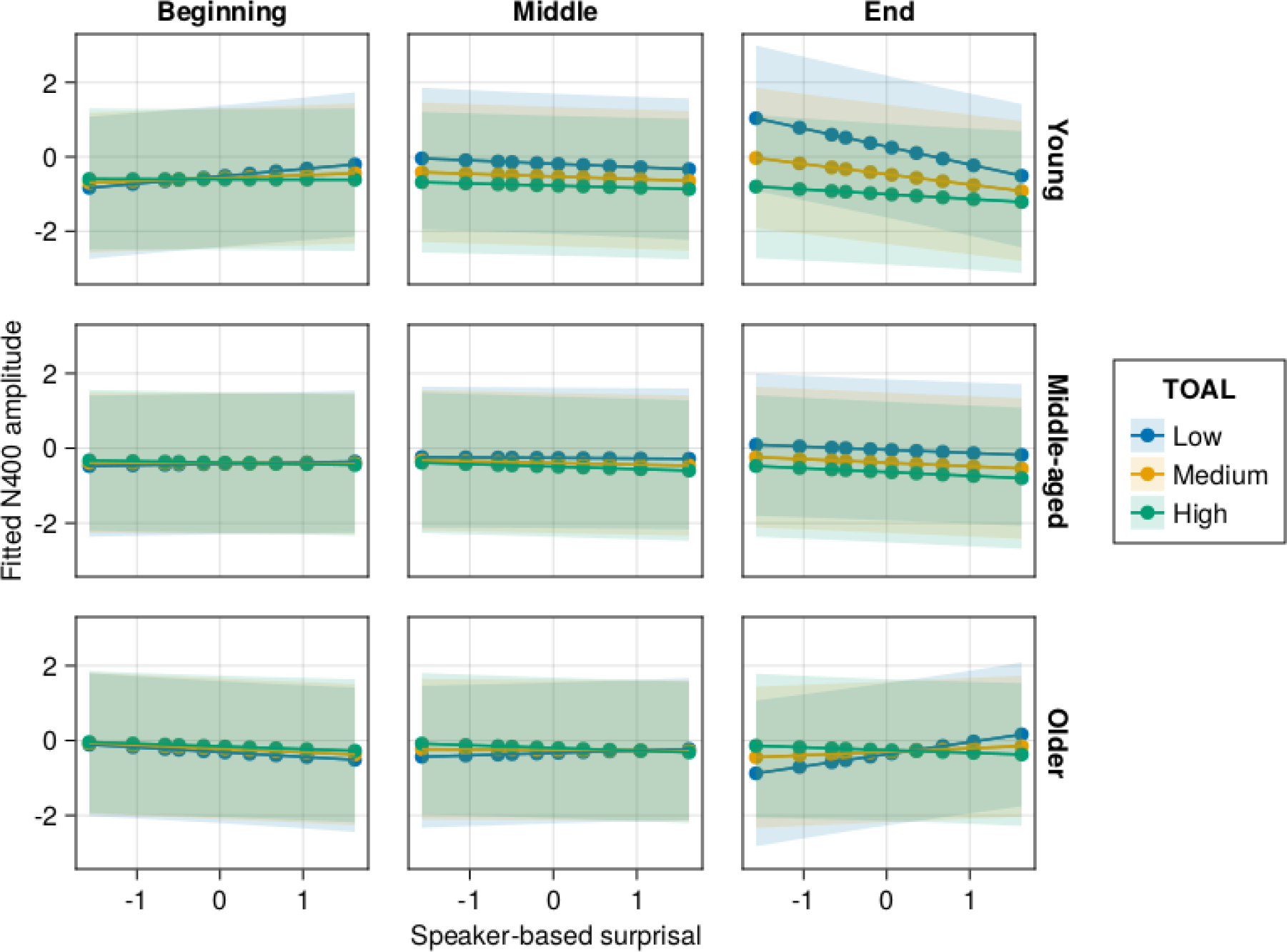
Effects of age and (age-residualised) oral language proficiency (TOAL-4) on changes in the relationship between speaker-based surprisal and N400 amplitude over the course of the experiment. Note that position in the experiment (operationalised via epoch in the statistical model) is trichotomised into beginning (estimated for the 5th quantile), middle (50th quantile) and end (95th quantile) for visualisation purposes only; epoch was included in the model as a continuous predictor. The same holds for age, which is trichotomised for visualisation purposes (estimated for ages 18, 50 and 83 as the minimum, median and maximum of ages in our sample) but was entered into the statistical model as a continuous predictor. Shaded ribbons signify standard errors.

**Figure 15:**
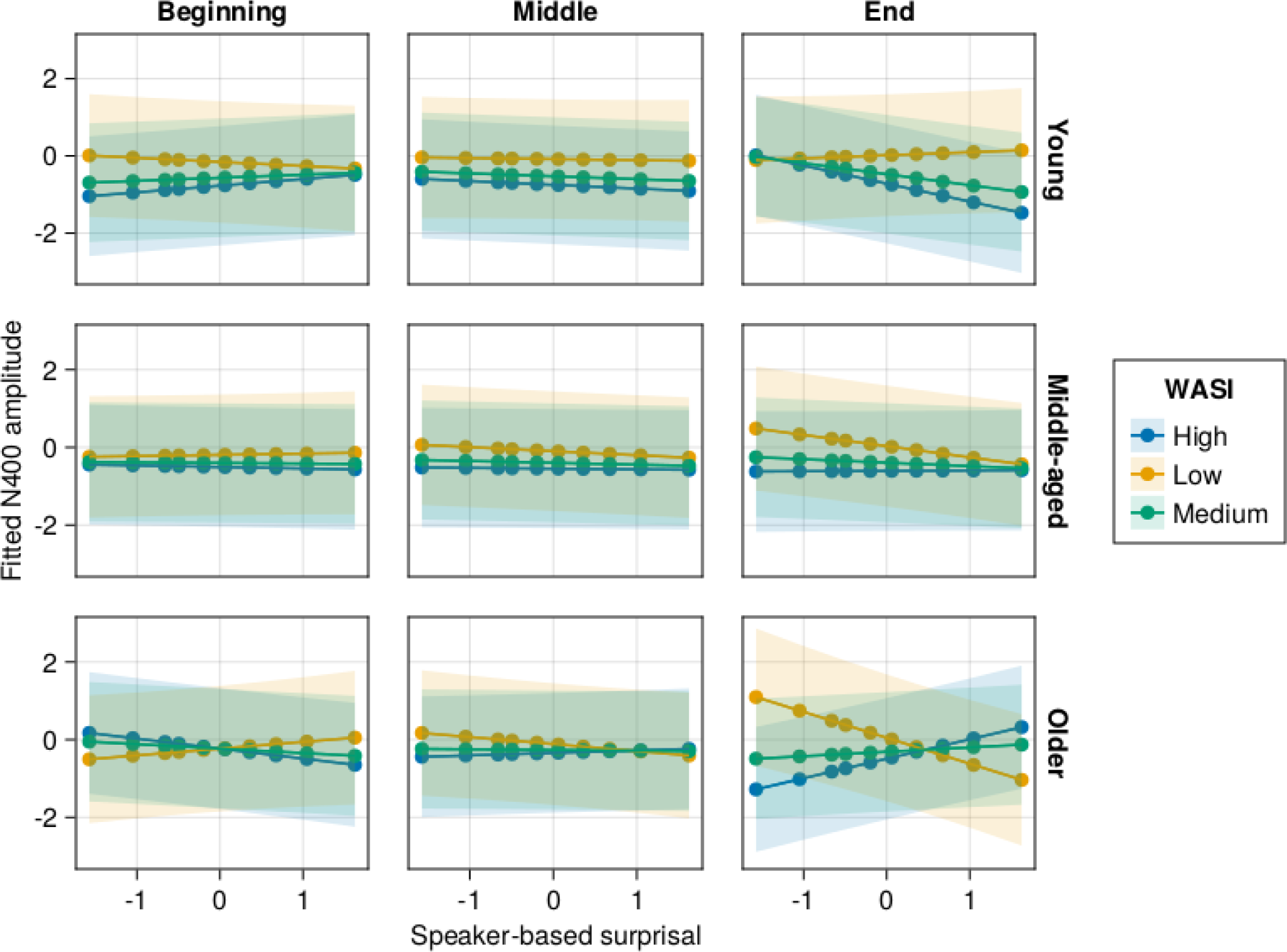
Effects of age and (age-residualised) general cognitive ability (WASI FSIQ-4) on changes in the relationship between speaker-based surprisal and N400 amplitude over the course of the experiment. Note that position in the experiment (operationalised via epoch in the statistical model) is trichotomised into beginning (estimated for the 5th quantile), middle (50th quantile) and end (95th quantile) for visualisation purposes only; epoch was included in the model as a continuous predictor. The same holds for age, which is trichotomised for visualisation purposes (estimated for ages 18, 50 and 83 as the minimum, median and maximum of ages in our sample) but was entered into the statistical model as a continuous predictor. Shaded ribbons signify standard errors.

### Appendix C Model summaries

**Table 3:**
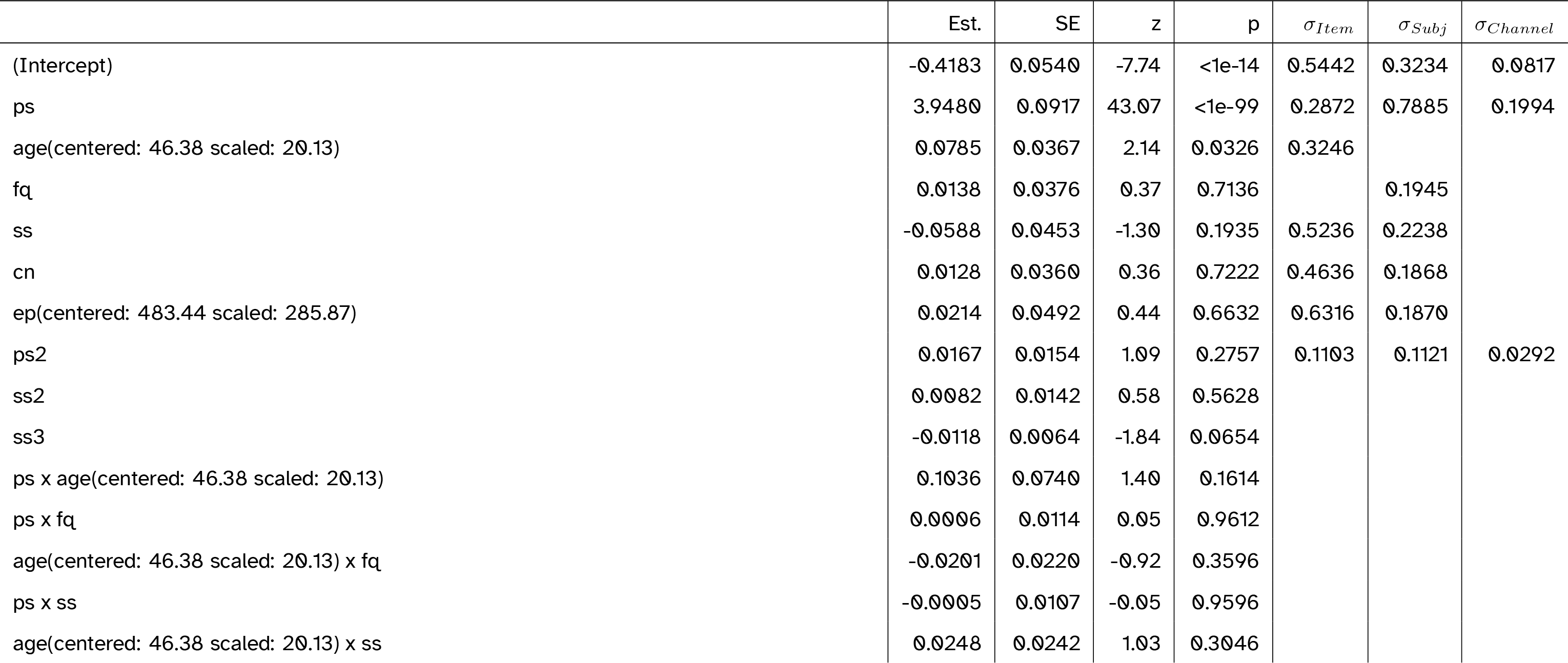
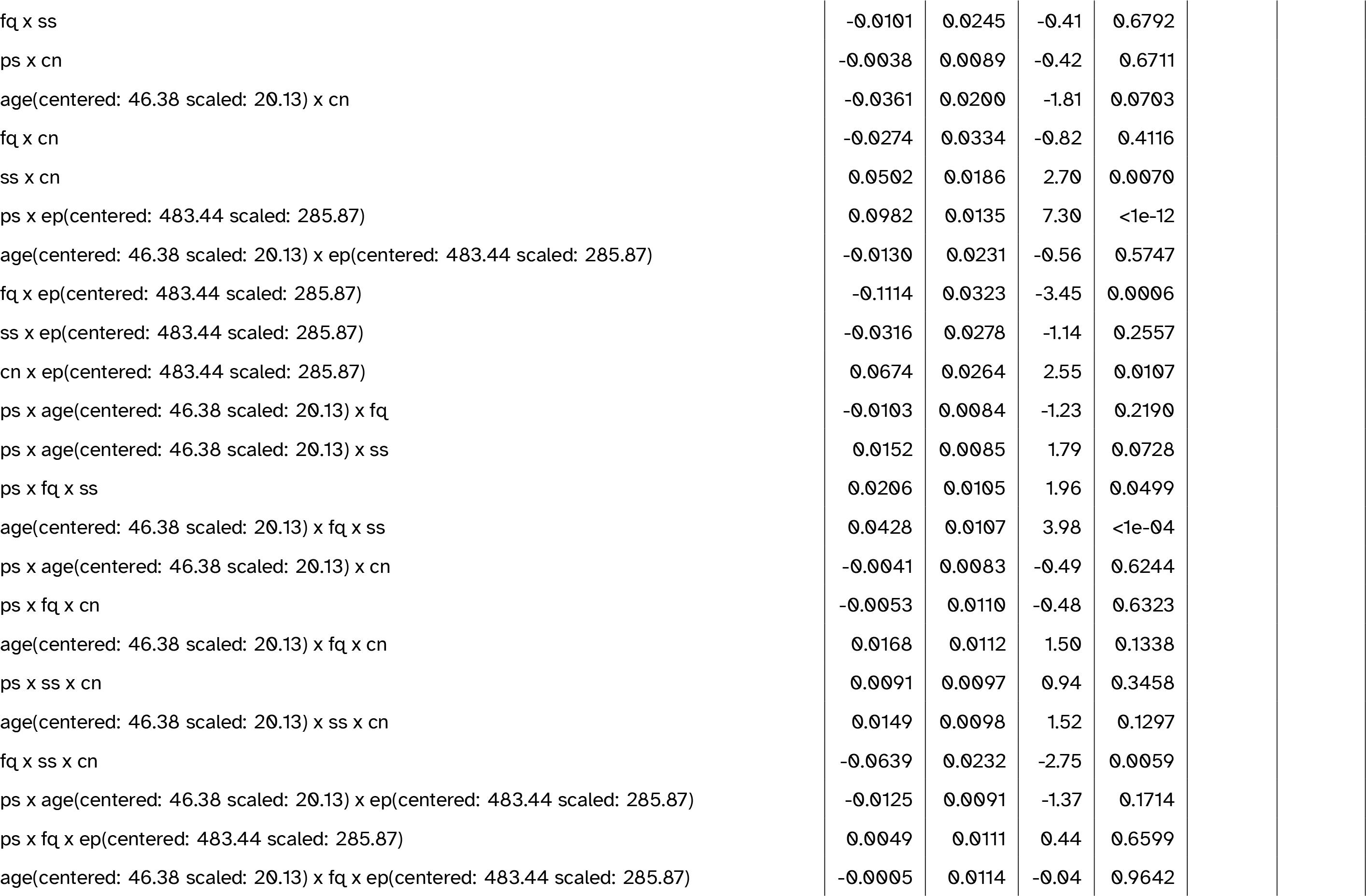
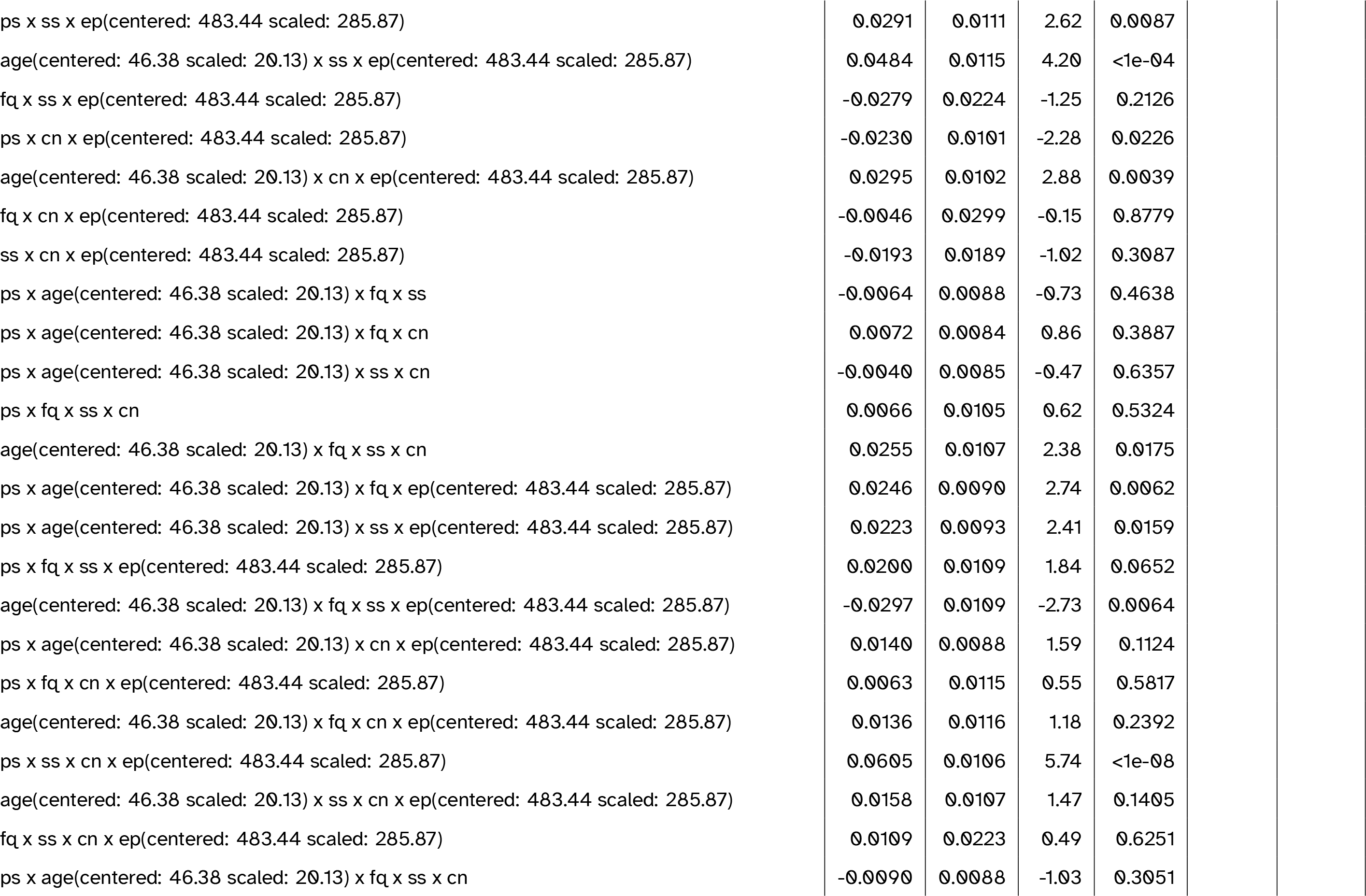
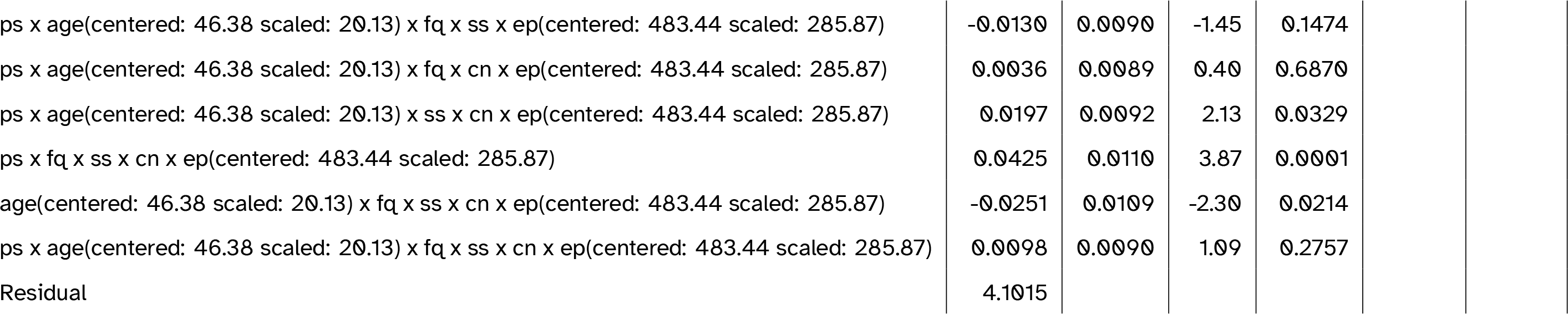
Full model summary for the best-fitting model including only age. Abbreviations: cn = canonicity; ep = epoch; fq = word frequency; ps = prestimulus amplitude; ps2 = quadratic prestimulus amplitude; ss = speaker-based surprisal; ss2 = quadratic speaker-based surprisal; ss3 = cubic speaker-based surprisal

**Table 4:**
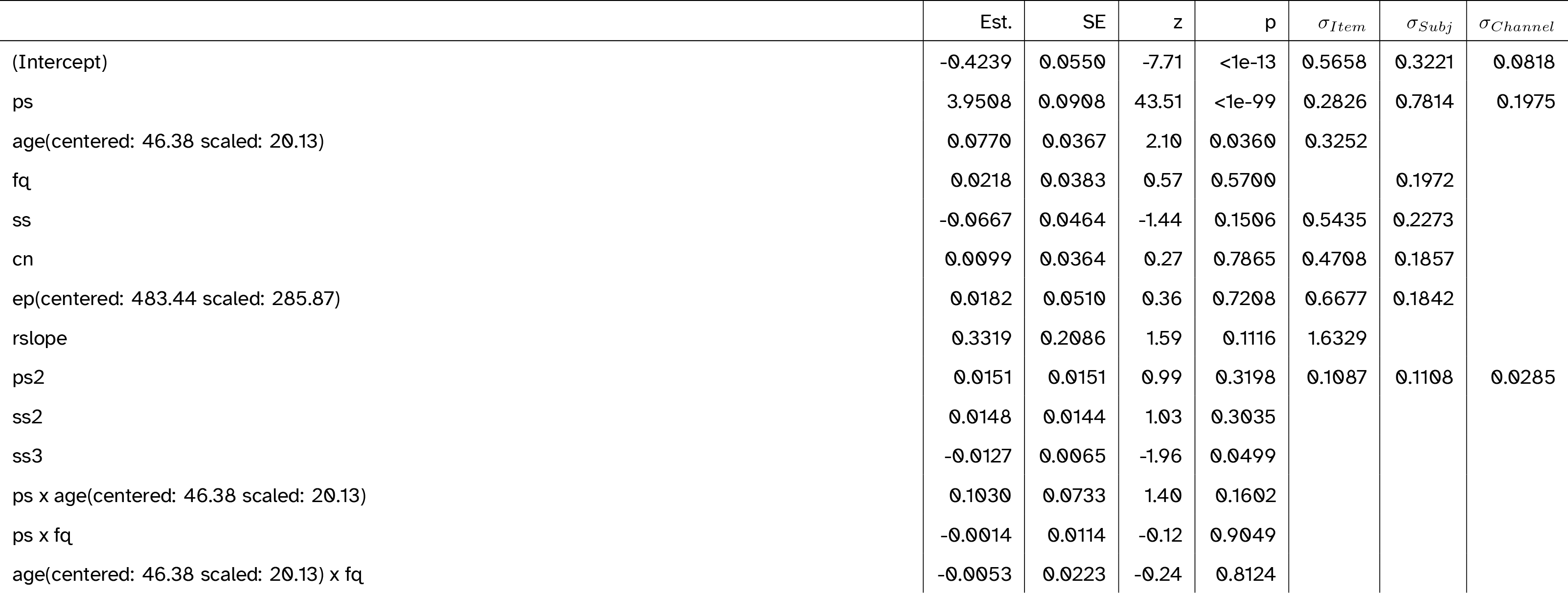
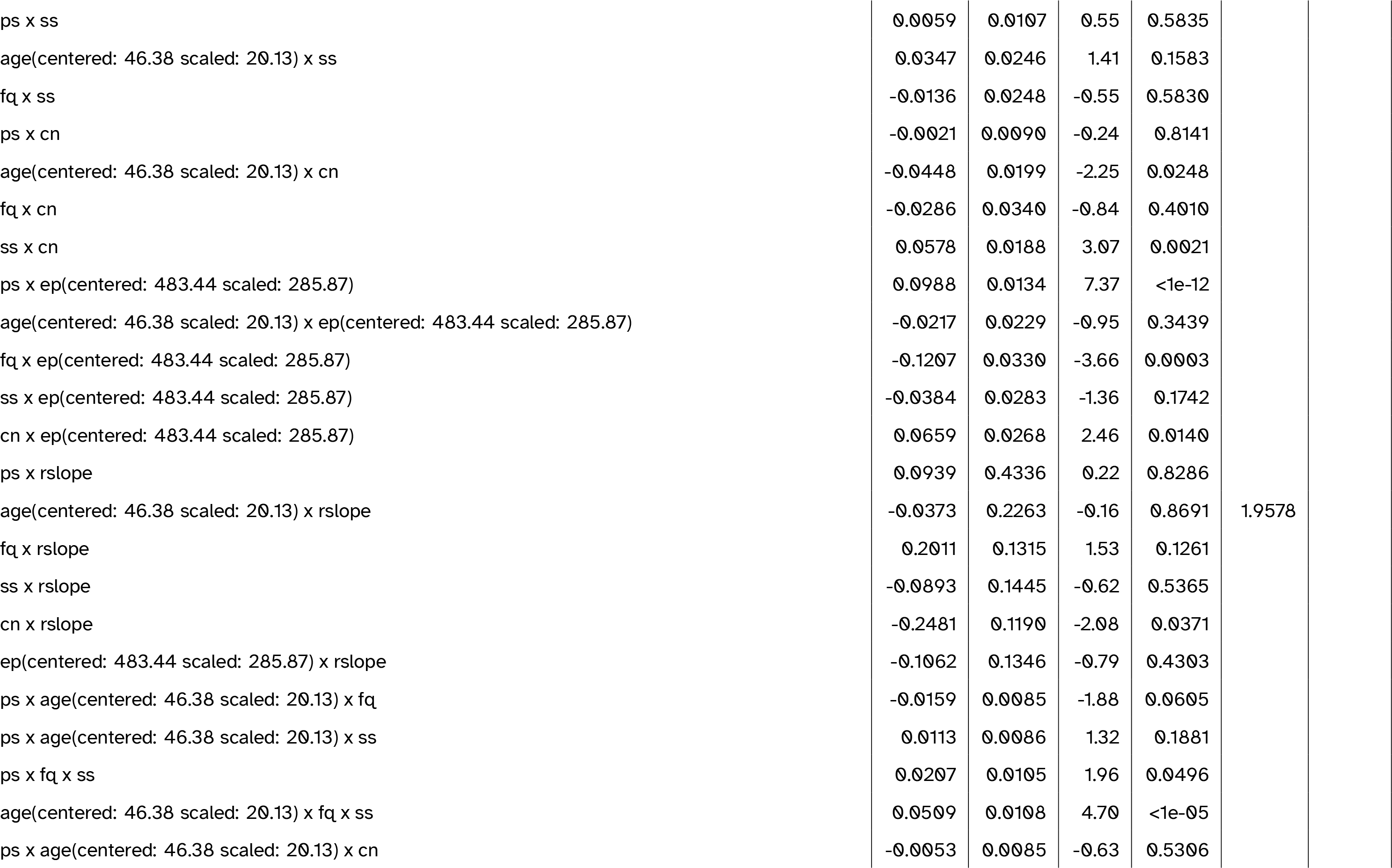
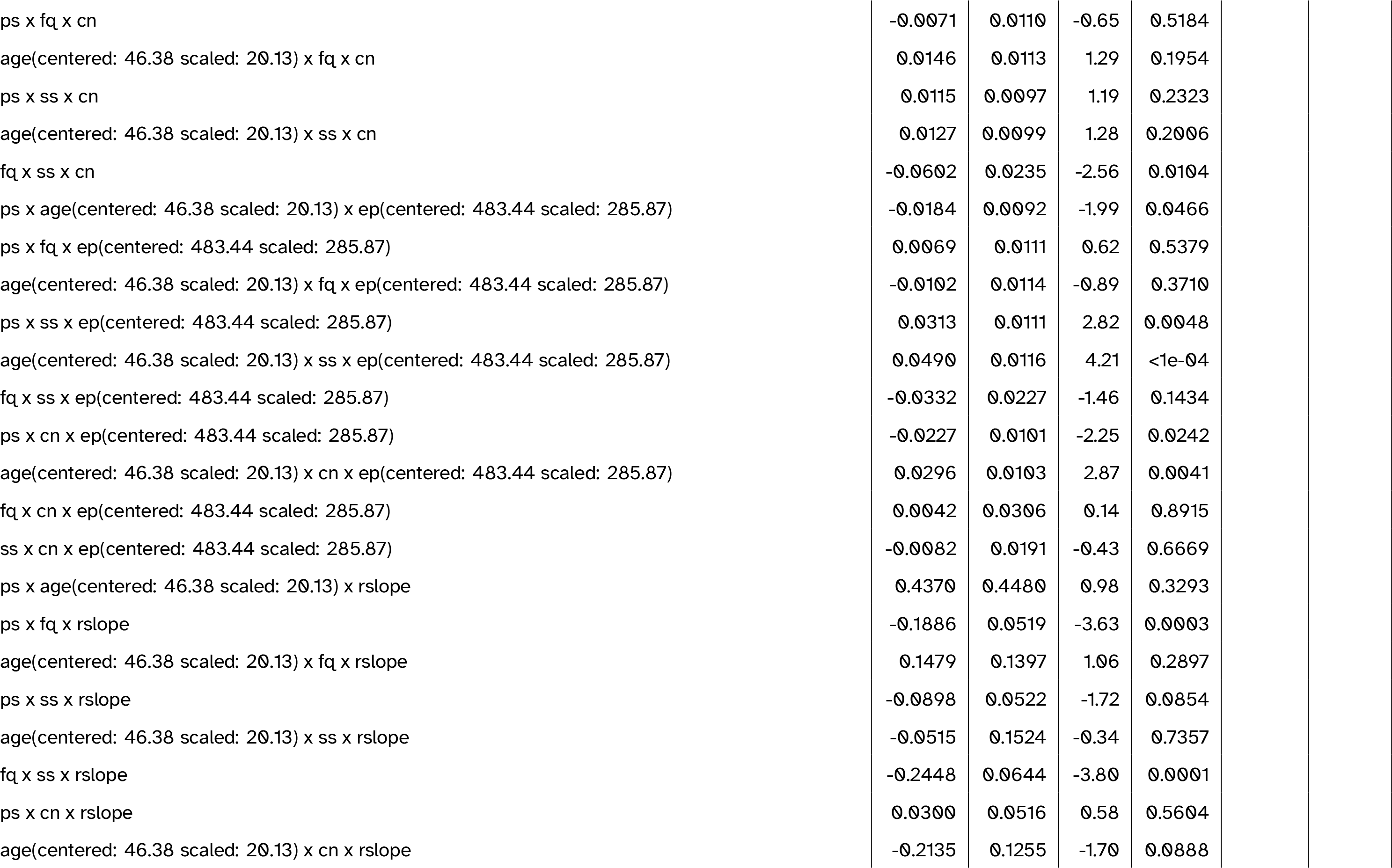
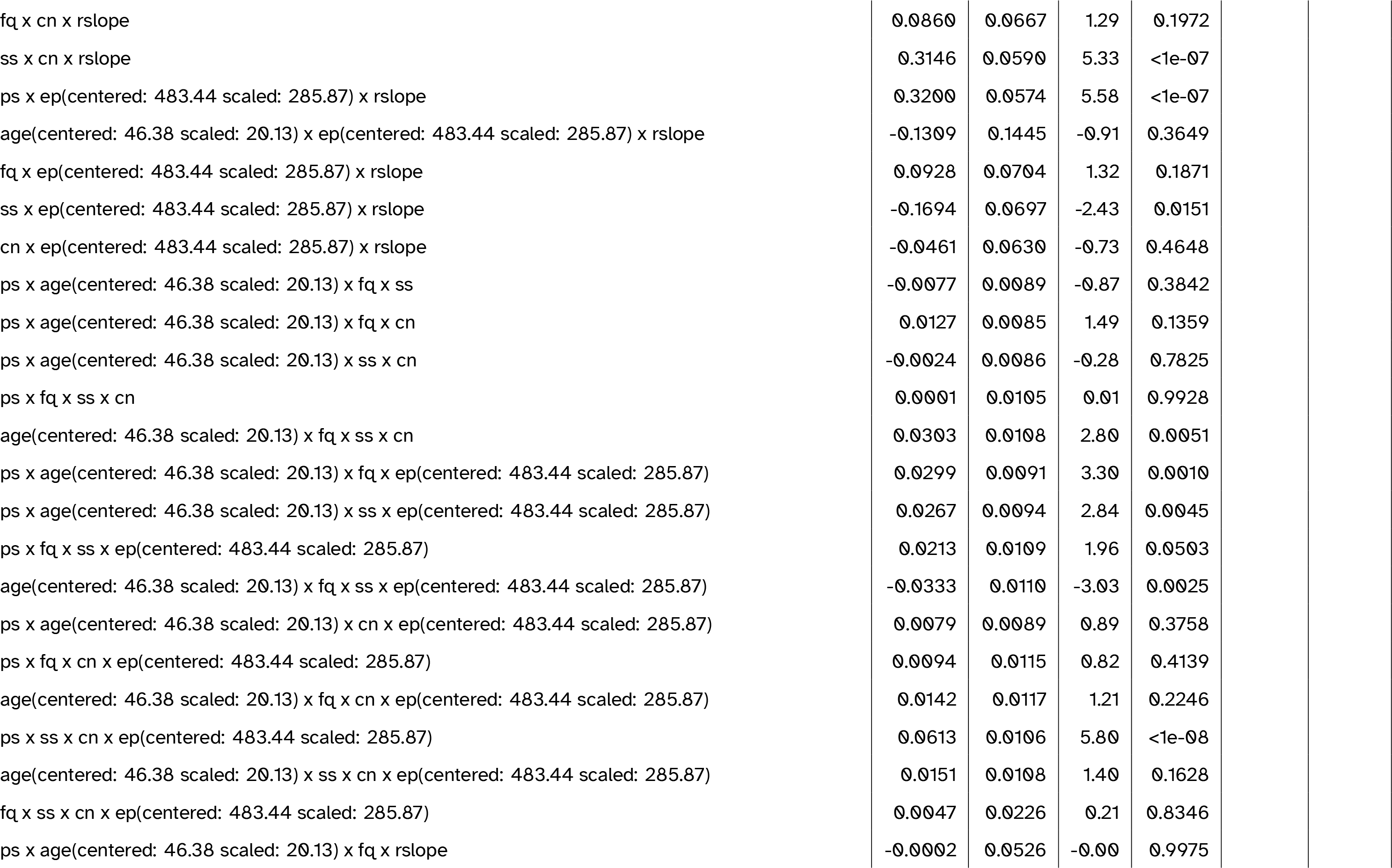
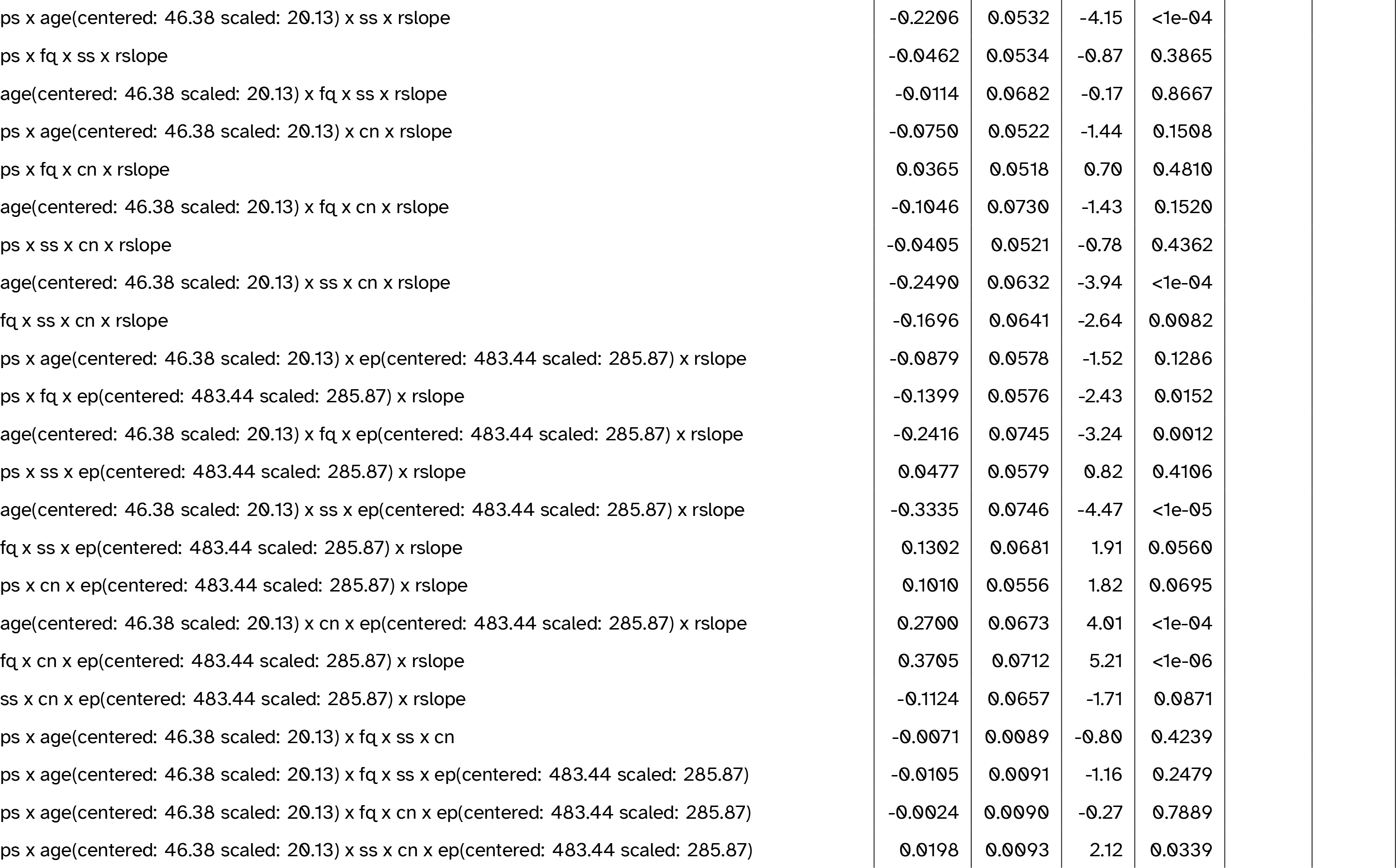
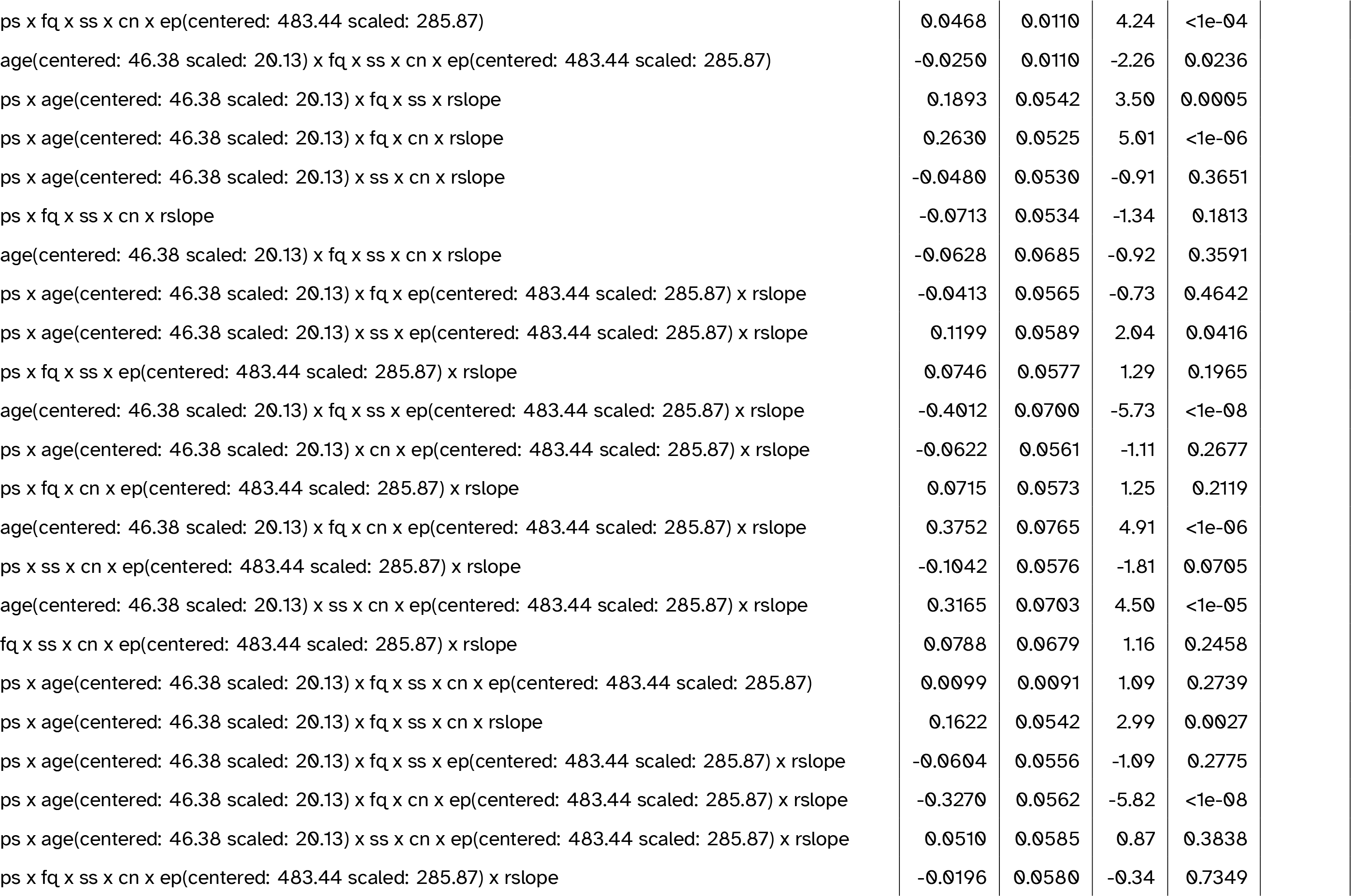
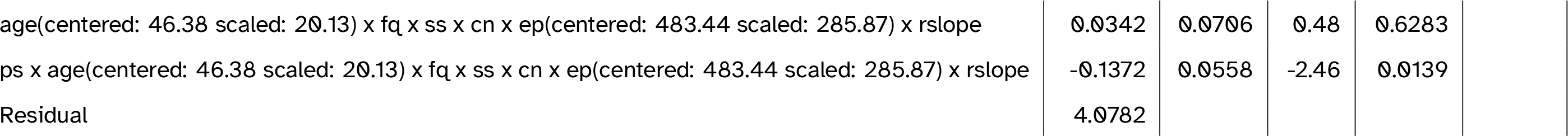
Full model summary for the best-fitting model including aperiodic (1/f) slope. Abbreviations: cn = canonicity; ep = epoch; fq = word frequency; ps = prestimulus amplitude; ps2 = quadratic prestimulus amplitude; rslope = aperiodic slope (residualised on age); ss = speaker-based surprisal; ss2 = quadratic speaker-based surprisal; ss3 = cubic speaker-based surprisal

**Table 5:**
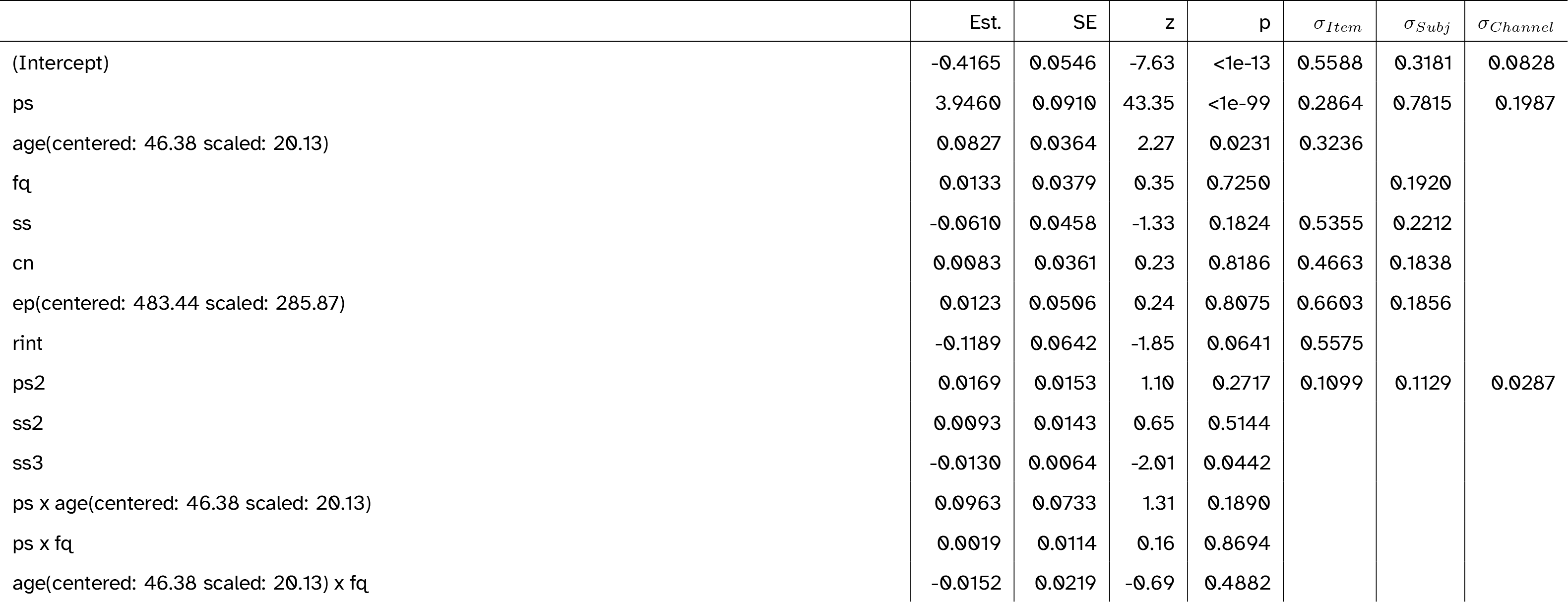
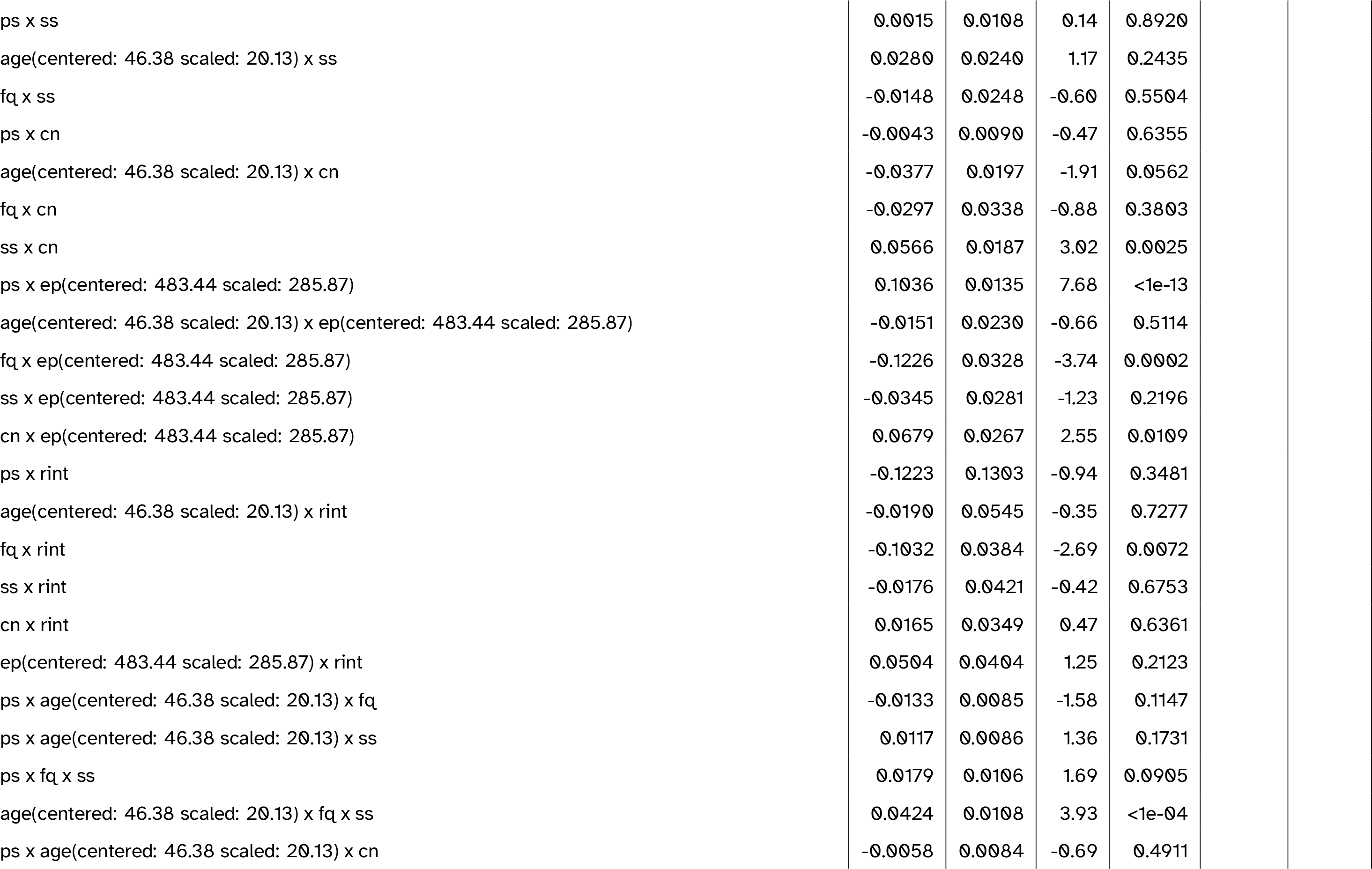
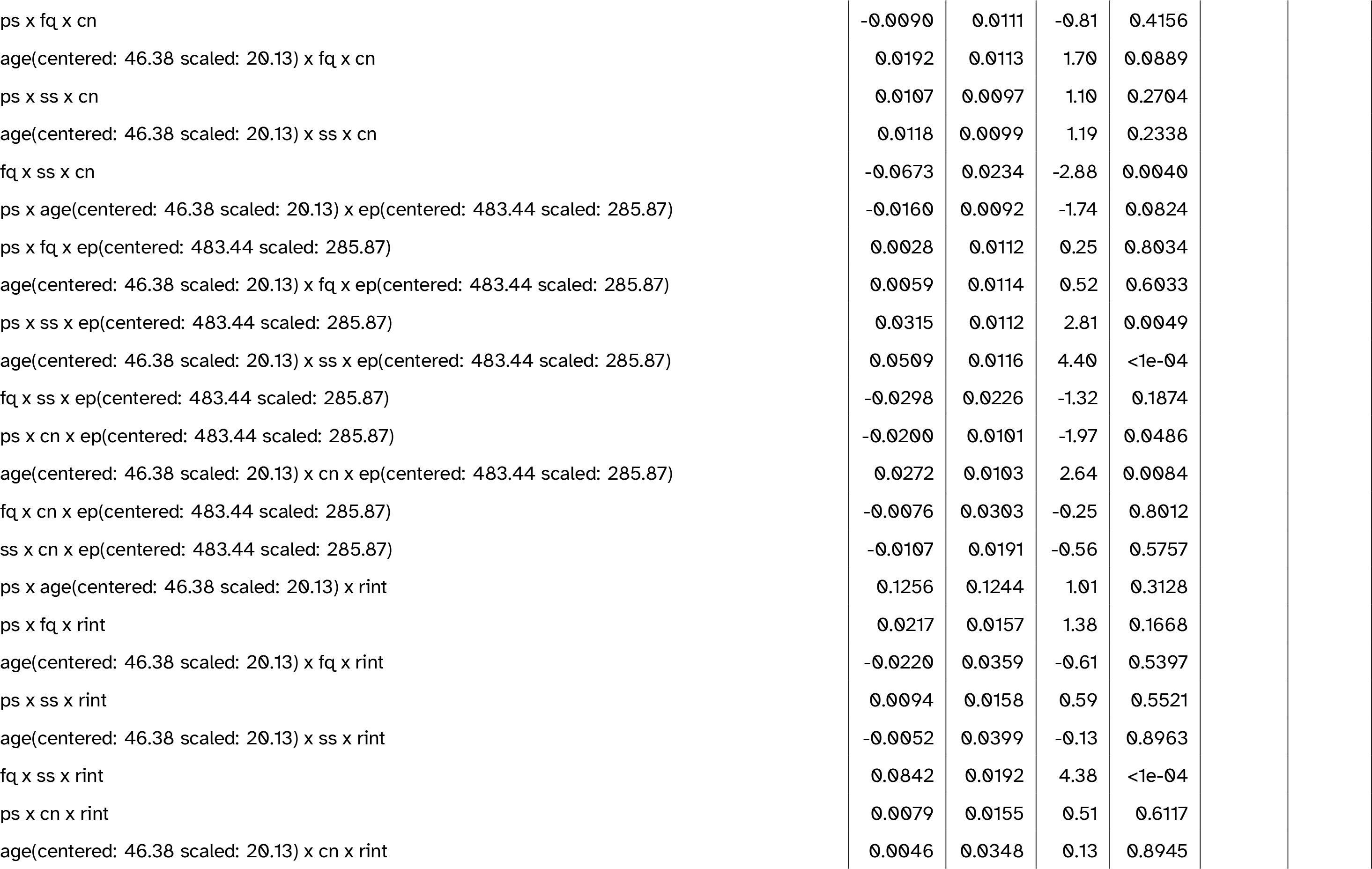
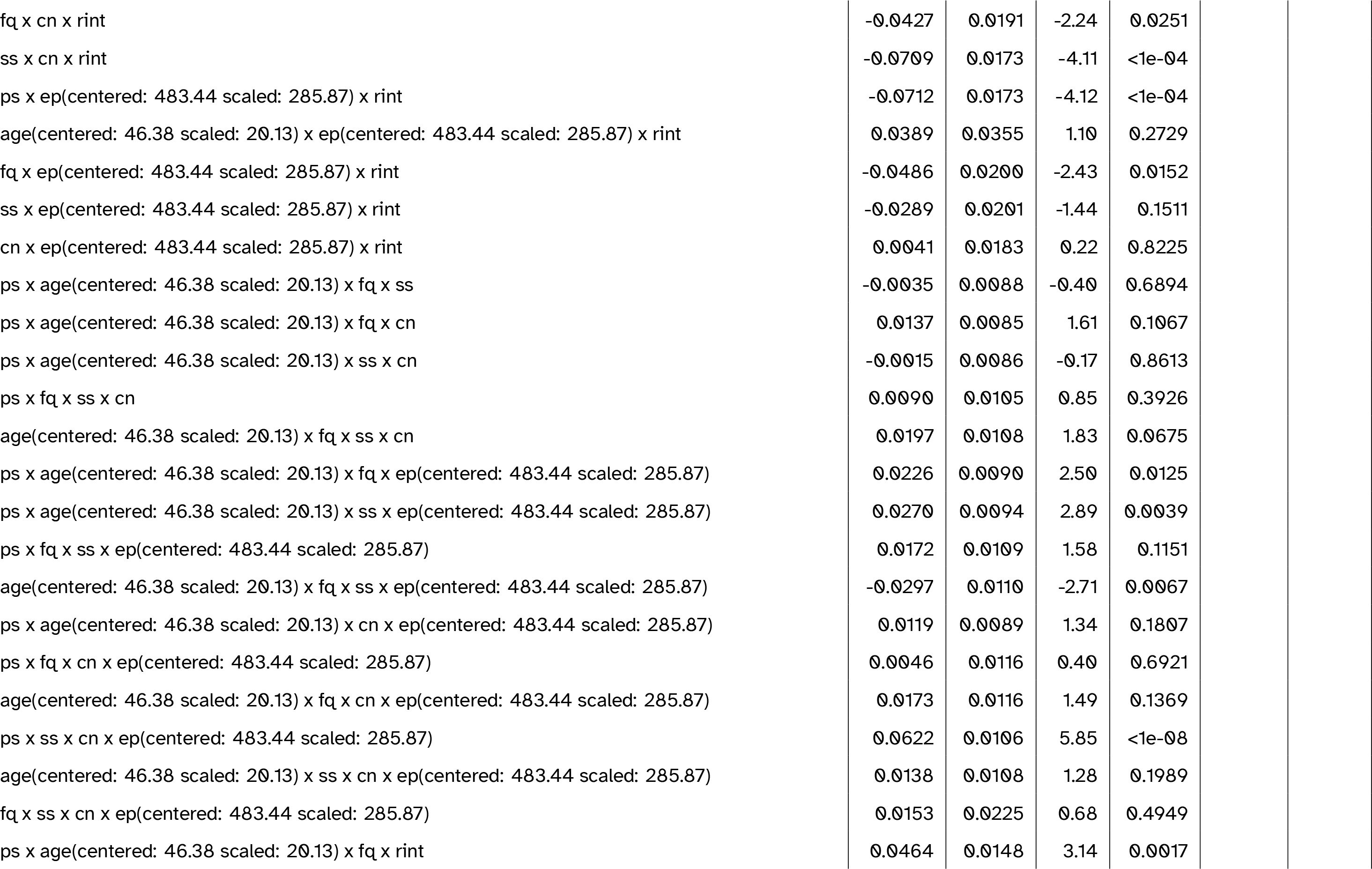
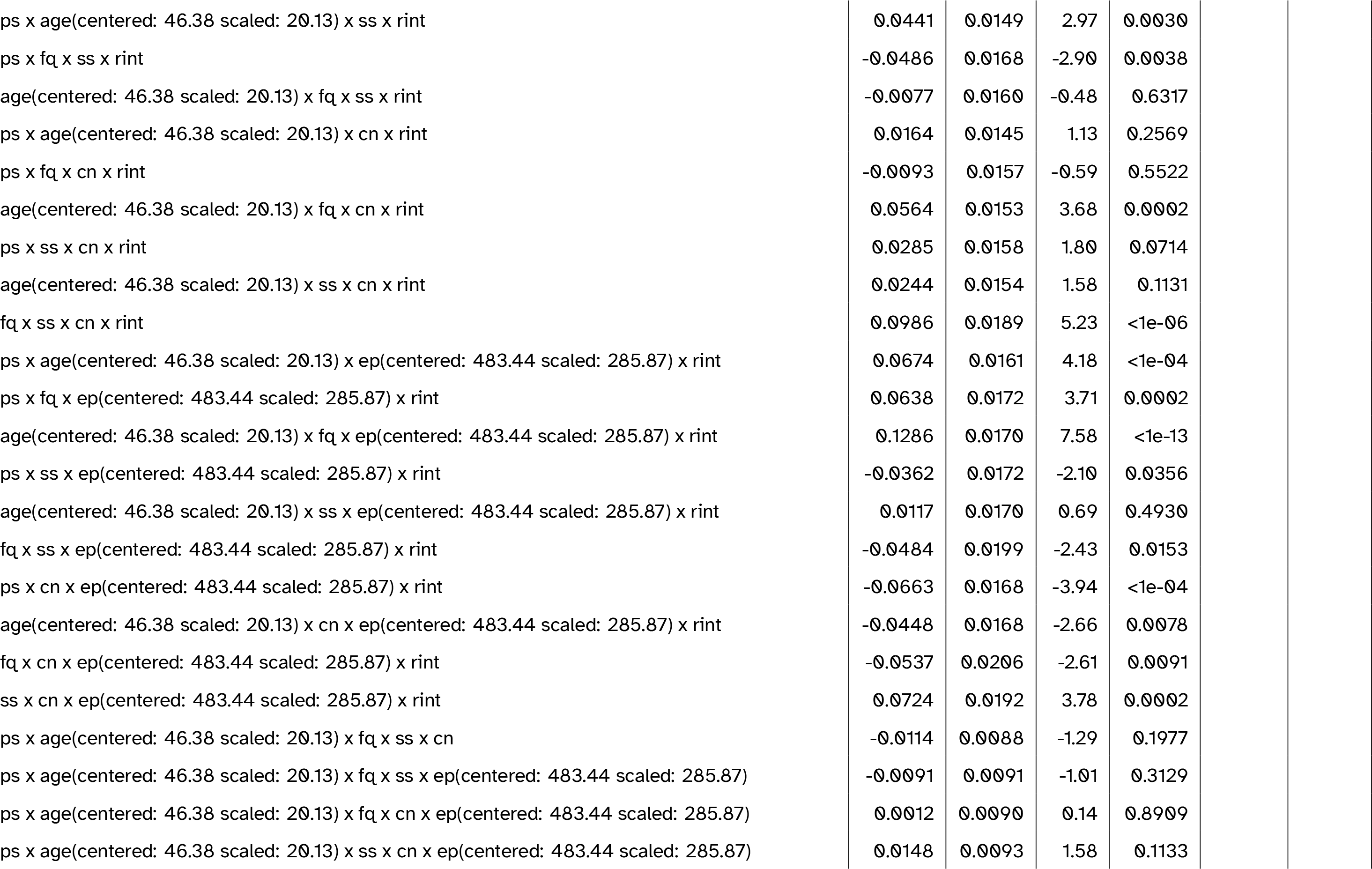
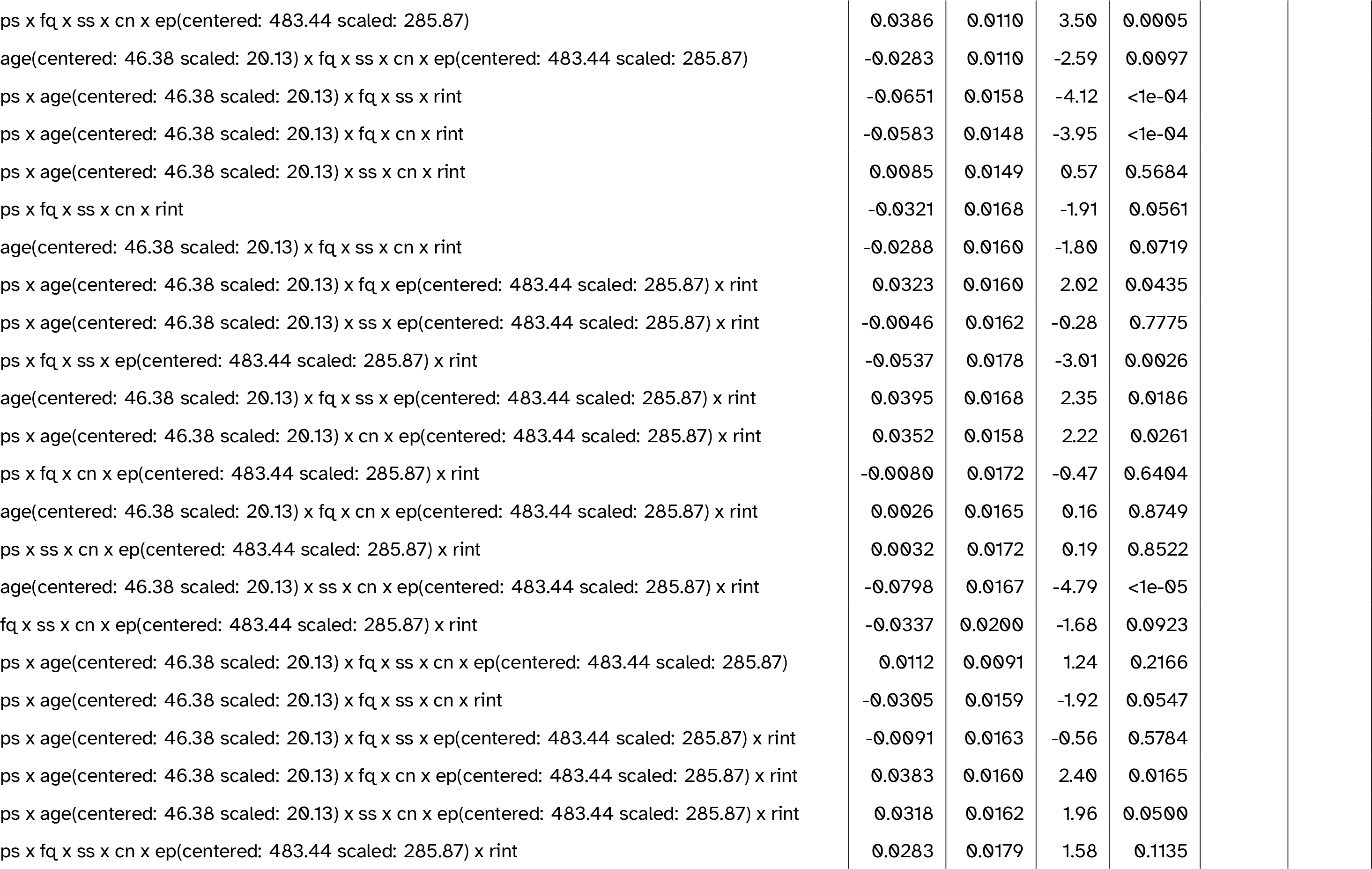
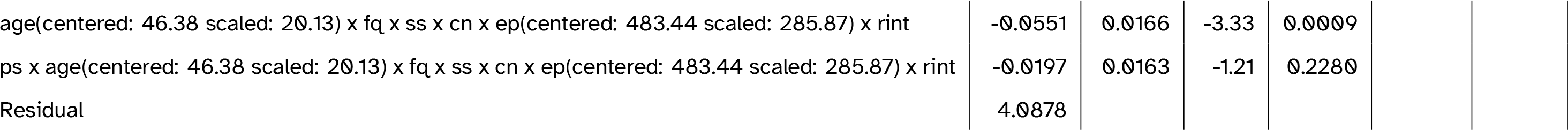
Full model summary for the best-fitting model including aperiodic (1/f) intercept. Abbreviations: cn = canonicity; ep = epoch; fq = word frequency; ps = prestimulus amplitude; ps2 = quadratic prestimulus amplitude; rint = aperiodic intercept (residualised on age); ss = speaker-based surprisal; ss2 = quadratic speaker-based surprisal; ss3 = cubic speakerbased surprisal

**Table 6:**
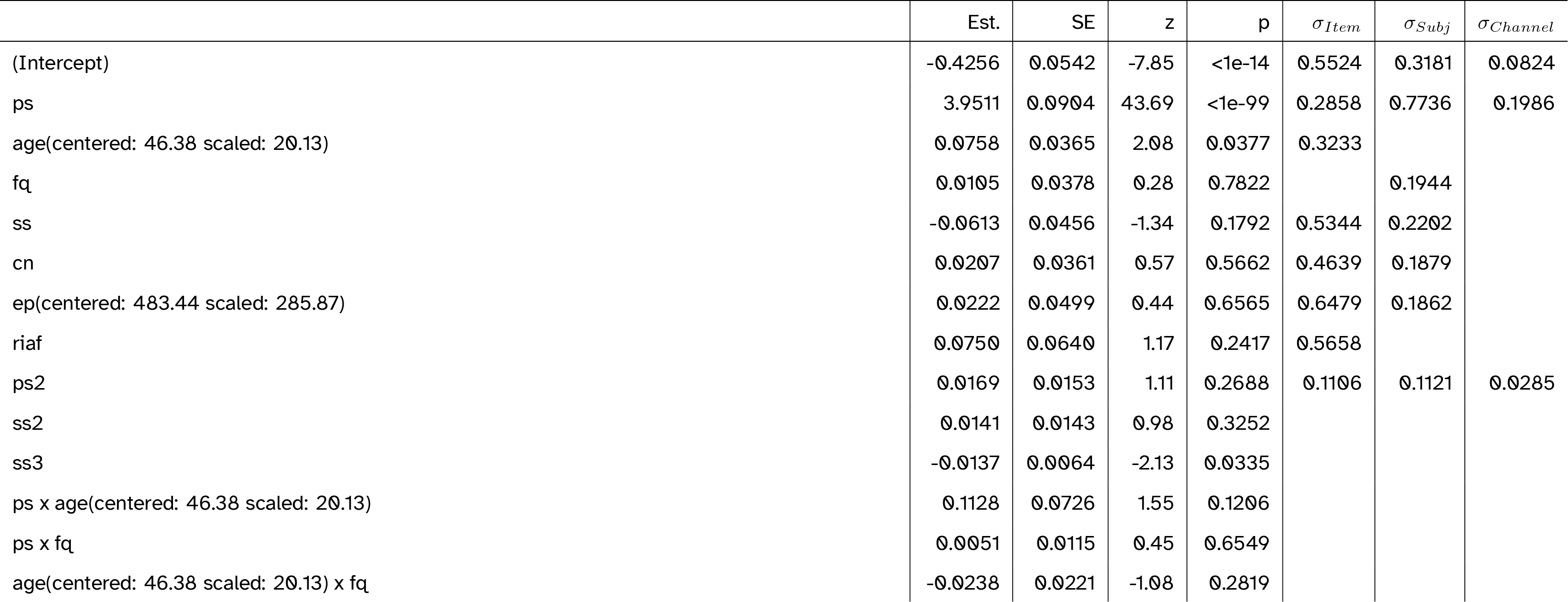
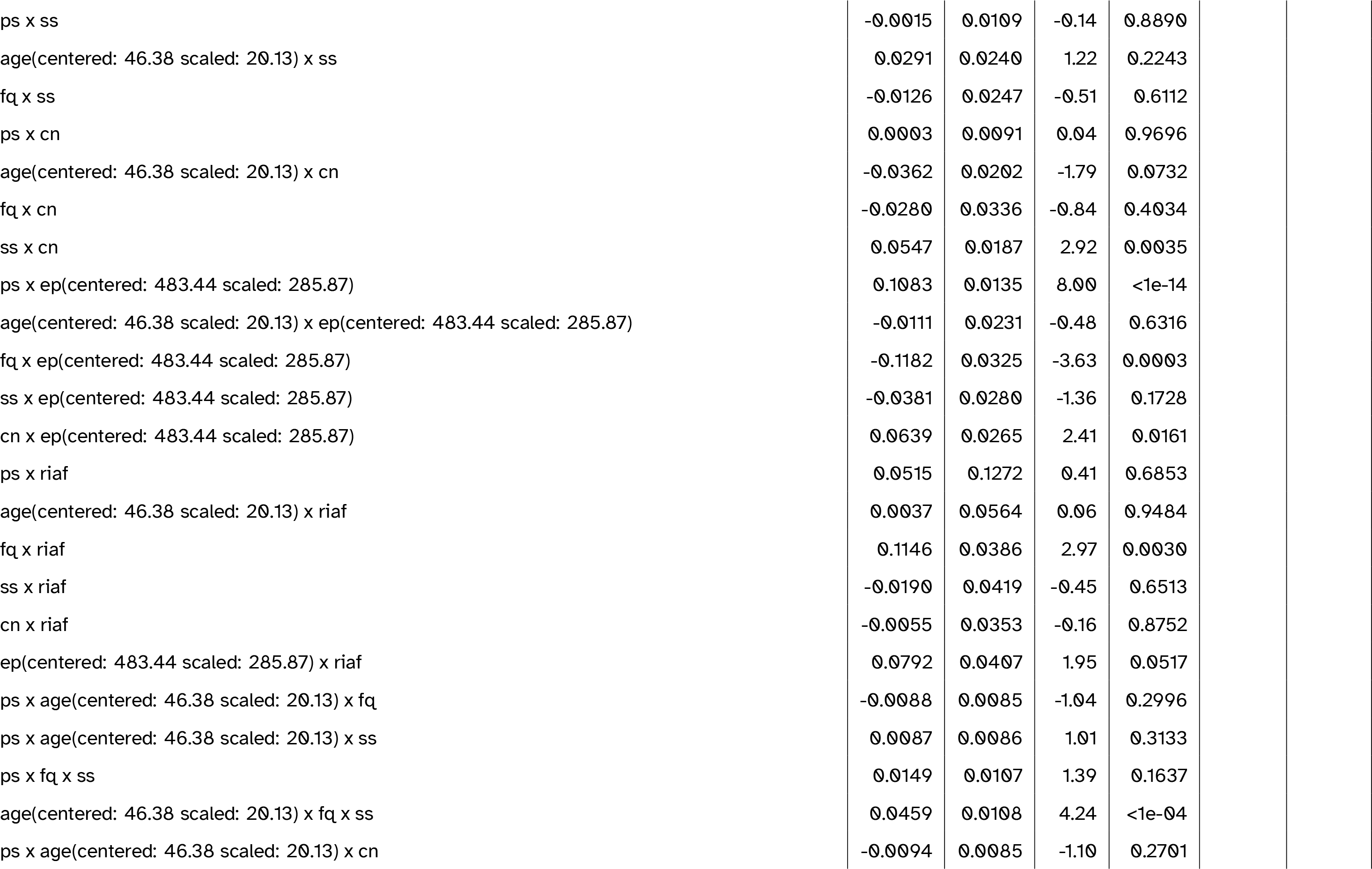
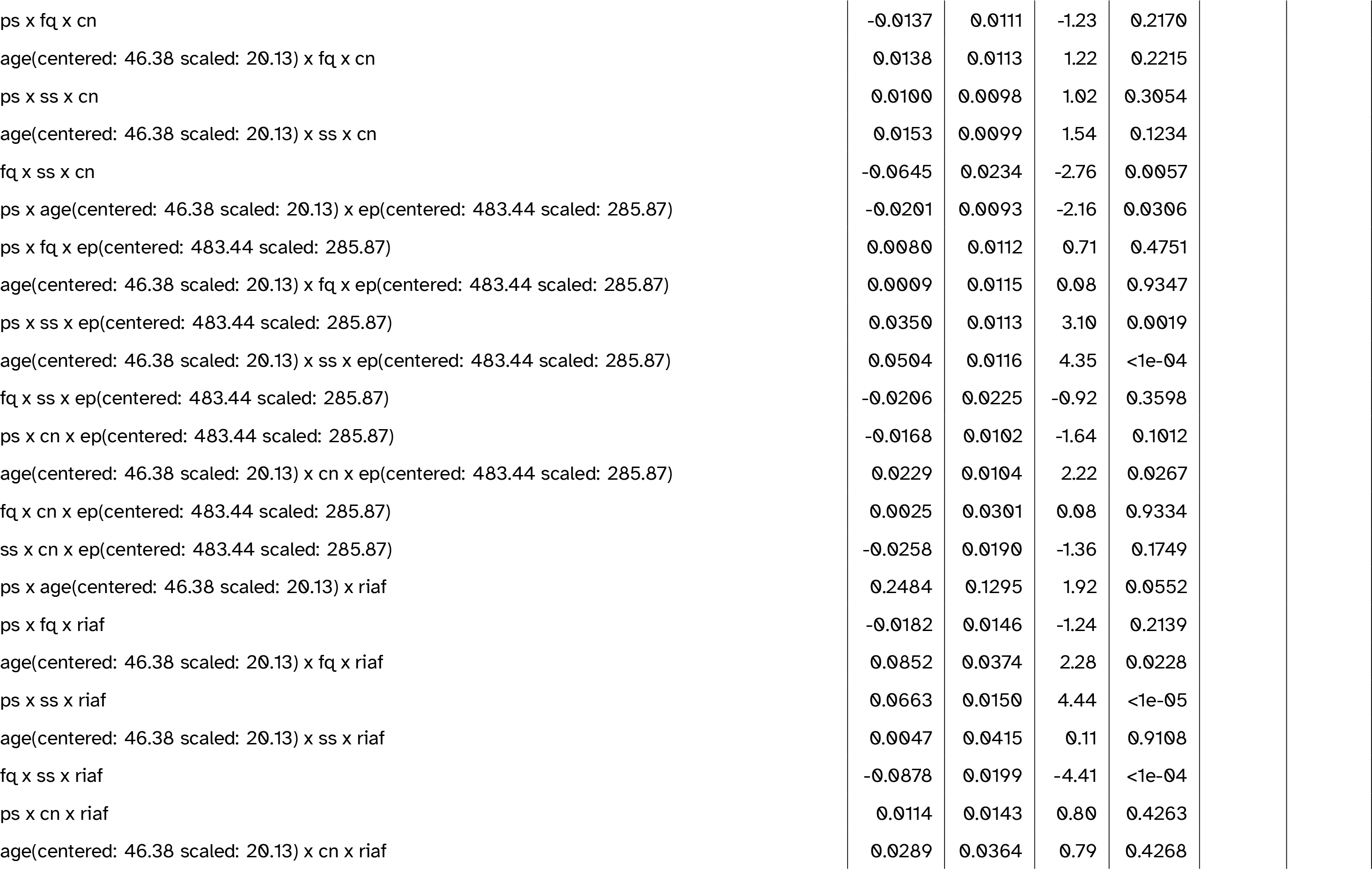
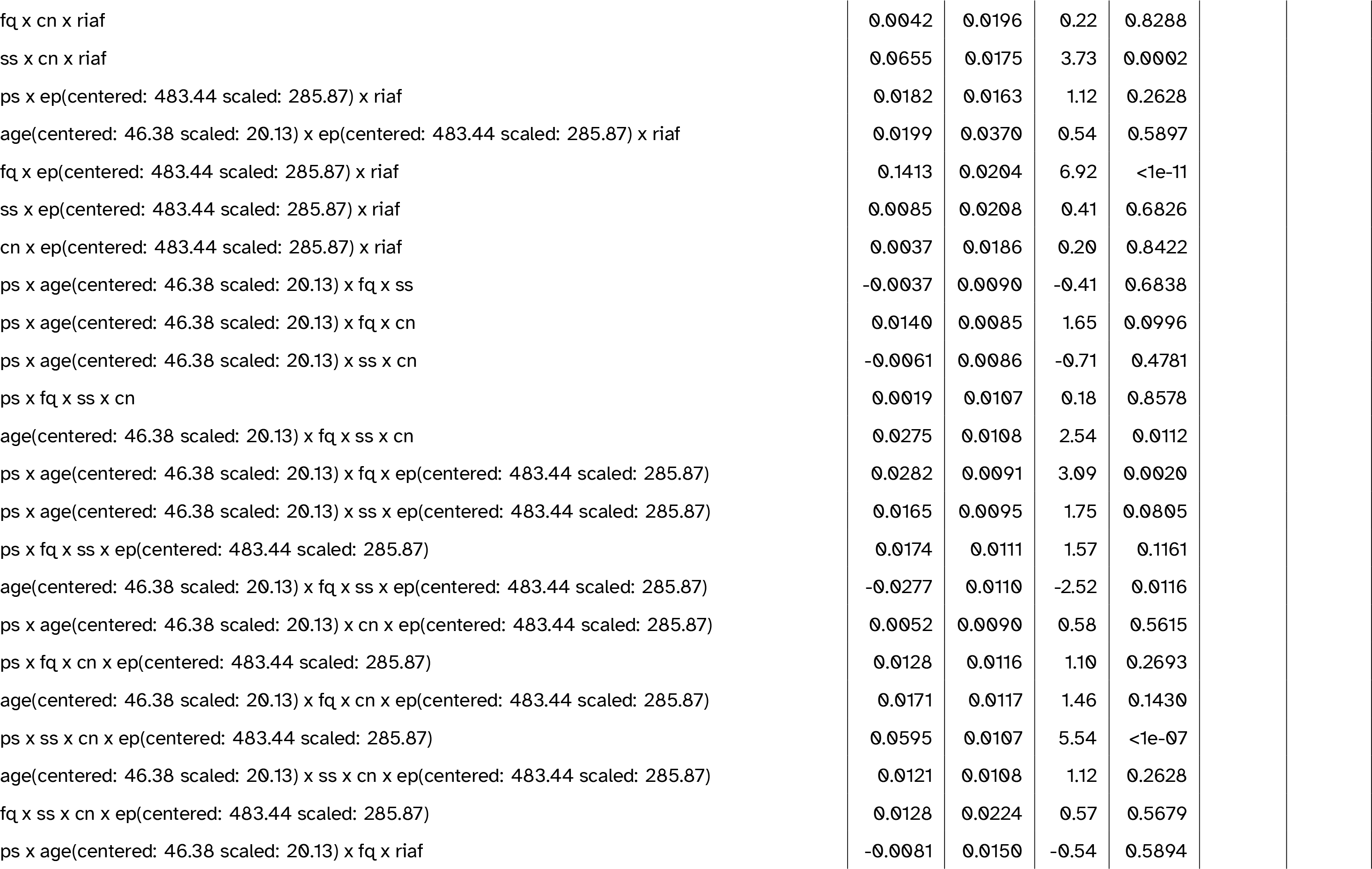
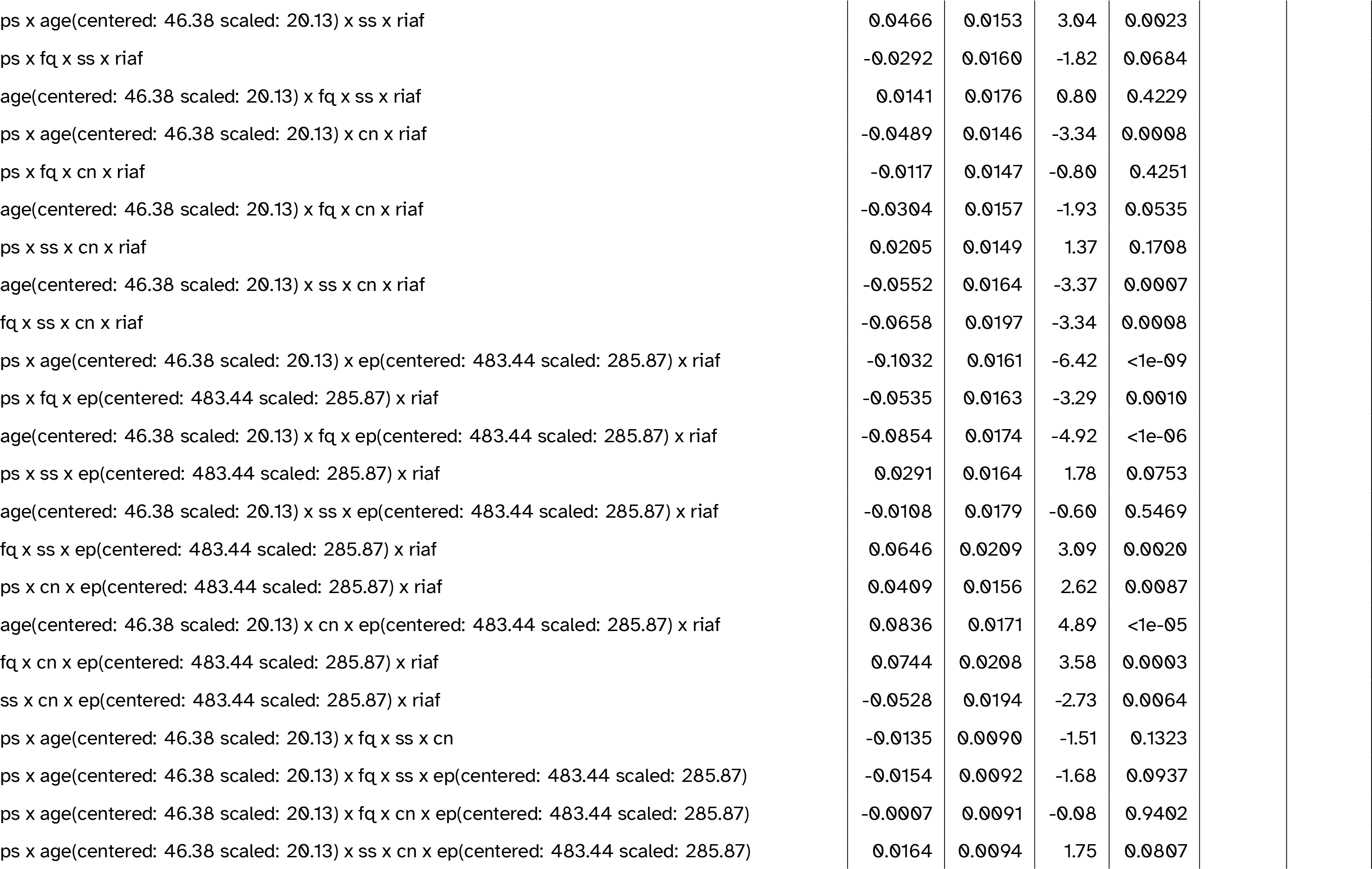
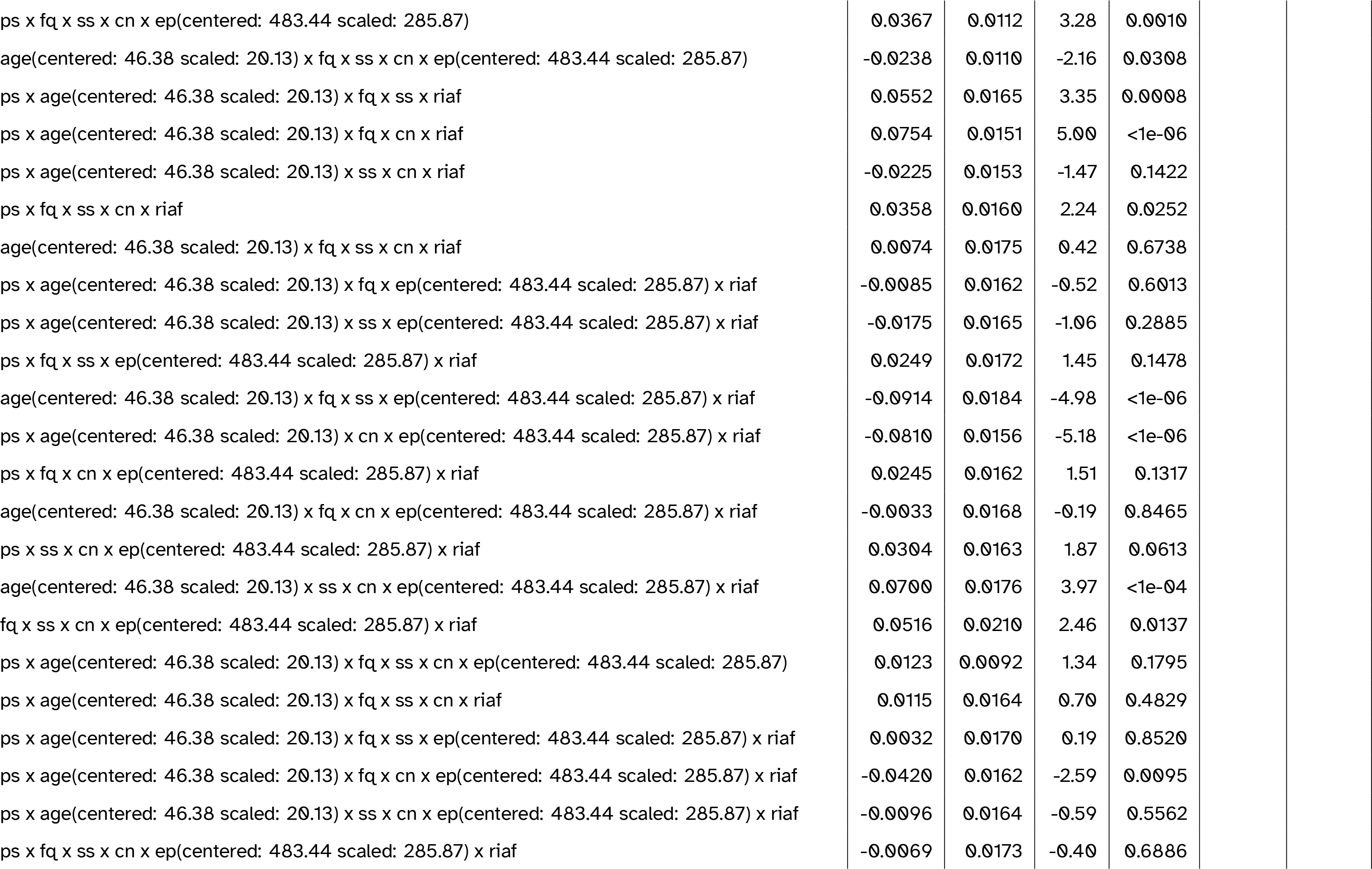
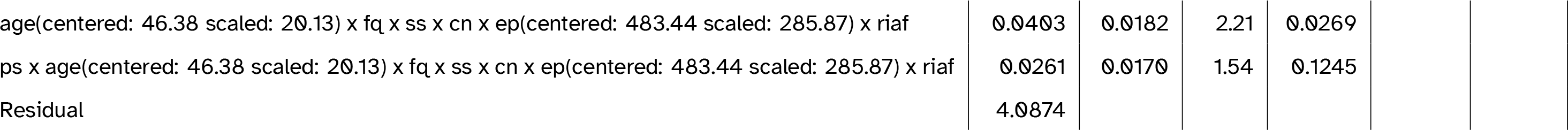
Full model summary for the best-fitting model including individual alpha frequency (IAF). Abbreviations: cn = canonicity; ep = epoch; fq = word frequency; ps = prestimulus amplitude; ps2 = quadratic prestimulus amplitude; riaf = IAF (residualised on age); ss = speaker-based surprisal; ss2 = quadratic speaker-based surprisal; ss3 = cubic speaker-based surprisal

**Table 7:**
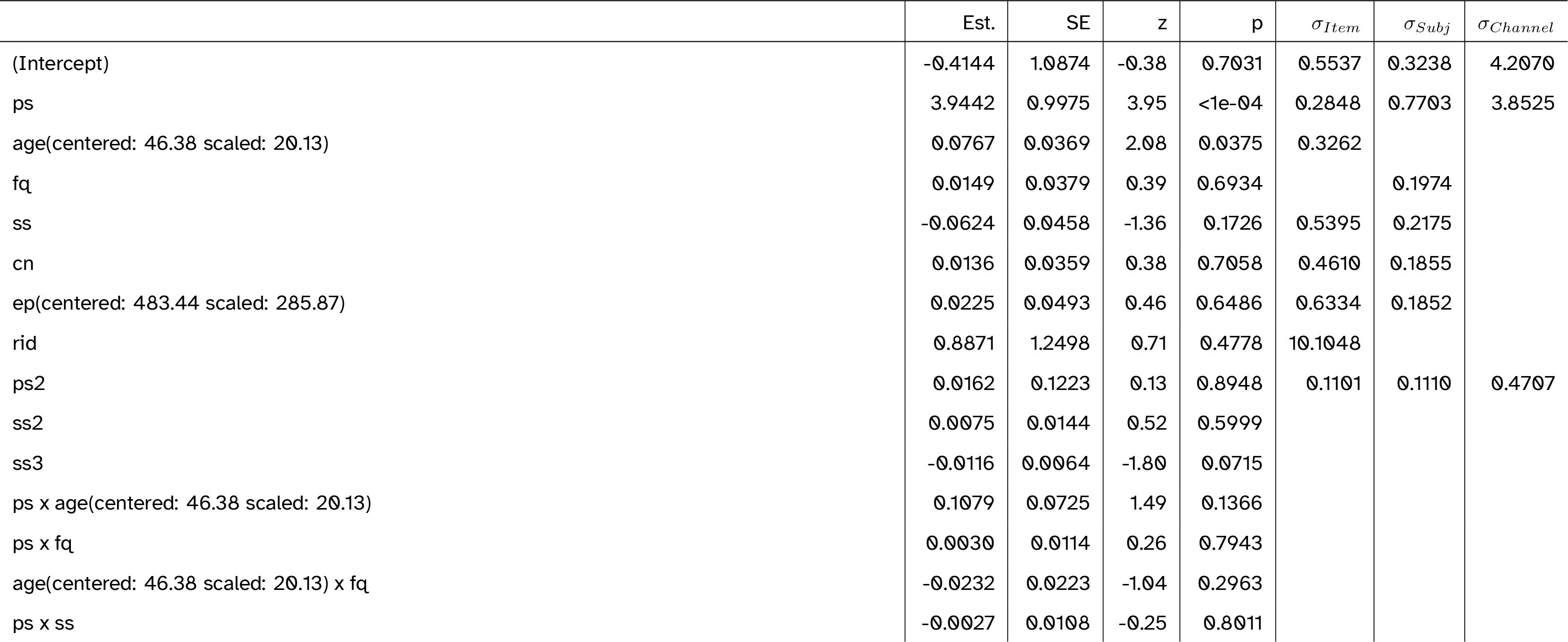
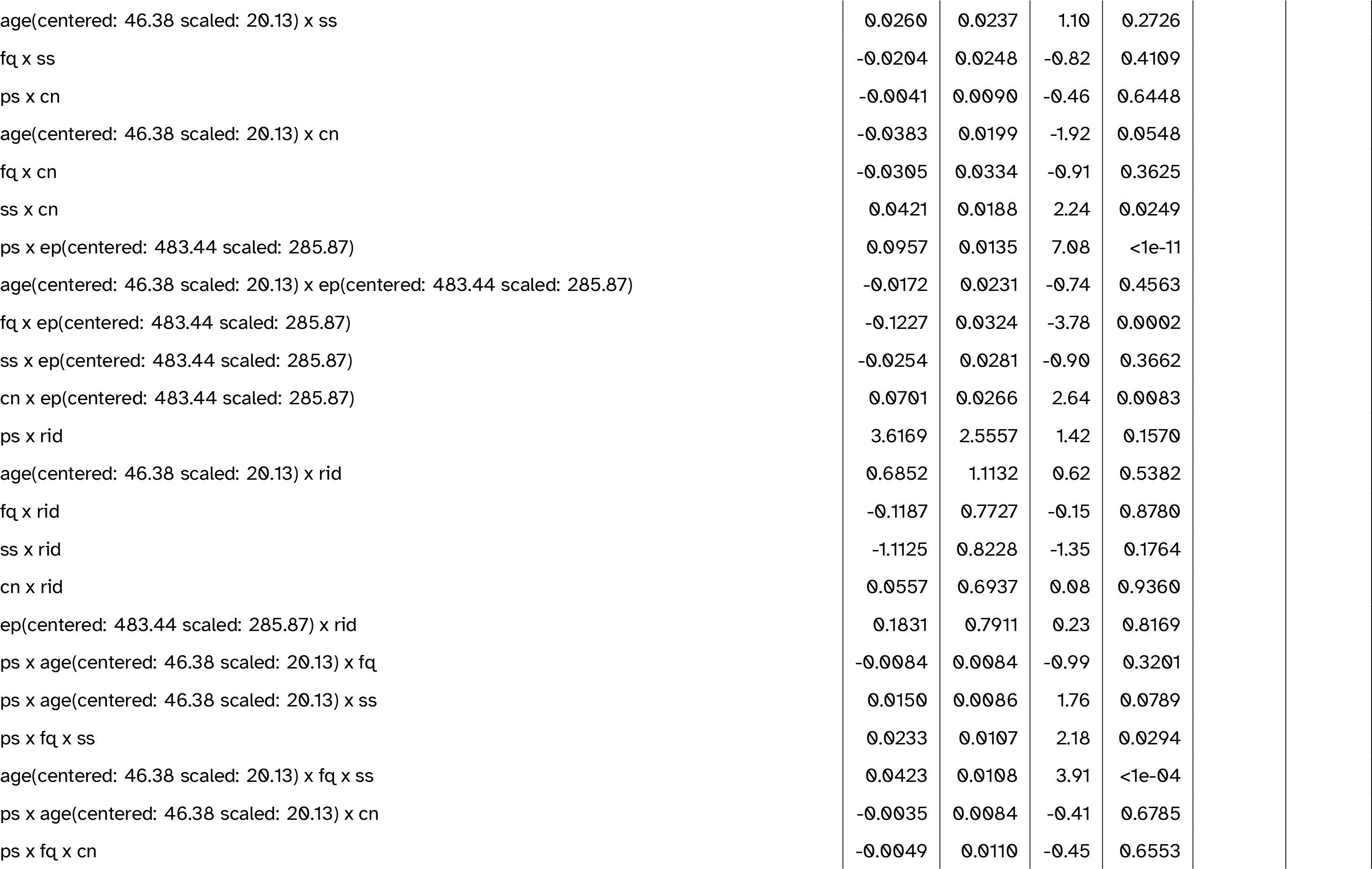
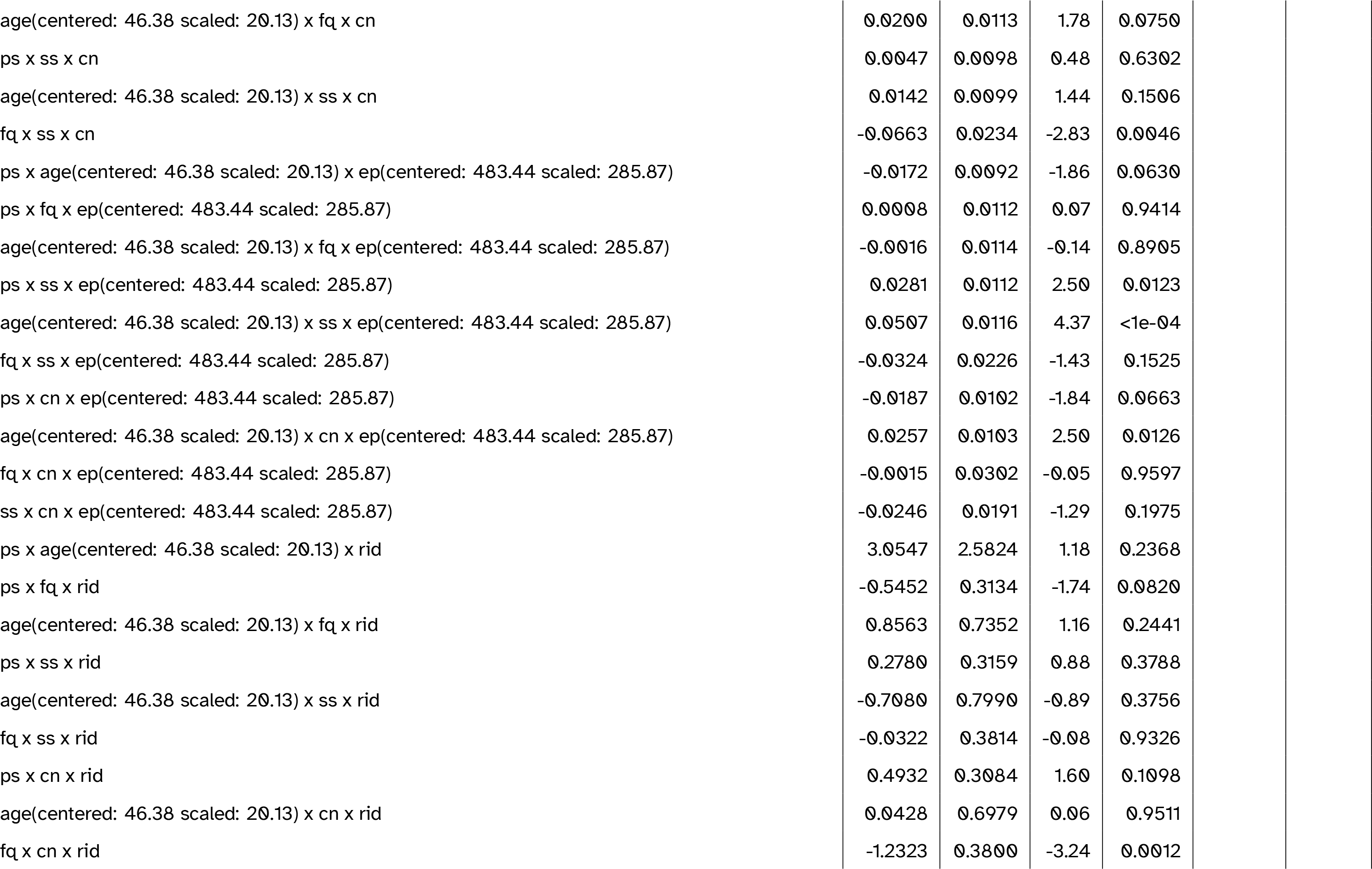
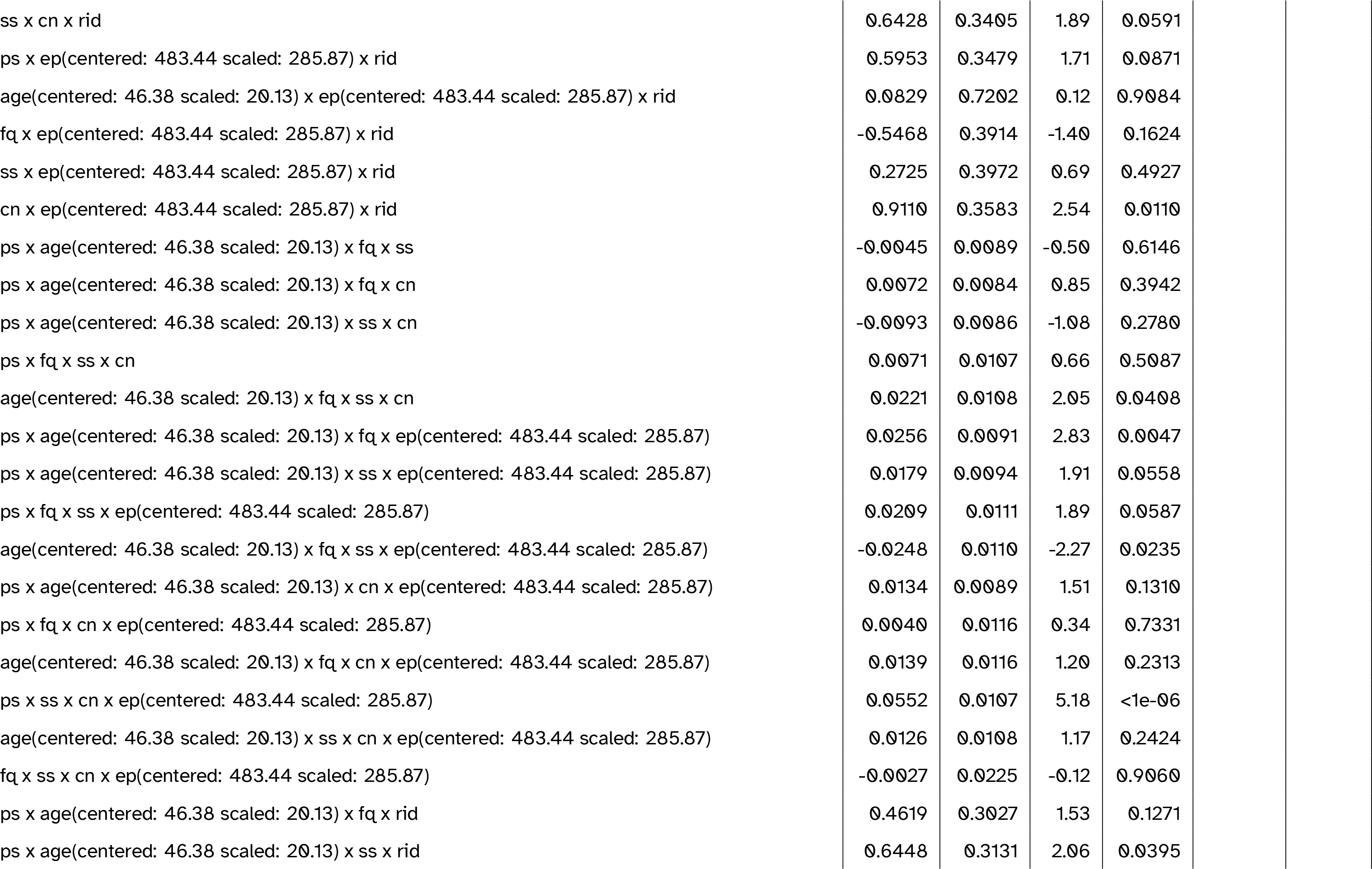
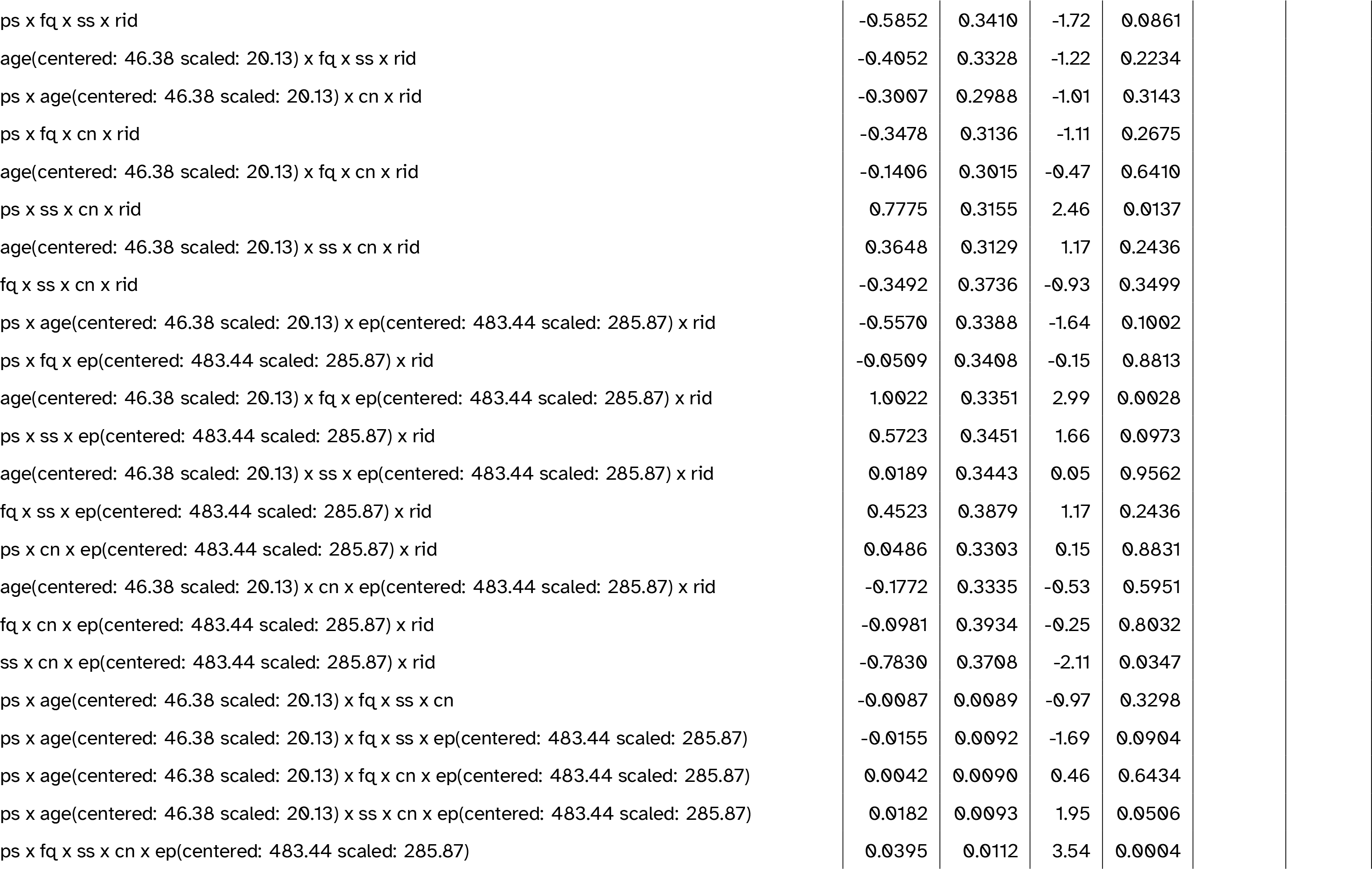
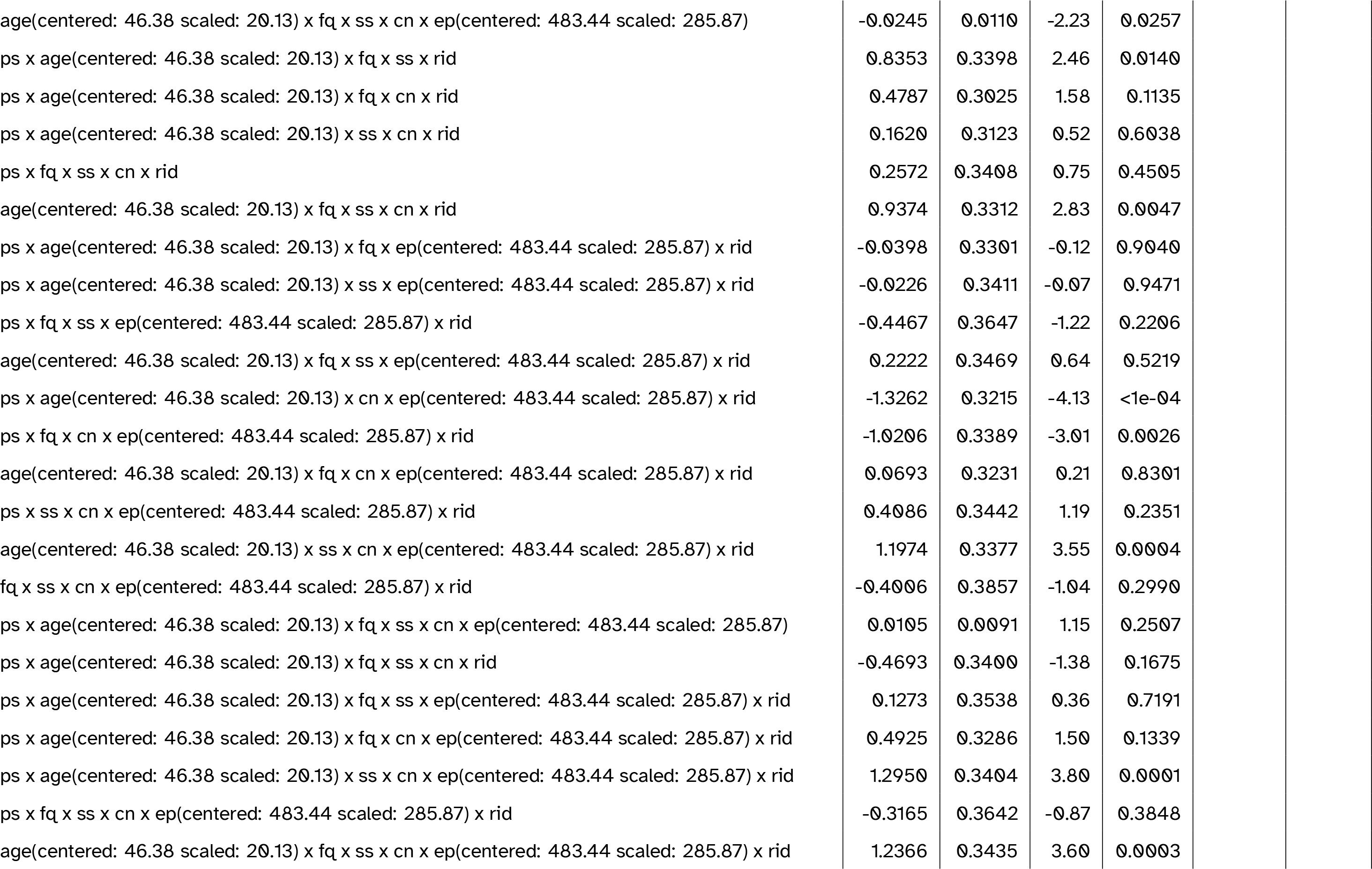
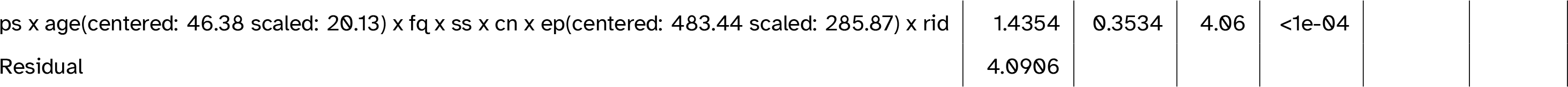
Full model summary for the best-fitting model including idea density (ID). Abbreviations: cn = canonicity; ep = epoch; fq = word frequency; ps = prestimulus amplitude; ps2 = quadratic prestimulus amplitude; rid = ID (residualised on age); ss = speaker-based surprisal; ss2 = quadratic speaker-based surprisal; ss3 = cubic speaker-based surprisal

**Table 8:**
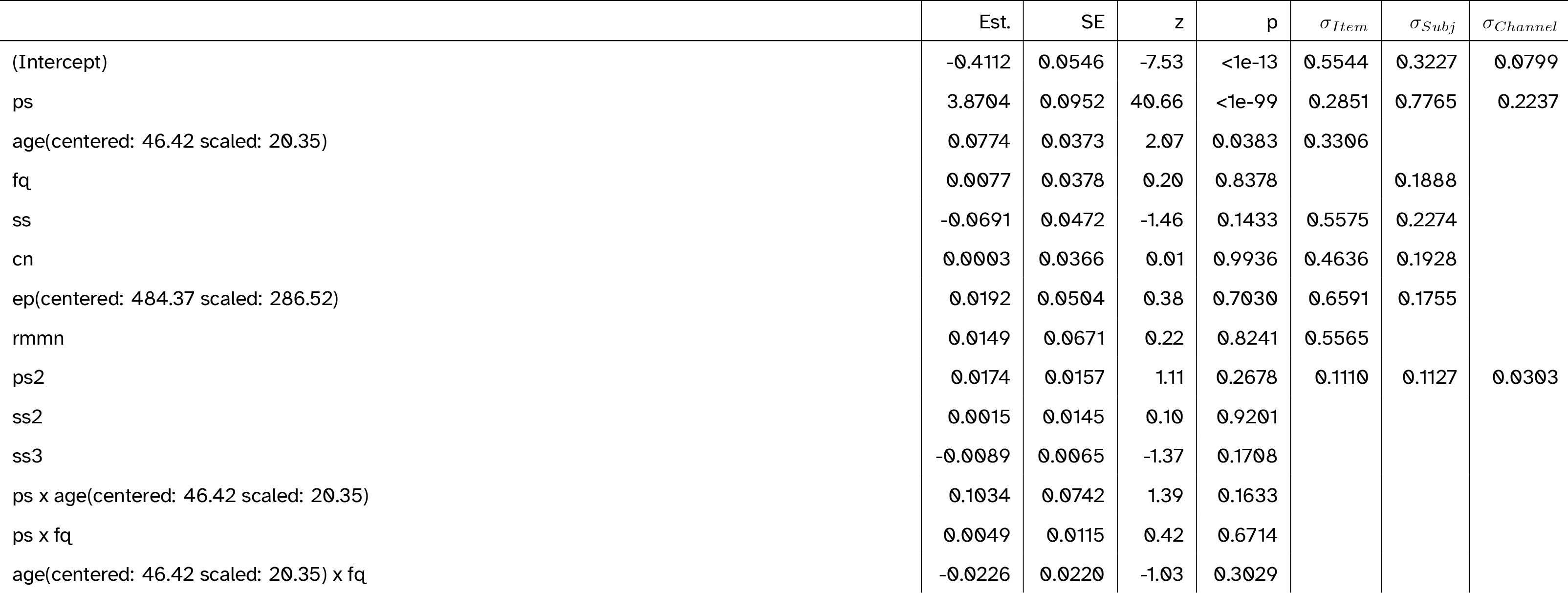
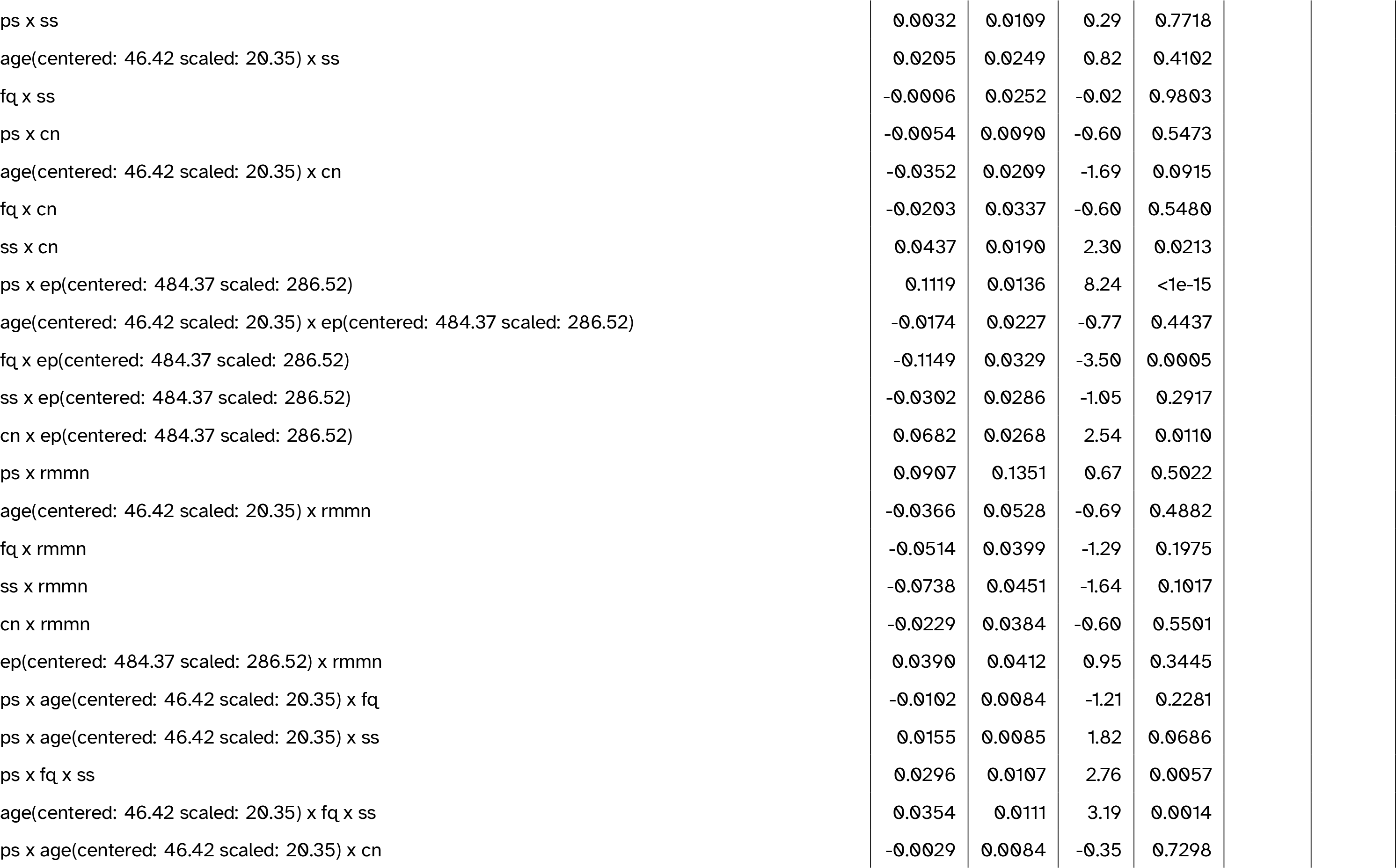
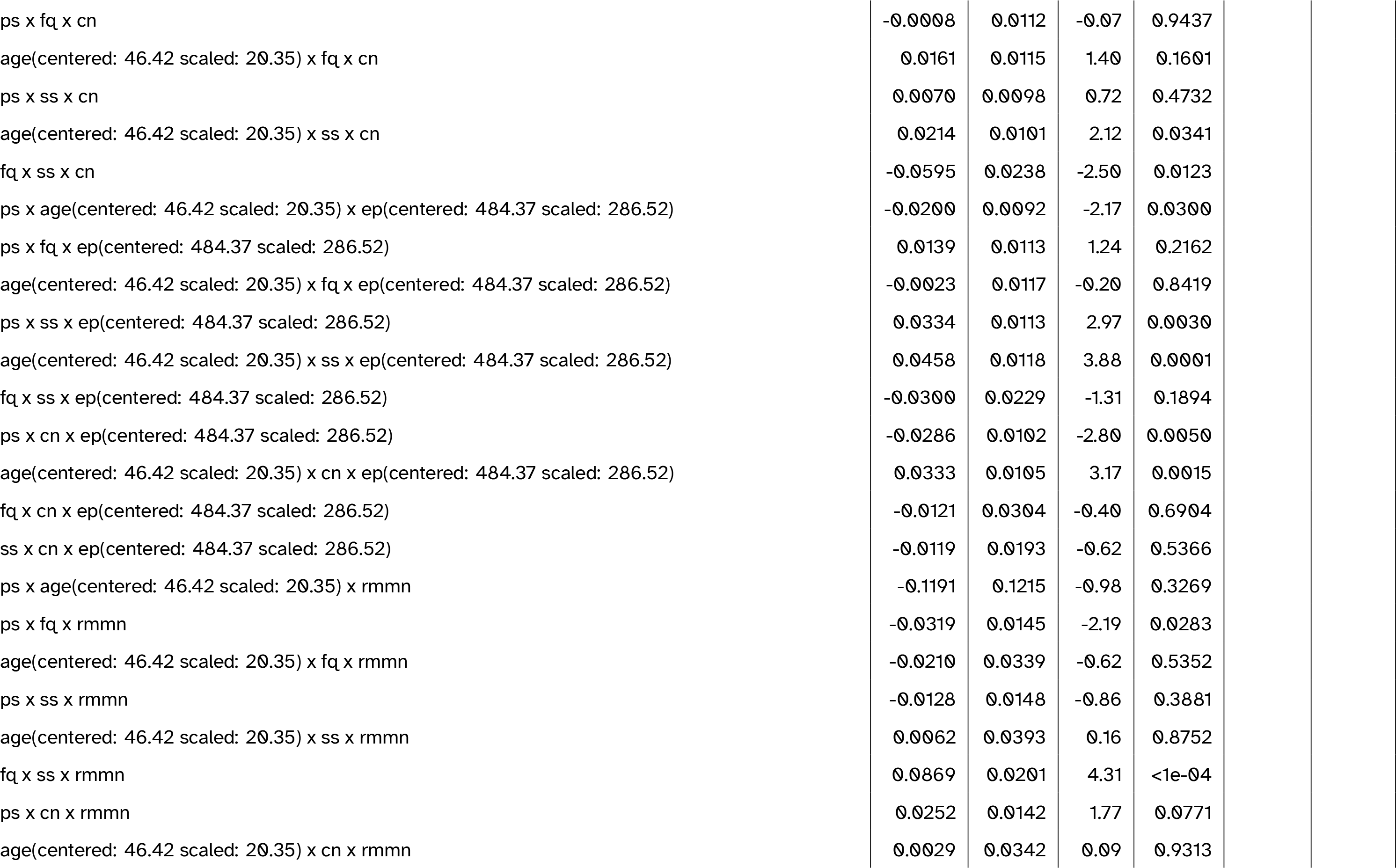
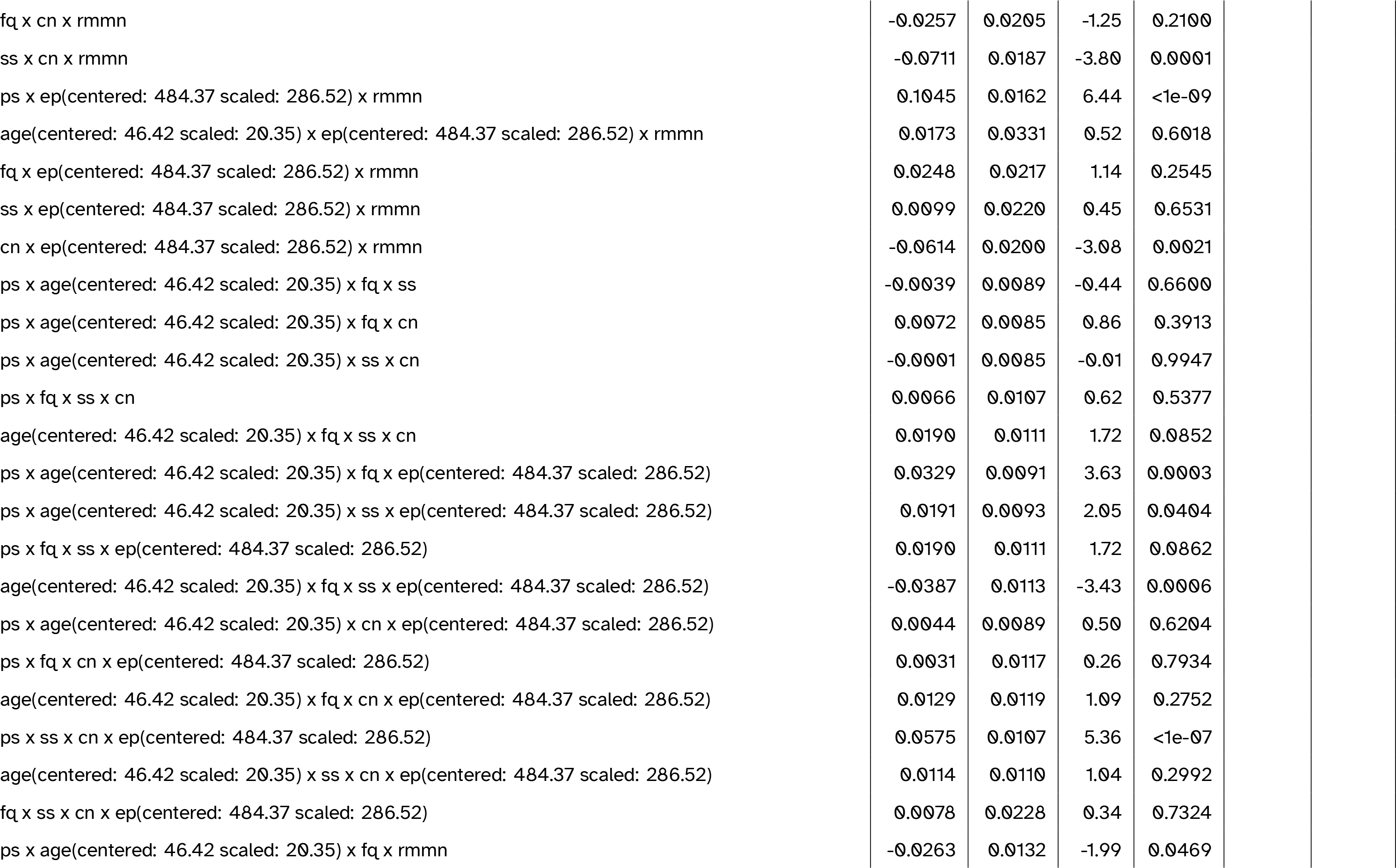
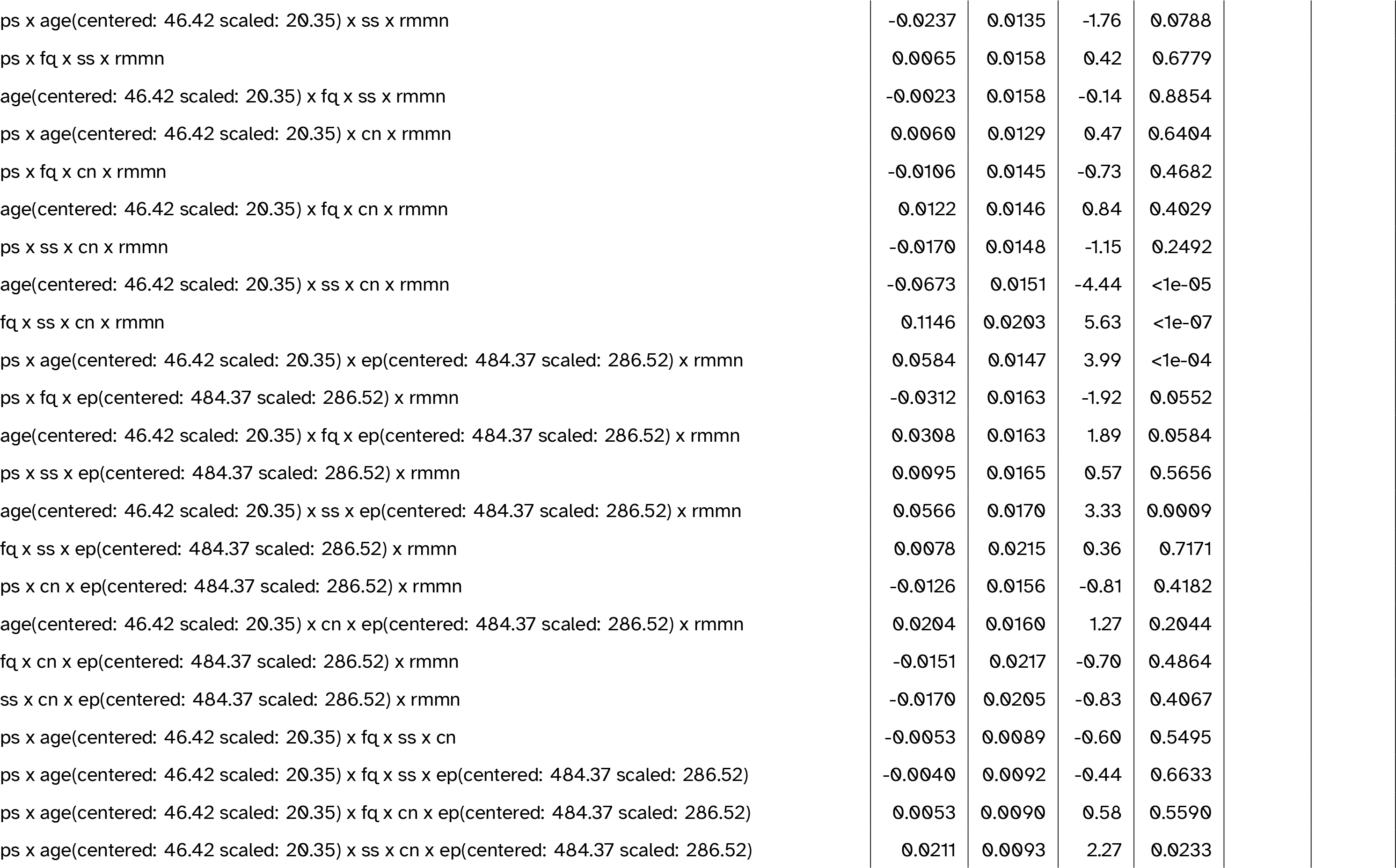
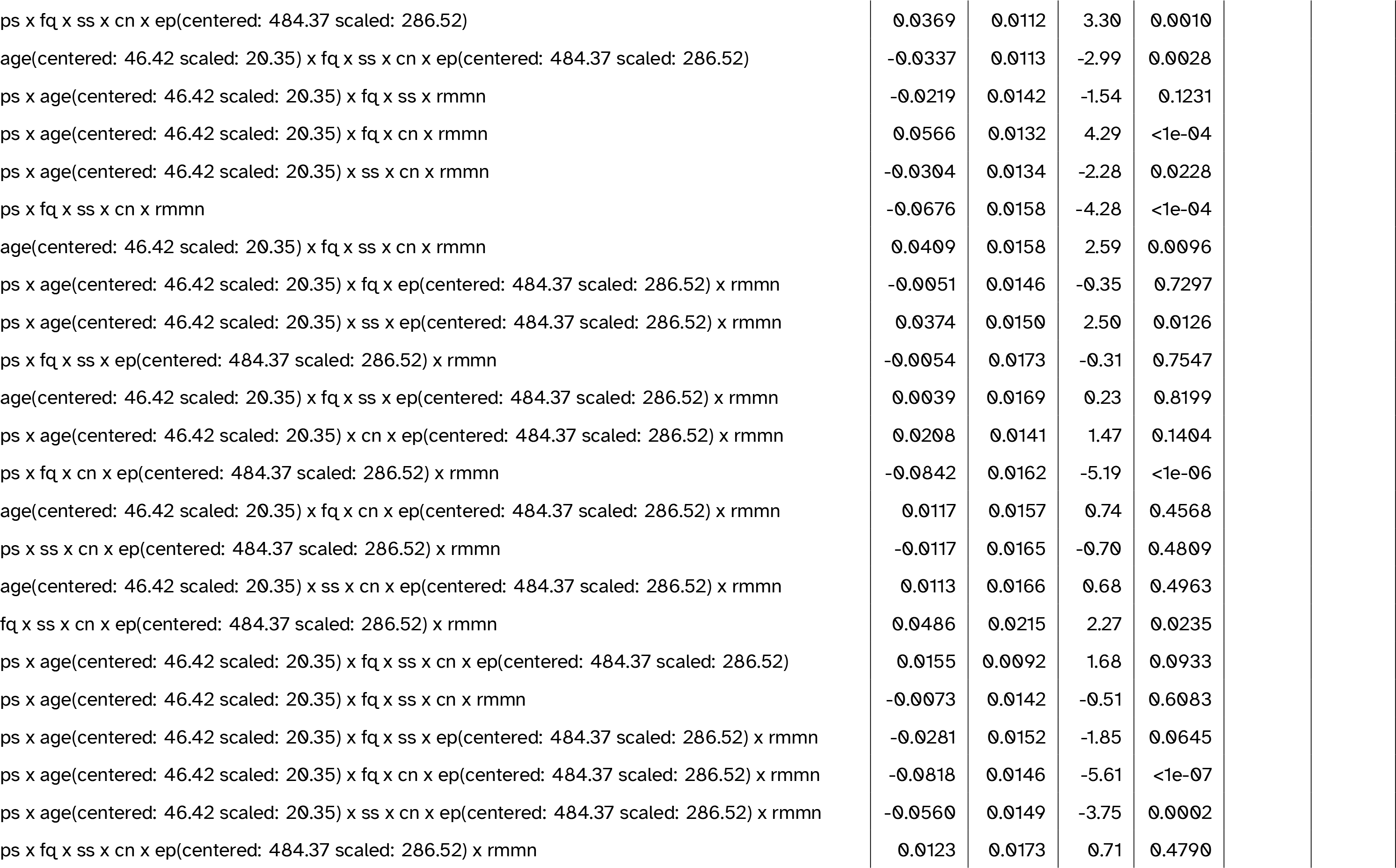
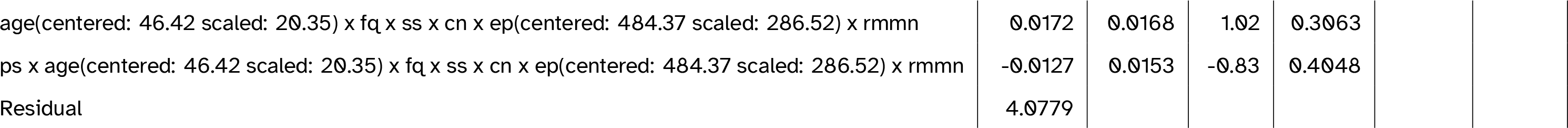
Full model summary for the best-fitting model including MMN amplitude. Abbreviations: cn = canonicity; ep = epoch; fq = word frequency; ps = prestimulus amplitude; ps2 = quadratic prestimulus amplitude; rmmn = MMN amplitude (residualised on age); ss = speaker-based surprisal; ss2 = quadratic speaker-based surprisal; ss3 = cubic speaker-based surprisal

**Table 9:**
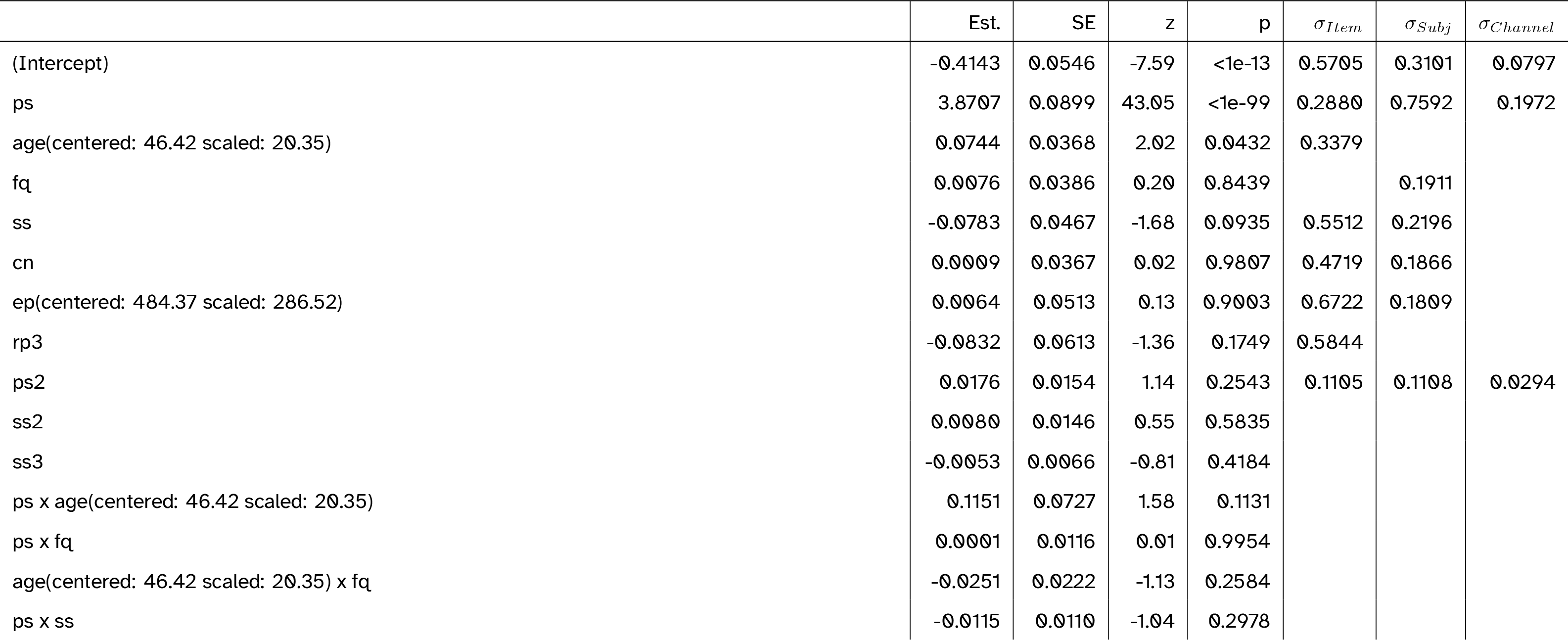
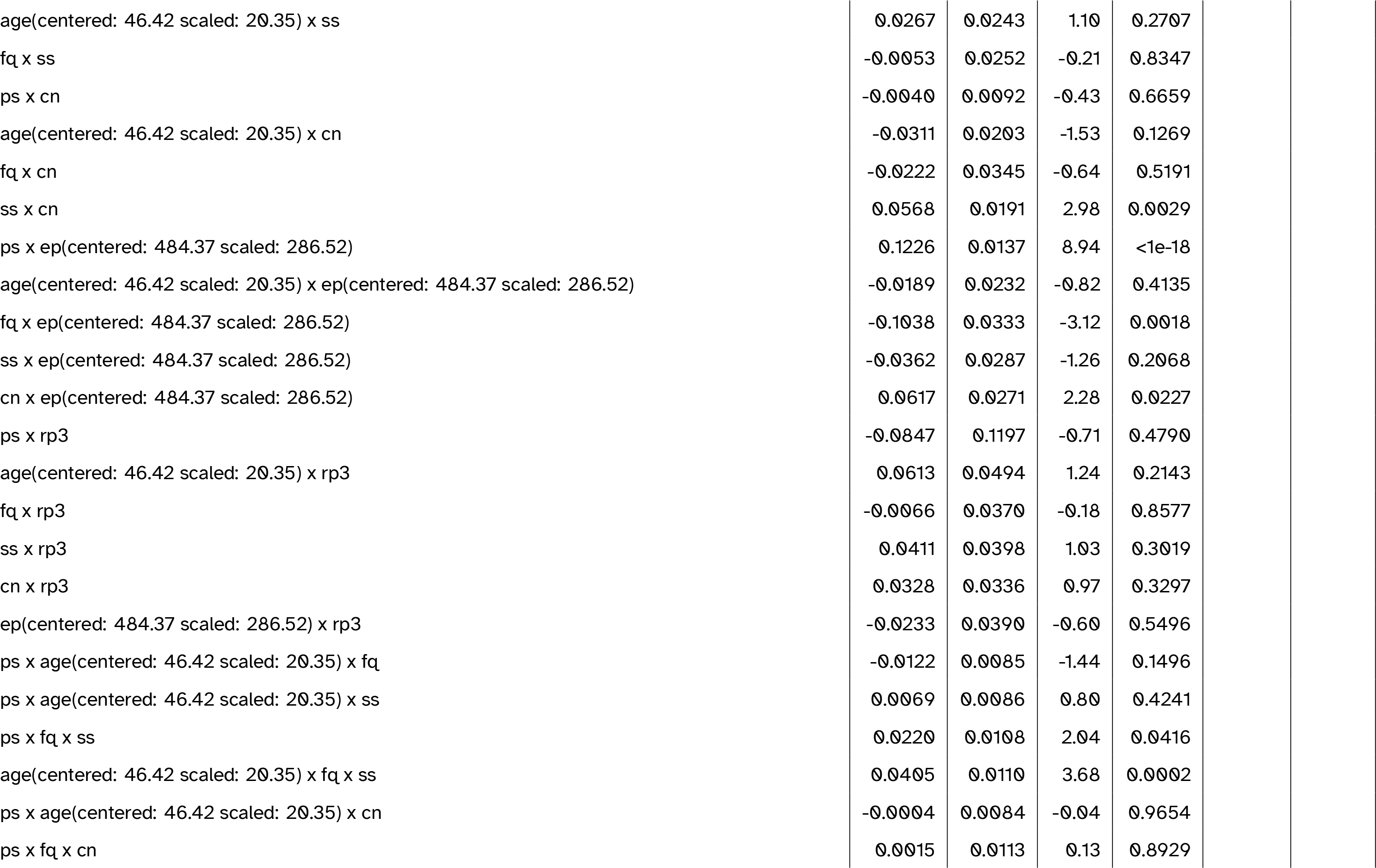
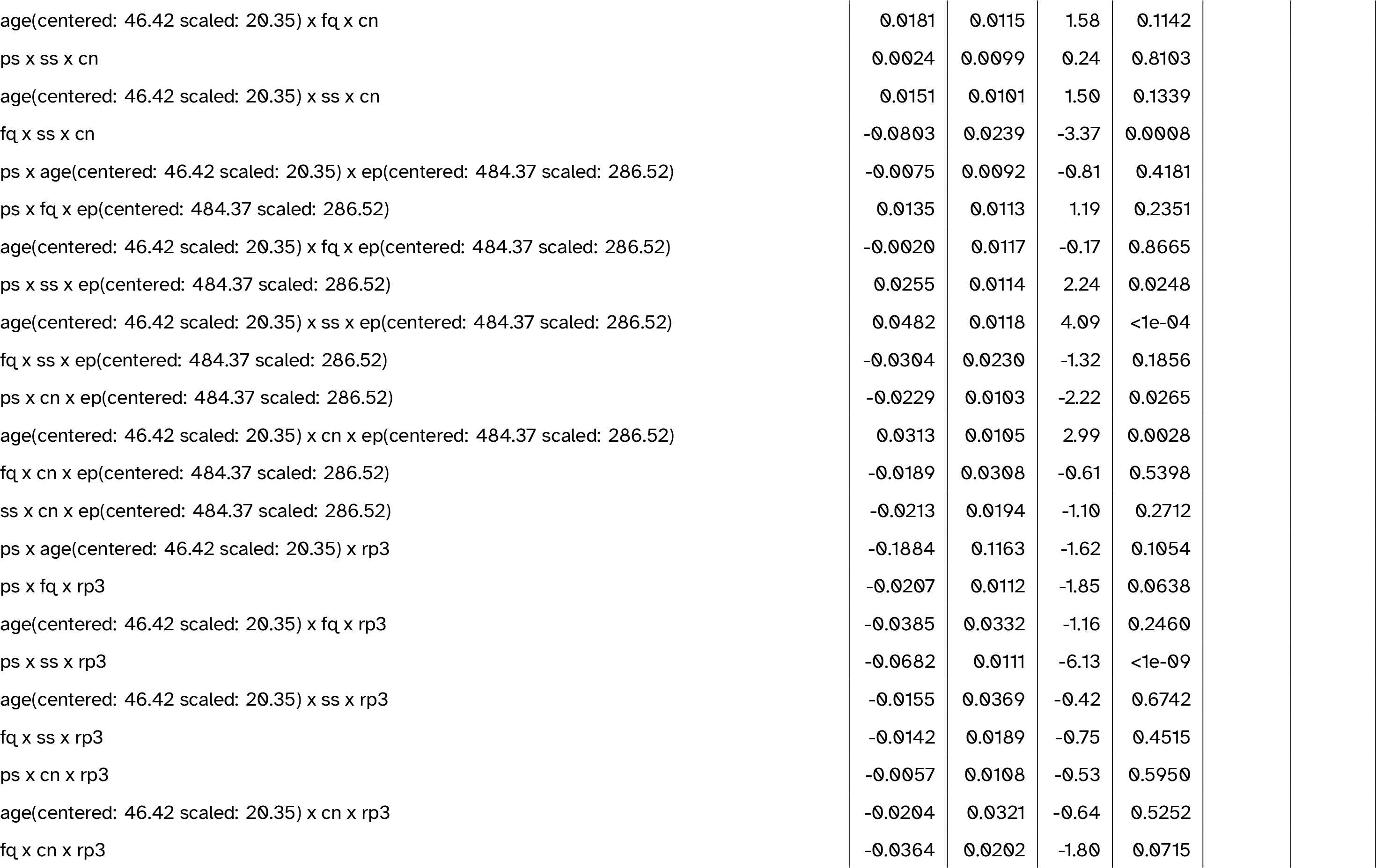
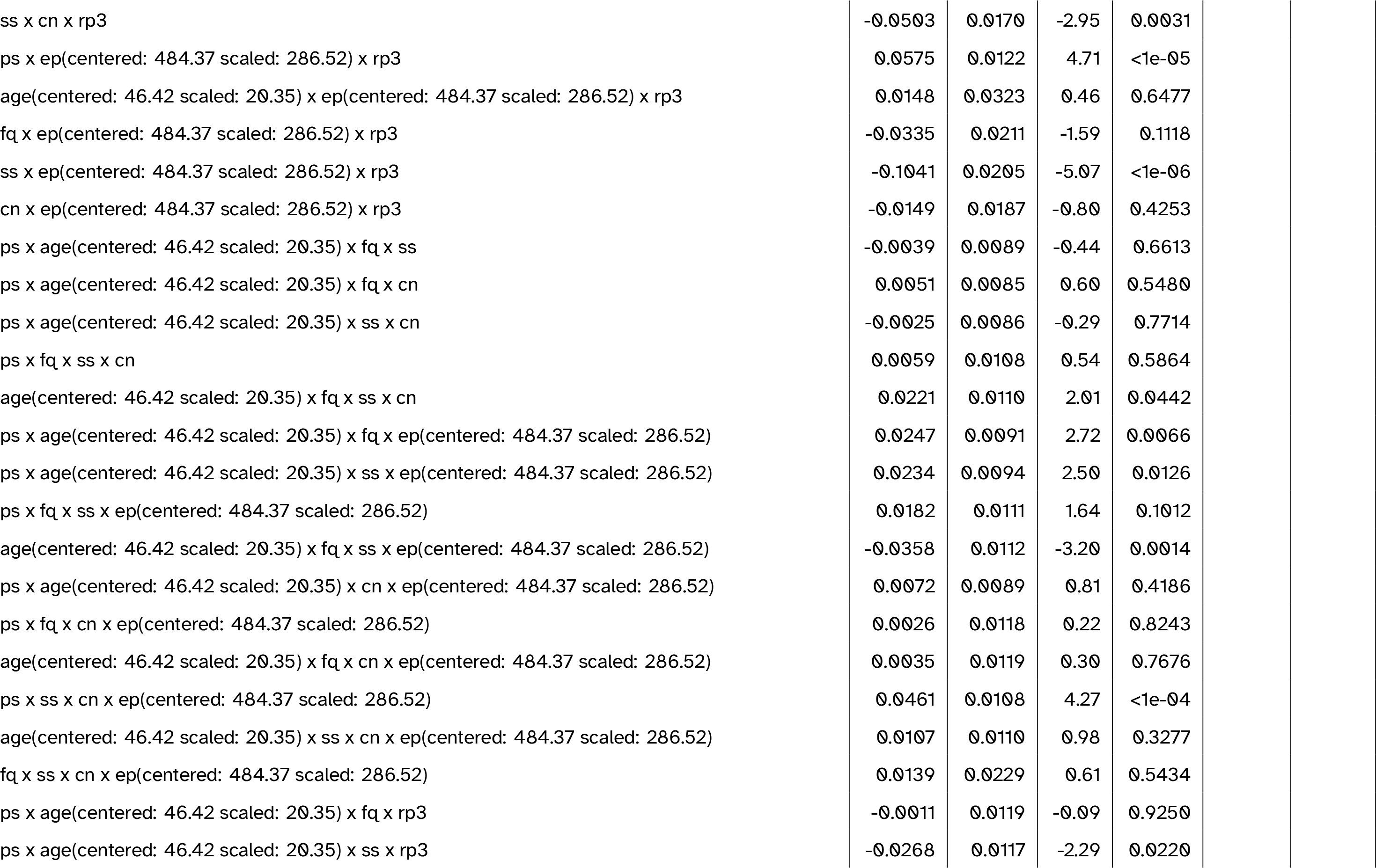
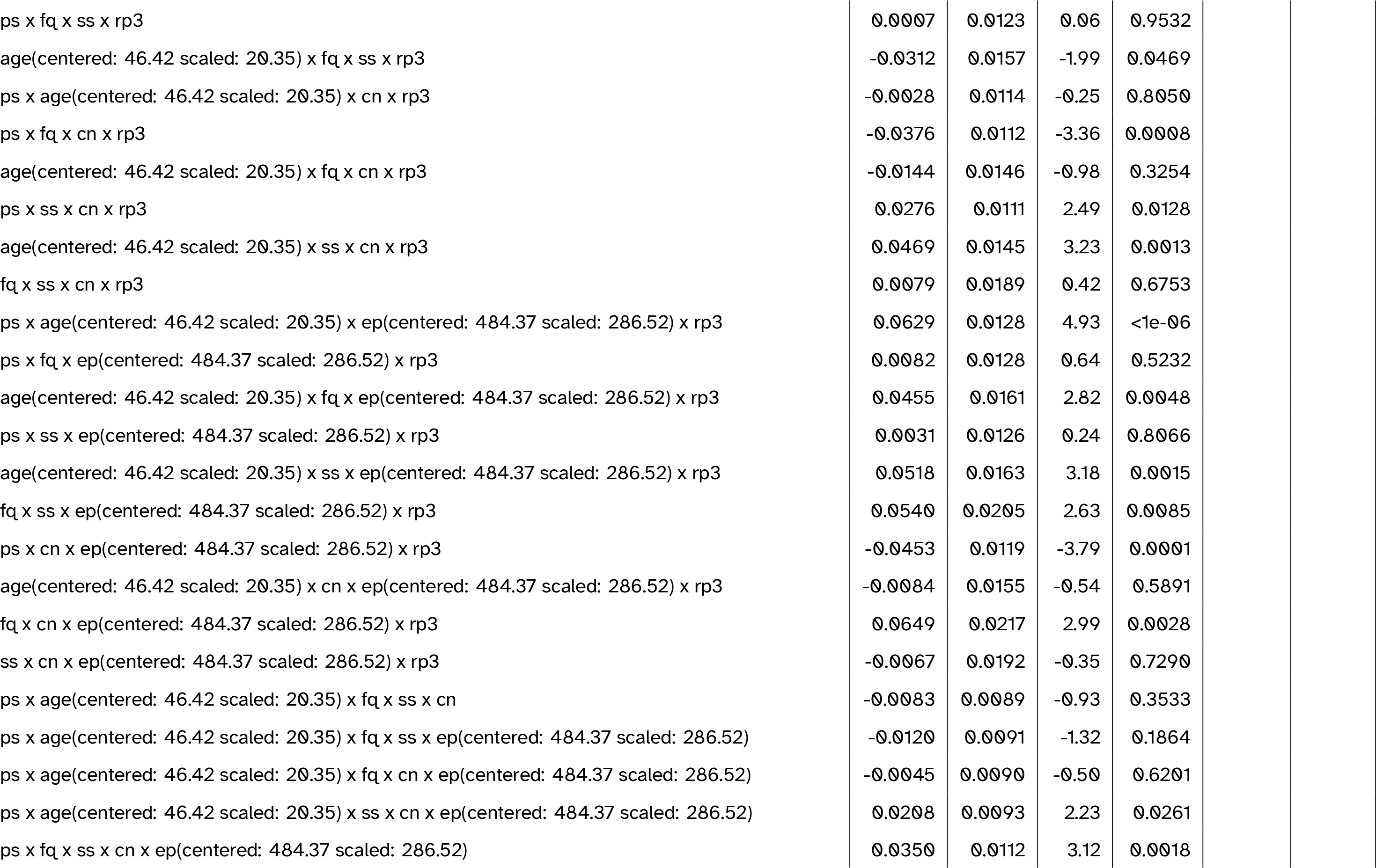
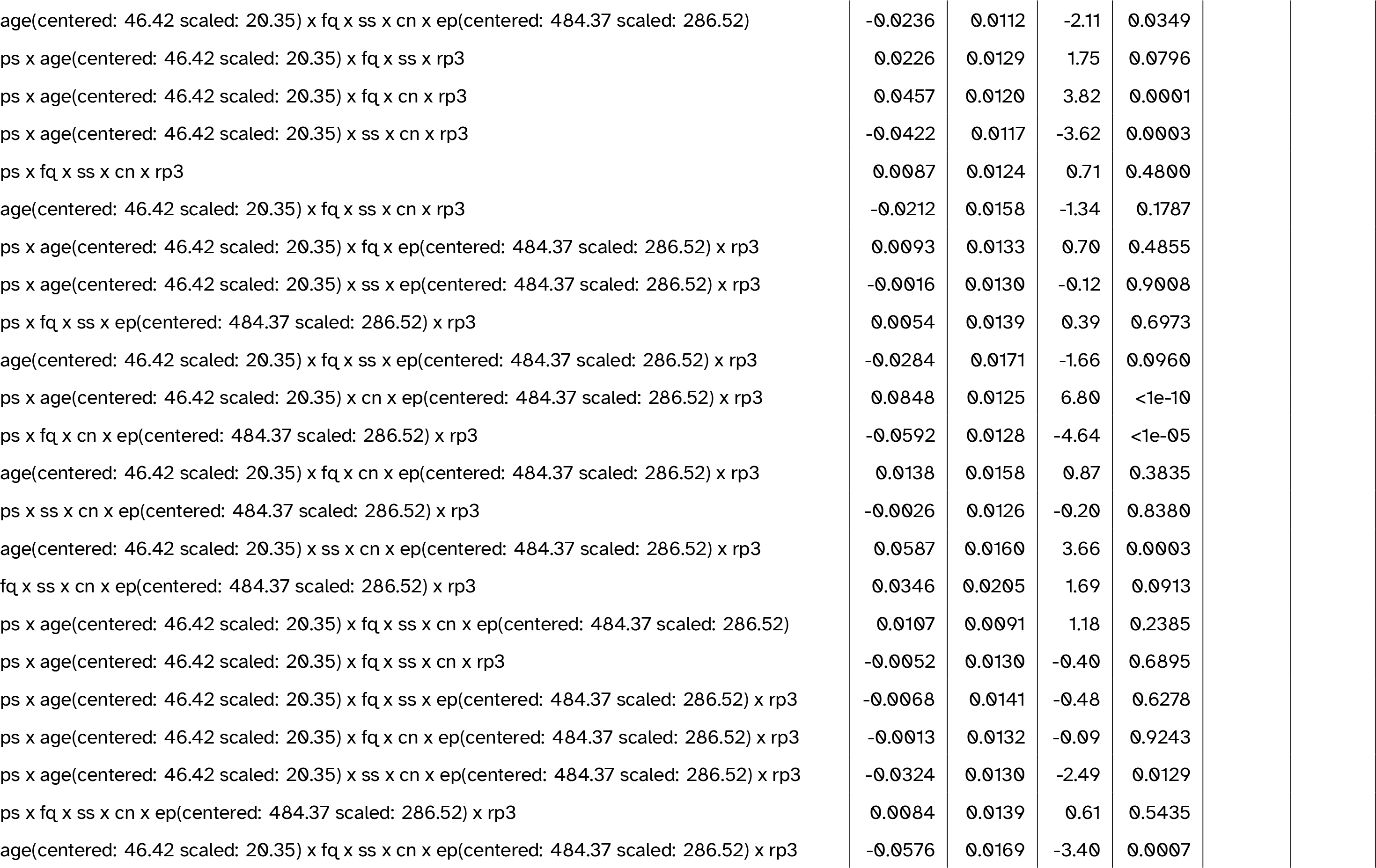
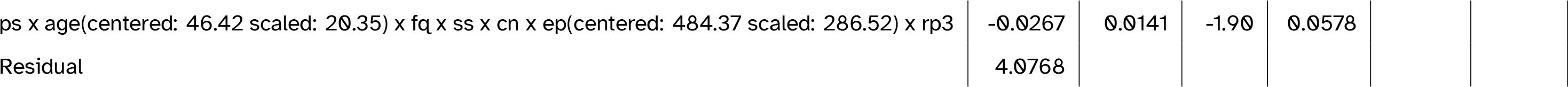
Full model summary for the best-fitting model including P3 amplitude. Abbreviations: cn = canonicity; ep = epoch; fq = word frequency; ps = prestimulus amplitude; ps2 = quadratic prestimulus amplitude; rp3 = P3 amplitude (residualised on age); ss = speaker-based surprisal; ss2 = quadratic speaker-based surprisal; ss3 = cubic speaker-based surprisal

**Table 10:**
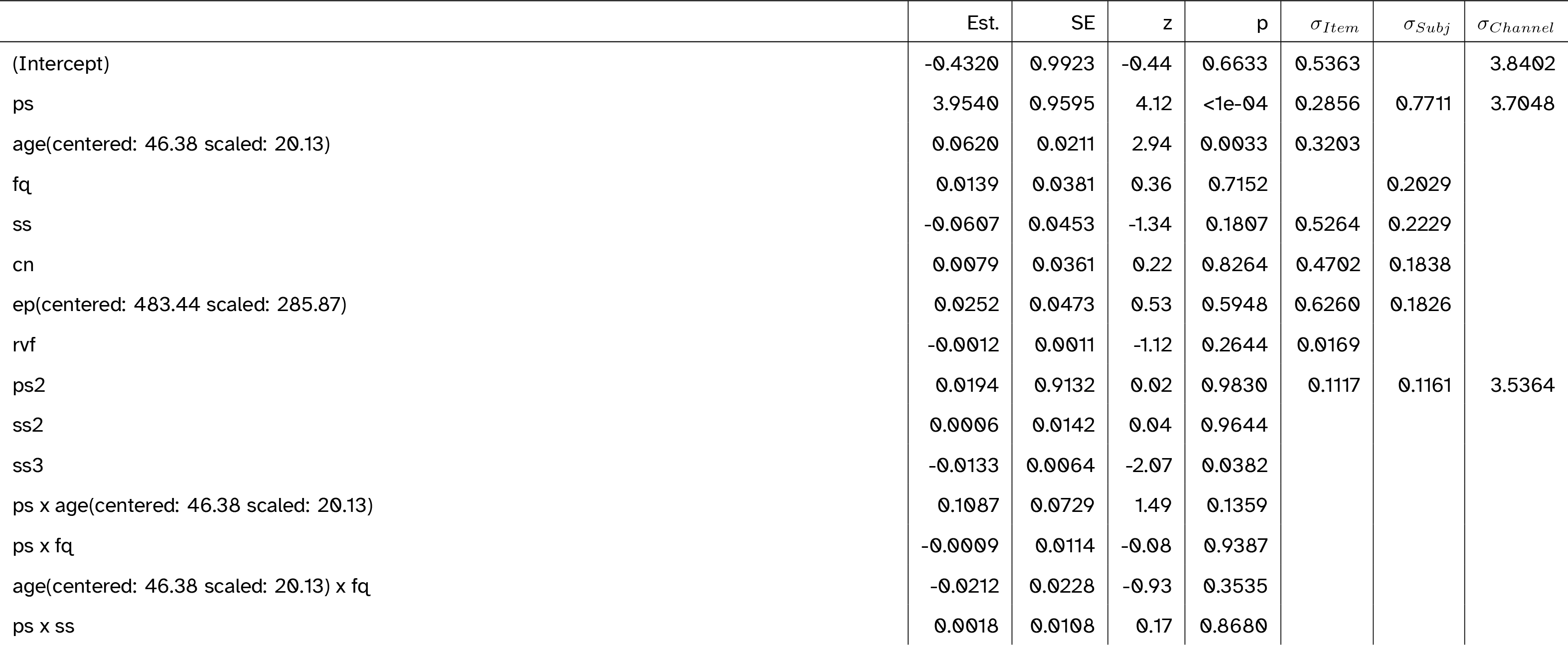
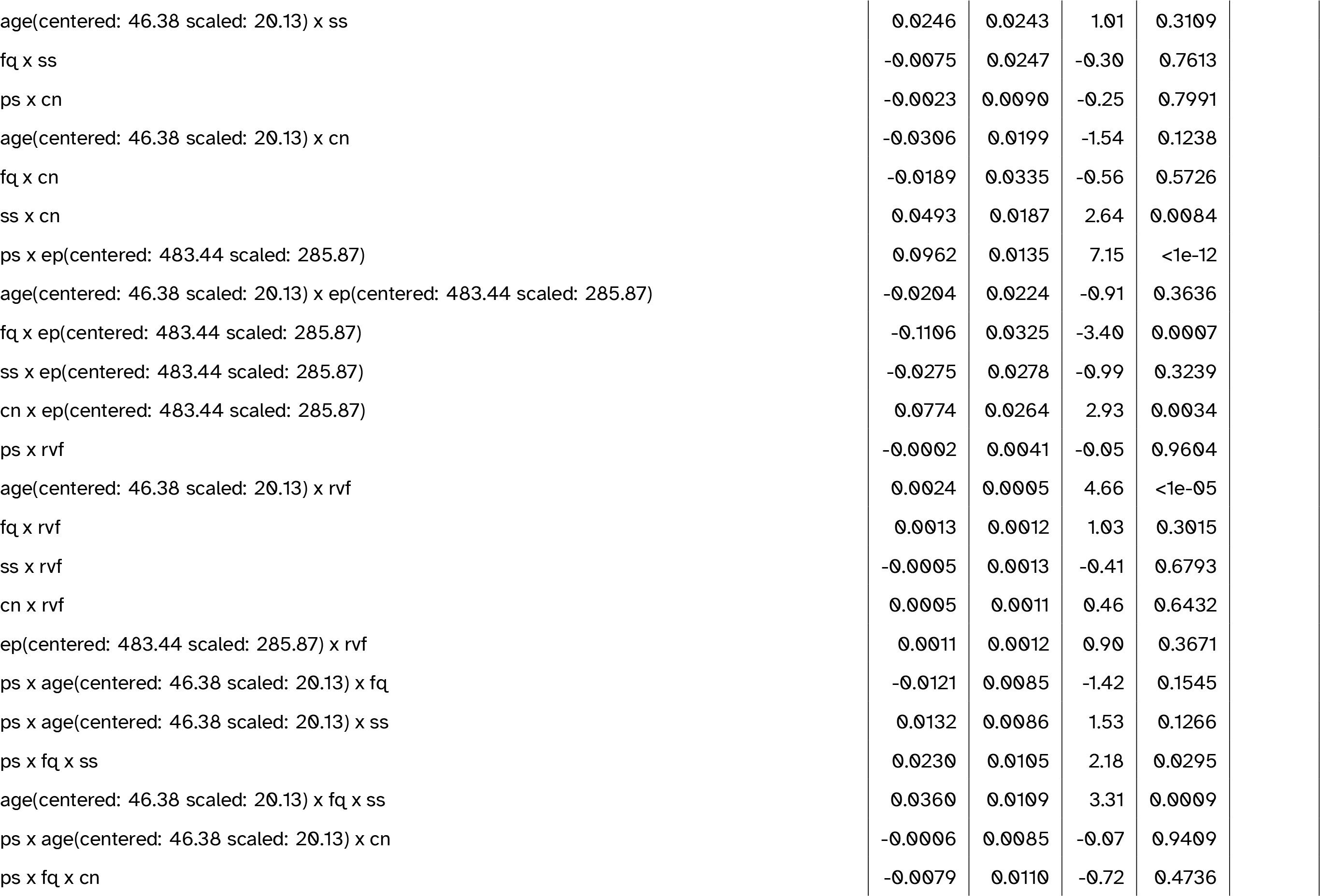
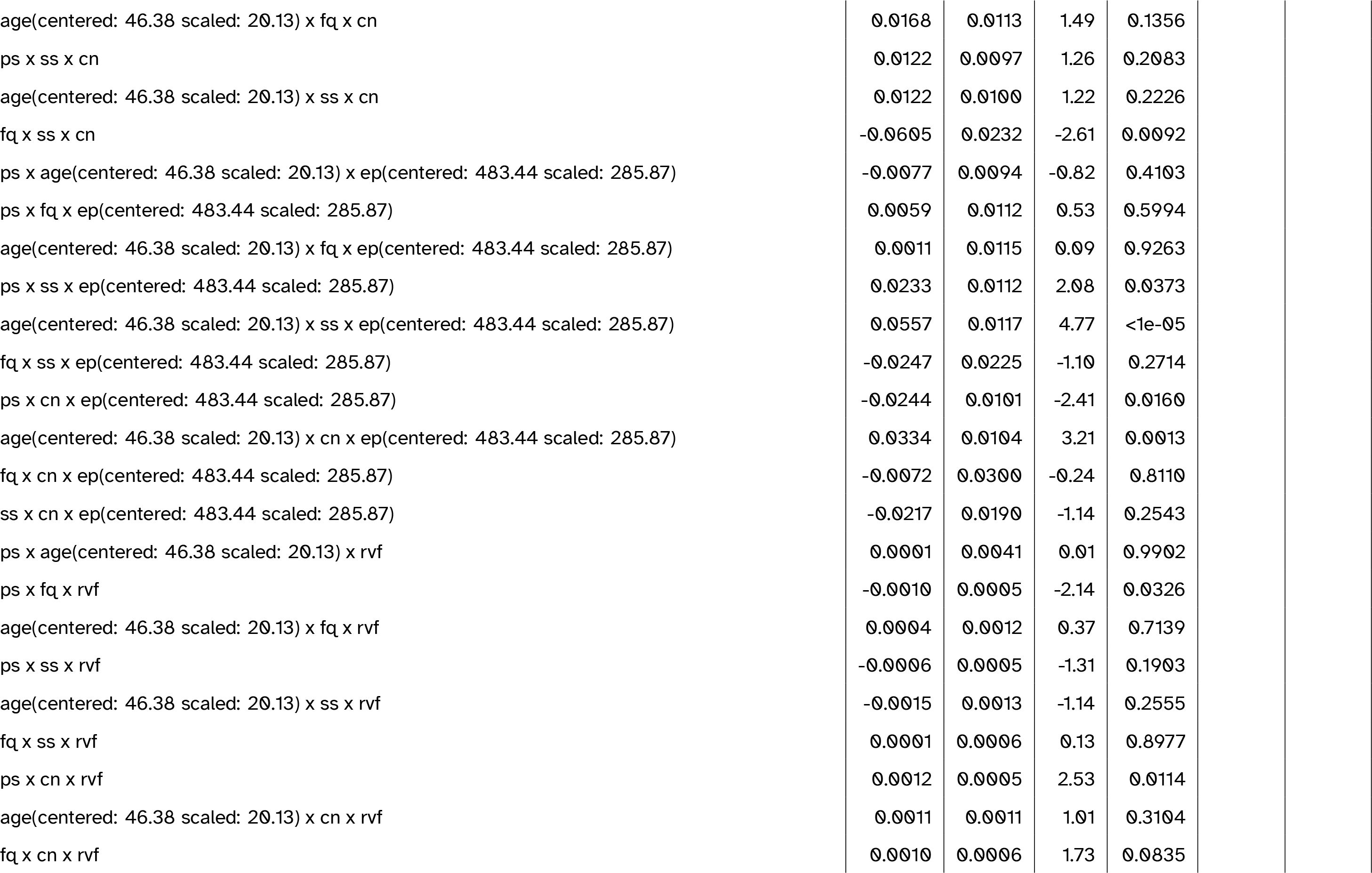
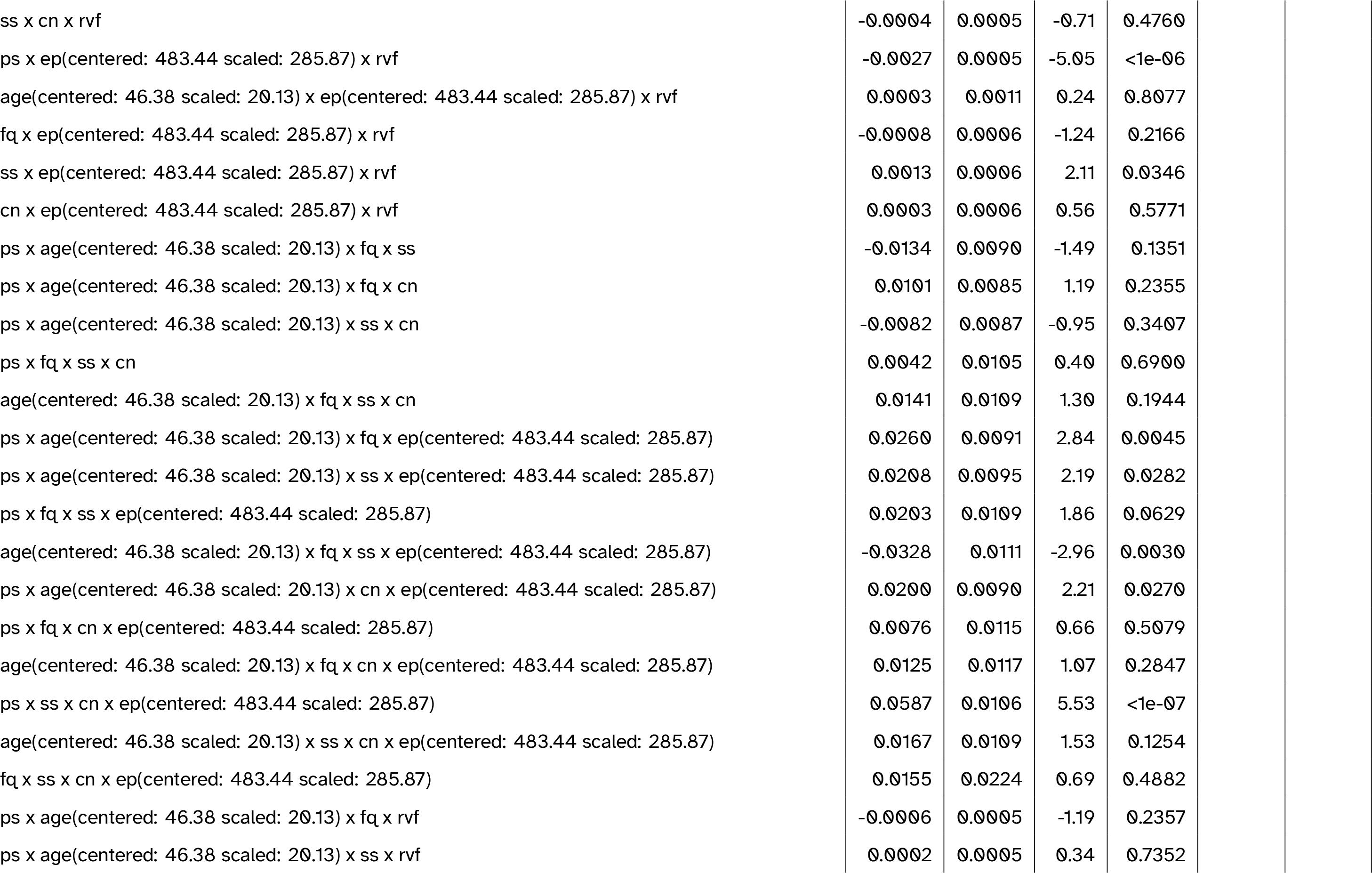
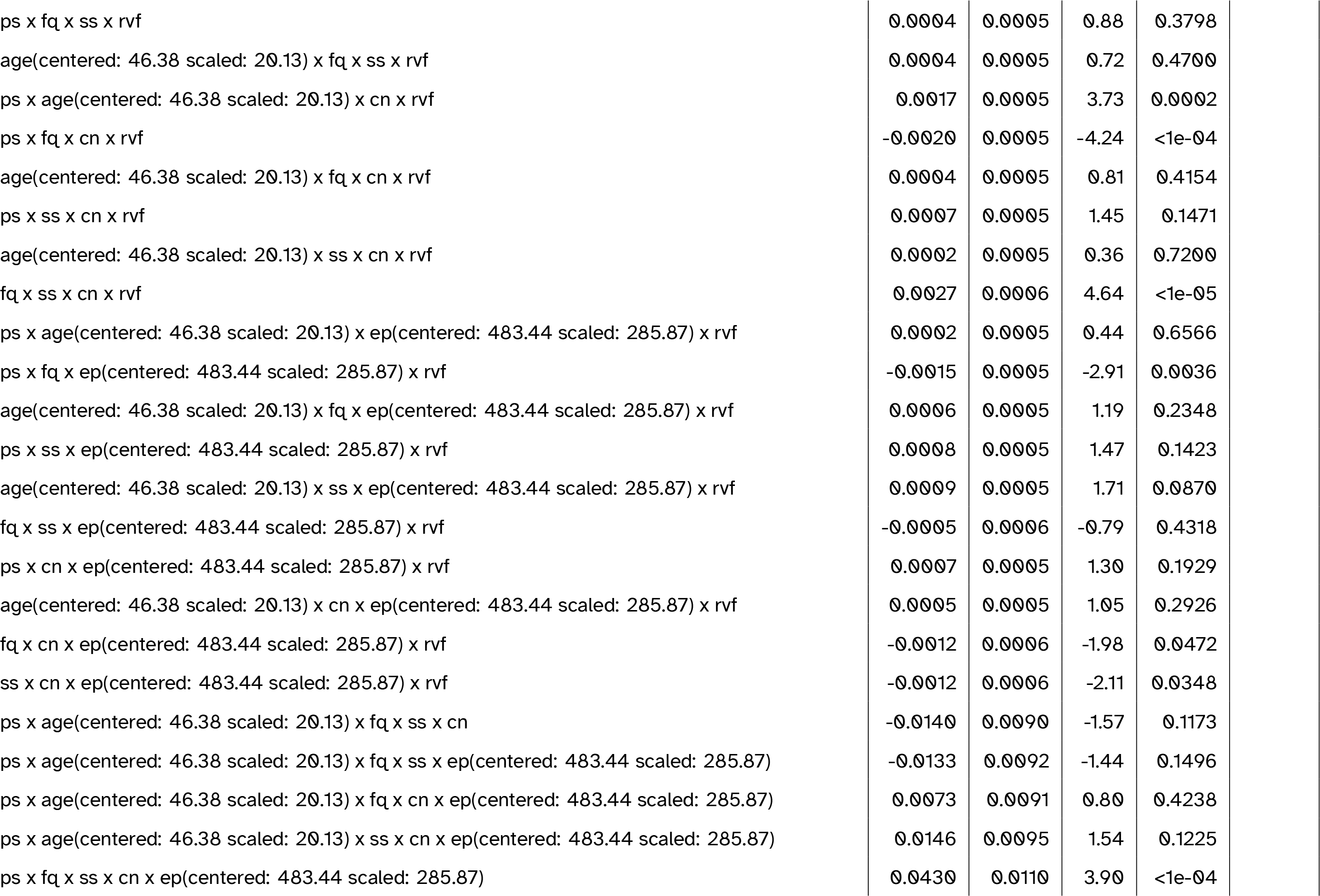
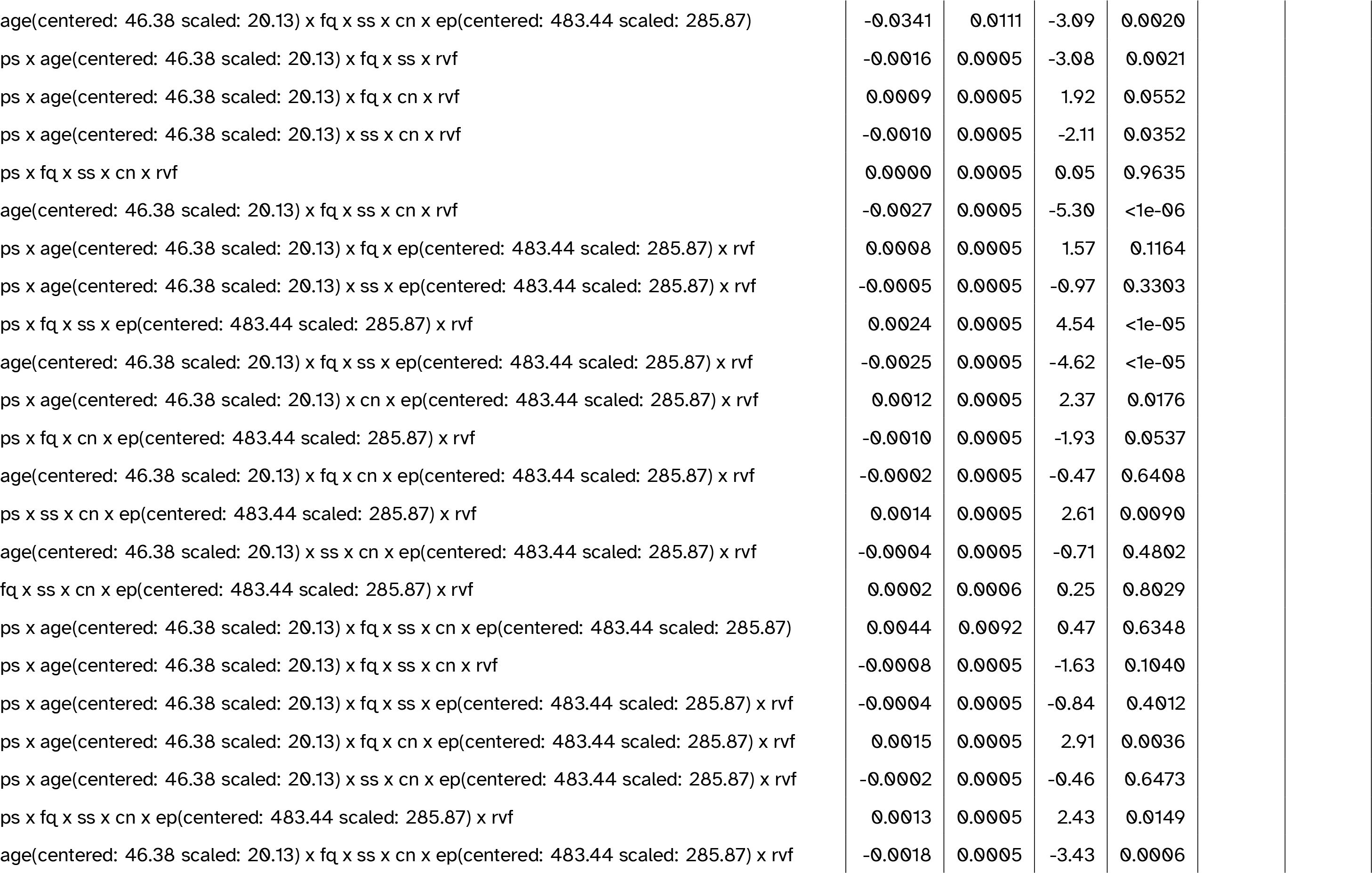
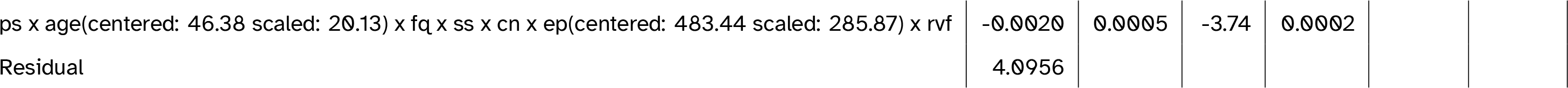
Full model summary for the best-fitting model including verbal fluency (VF). Abbreviations: cn = canonicity; ep = epoch; fq = word frequency; ps = prestimulus amplitude; ps2 = quadratic prestimulus amplitude; rvf = VF (residualised on age); ss = speaker-based surprisal; ss2 = quadratic speaker-based surprisal; ss3 = cubic speaker-based surprisal

**Table 11:**
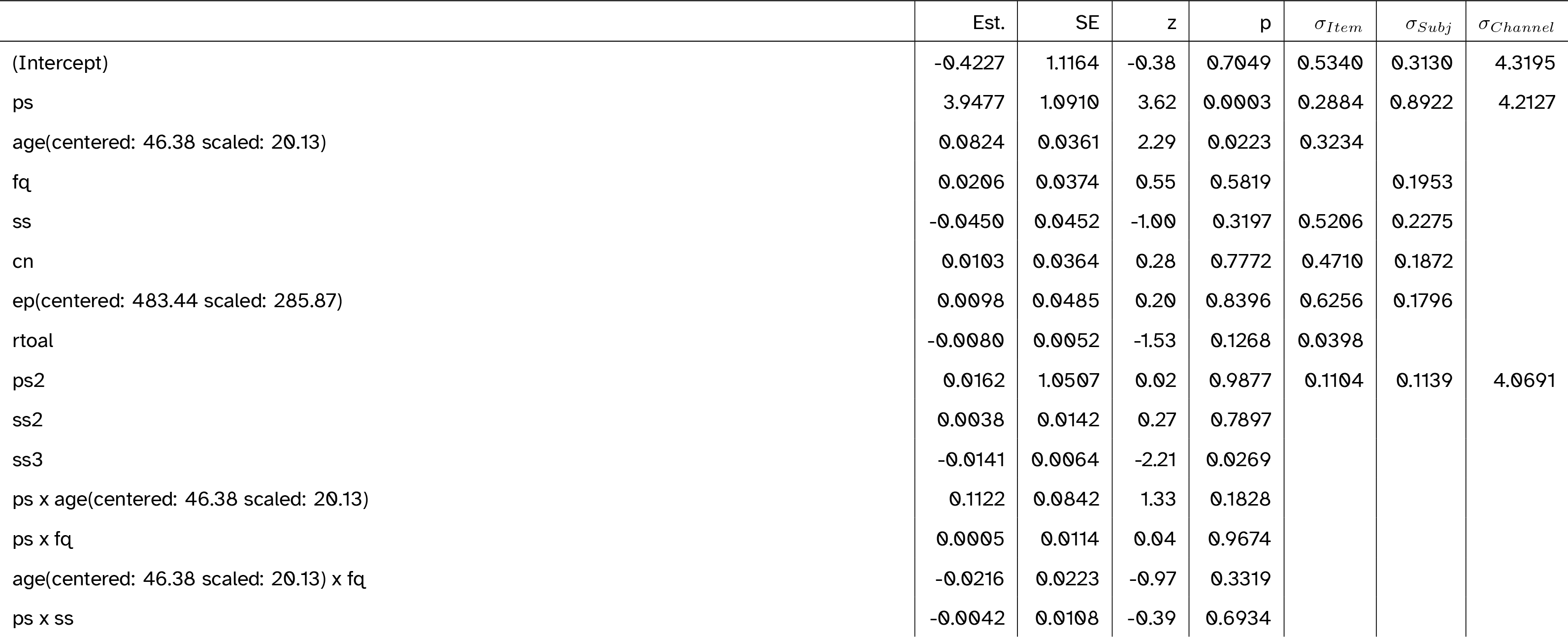
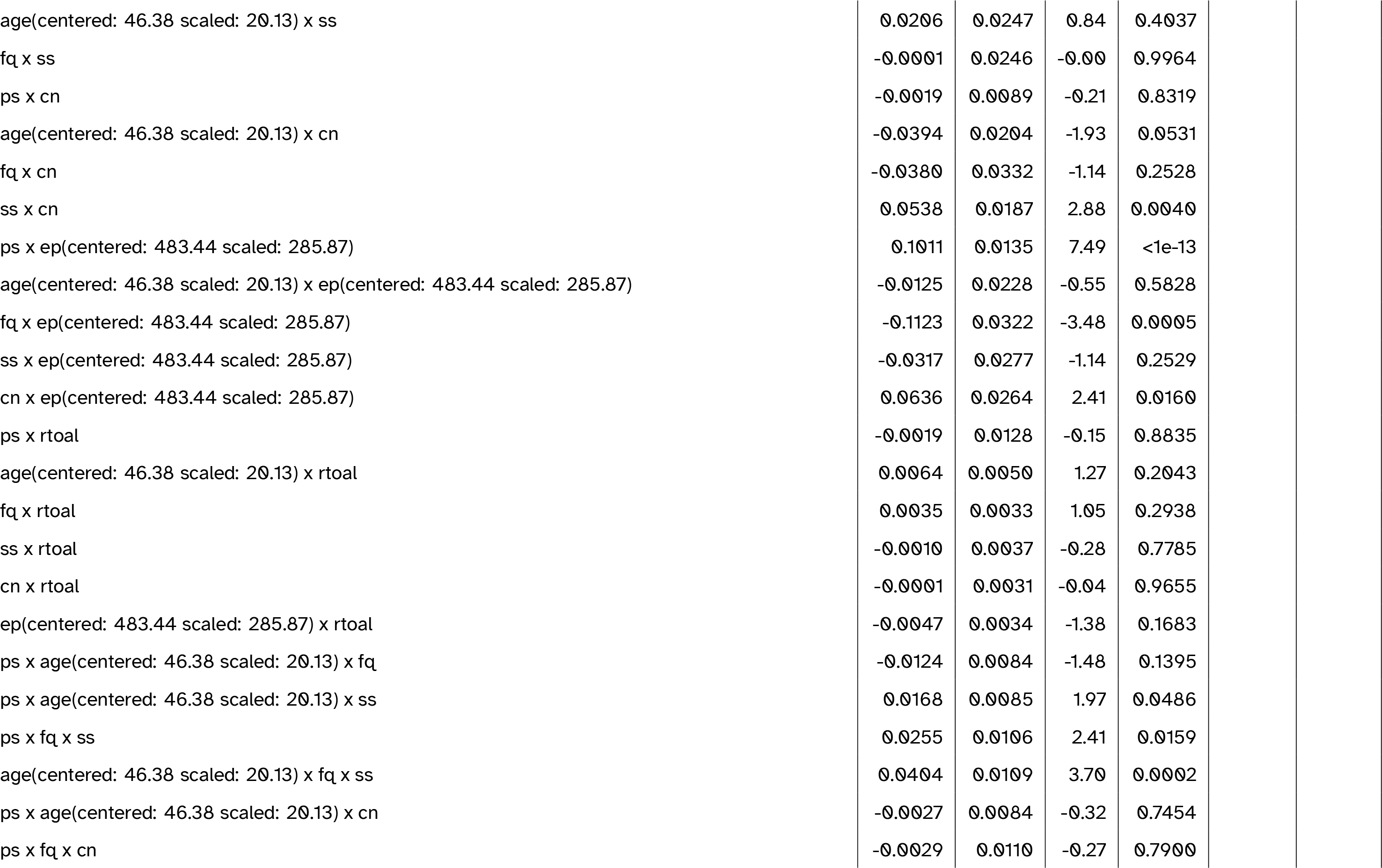
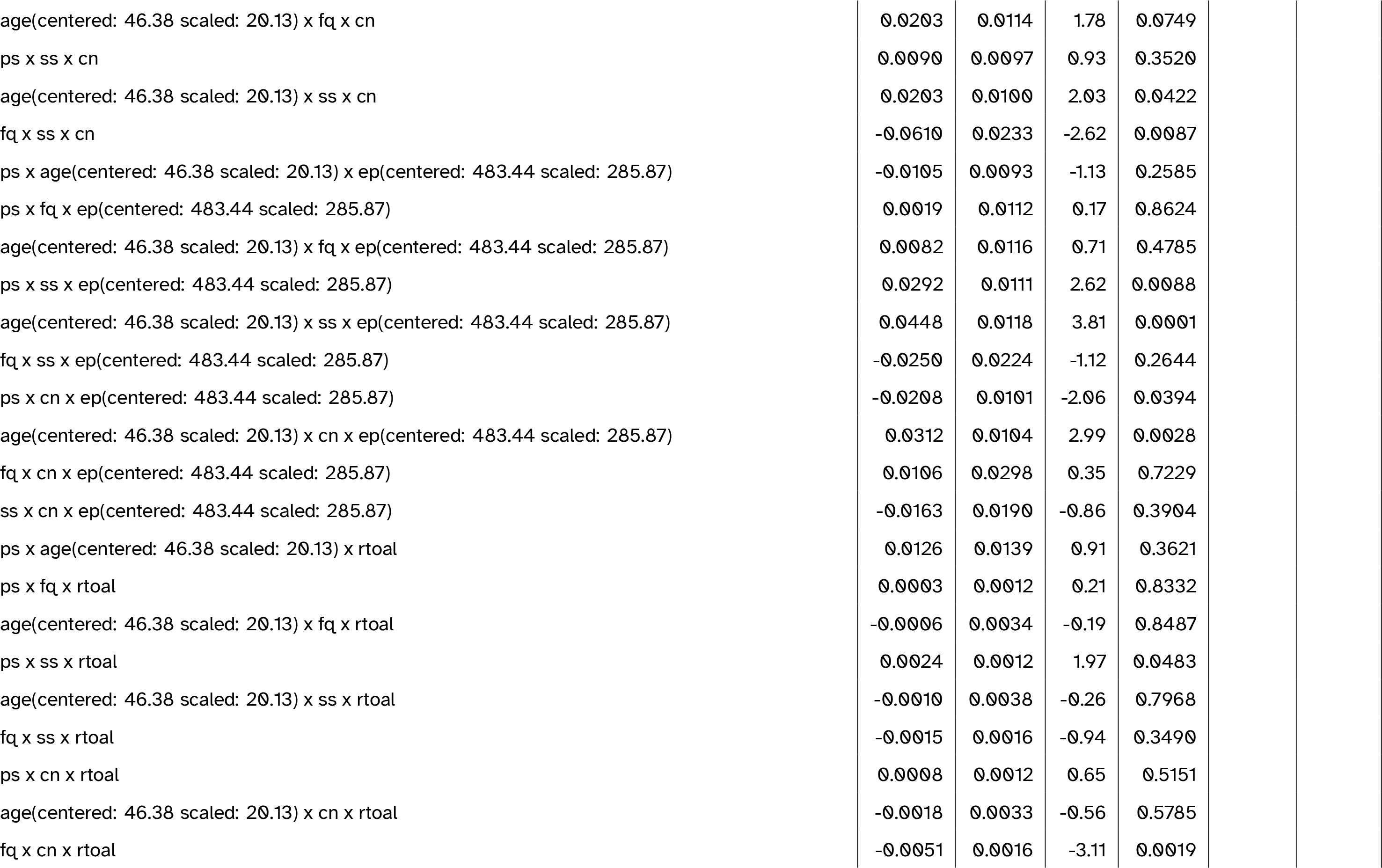
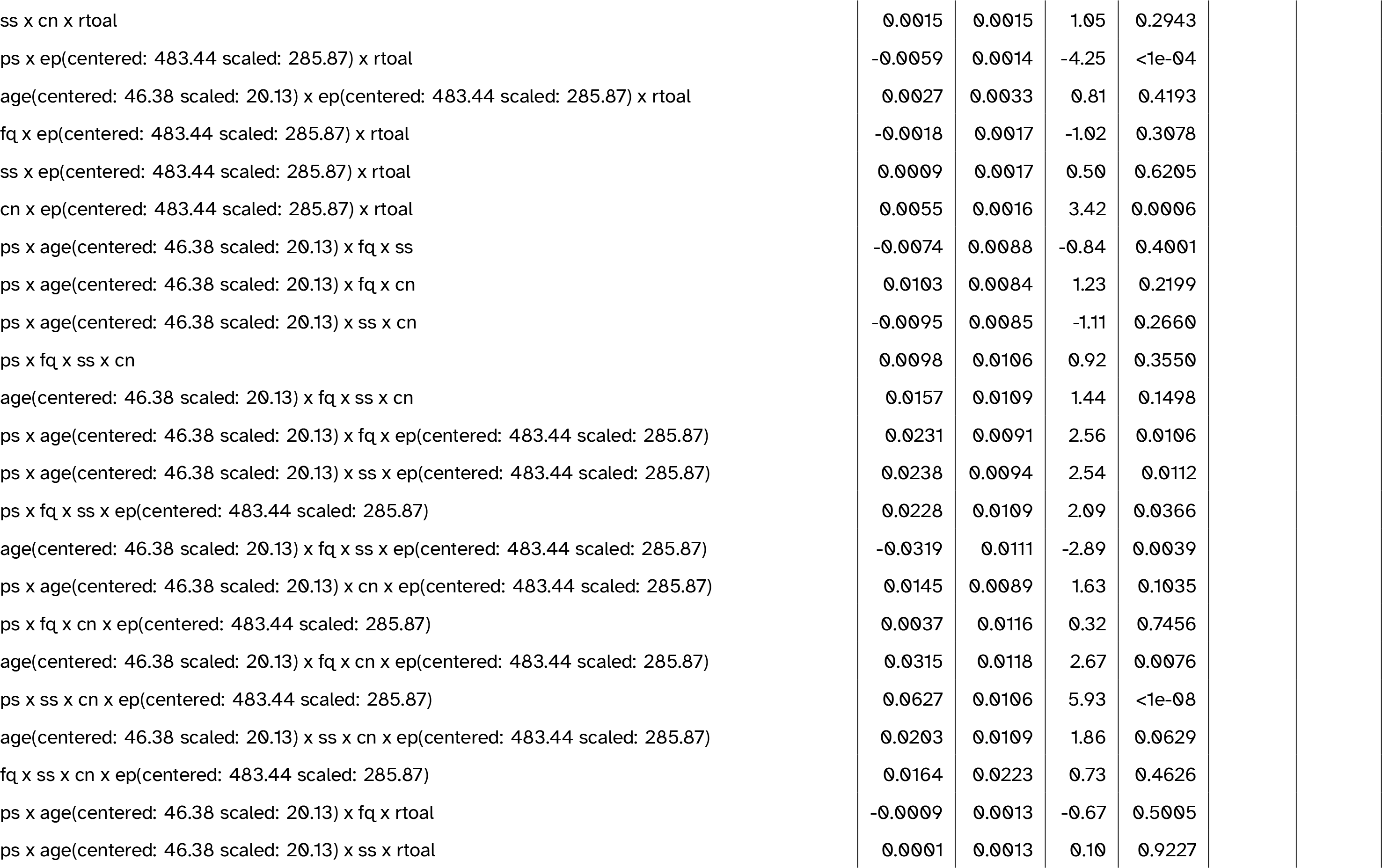
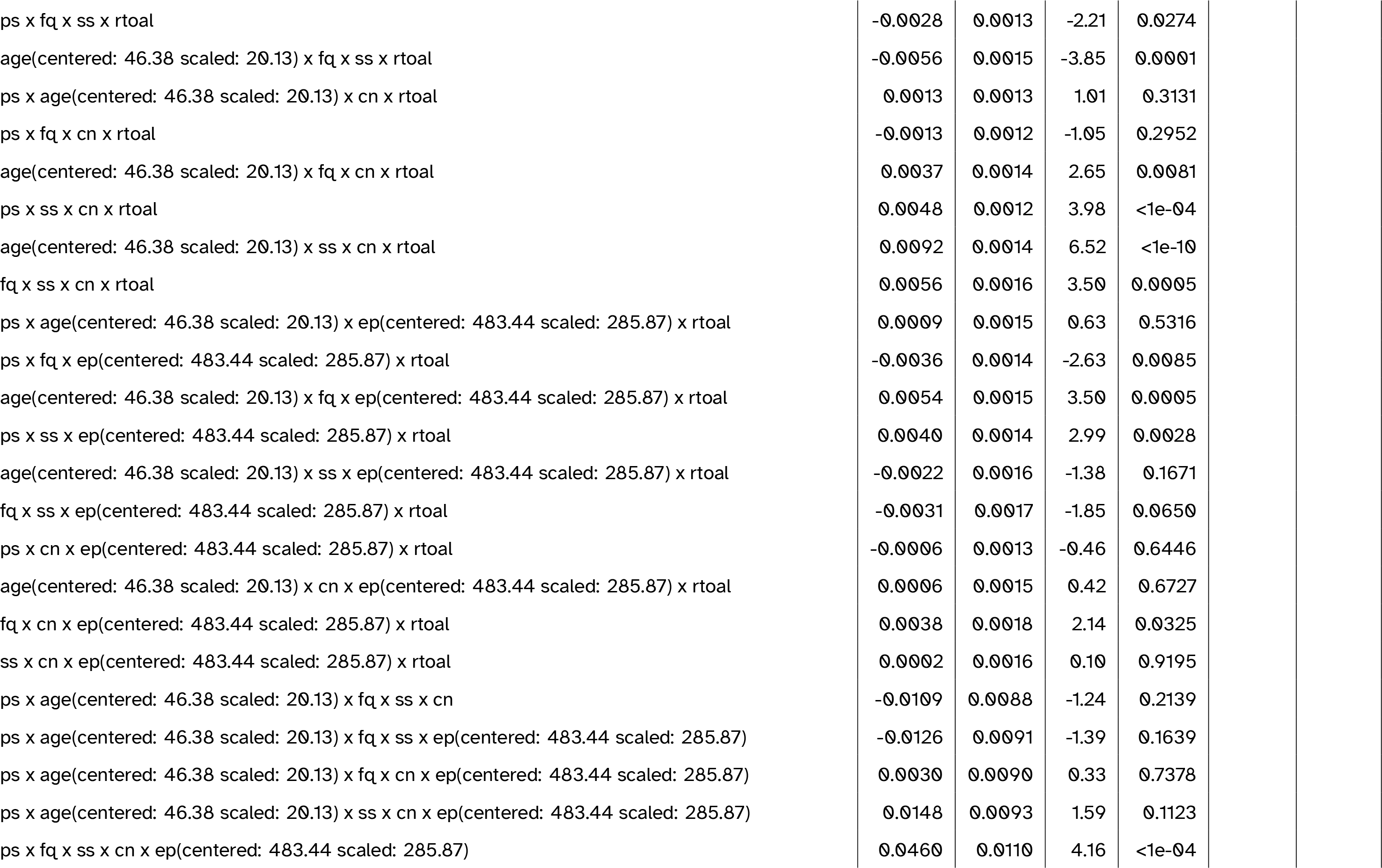
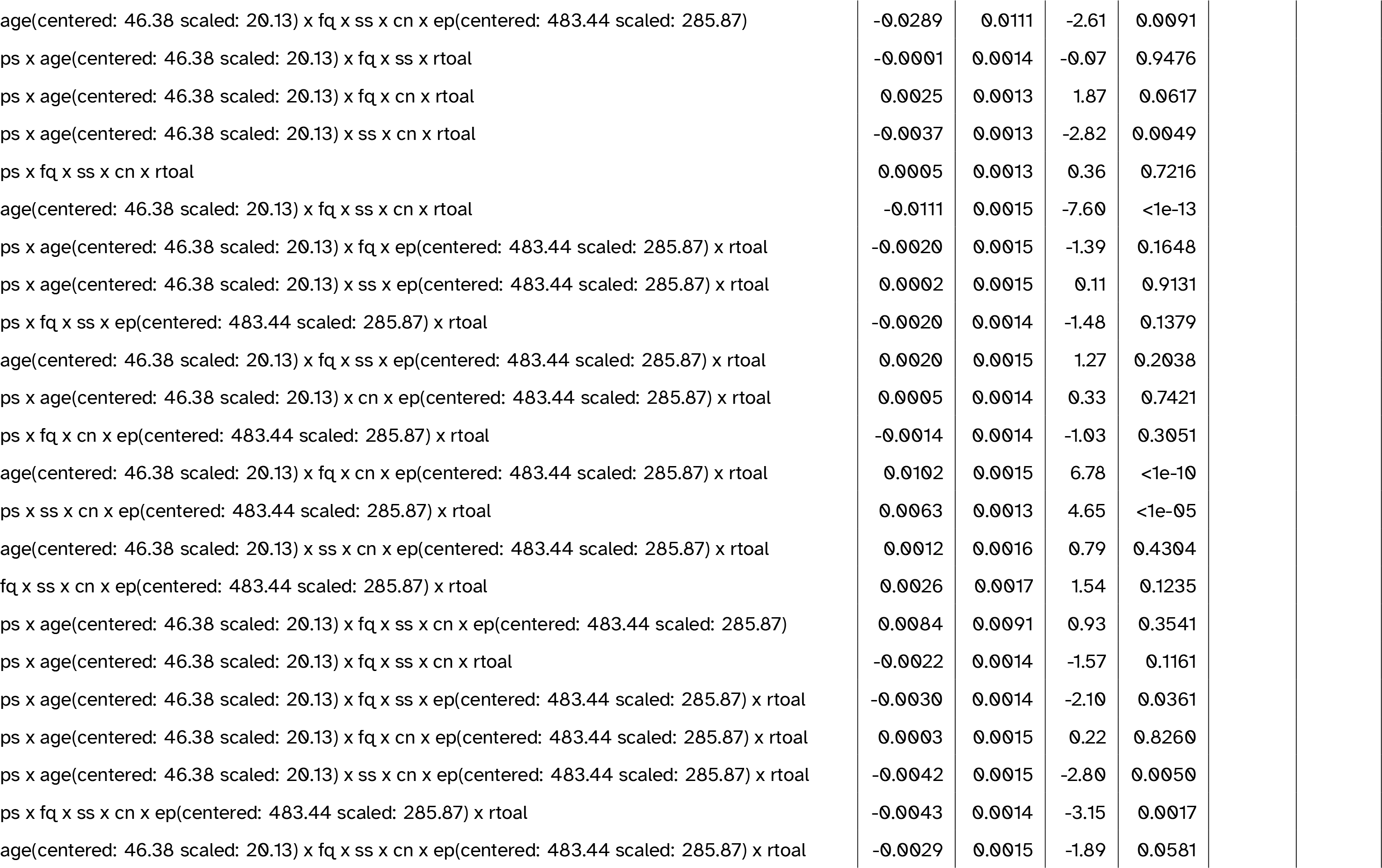
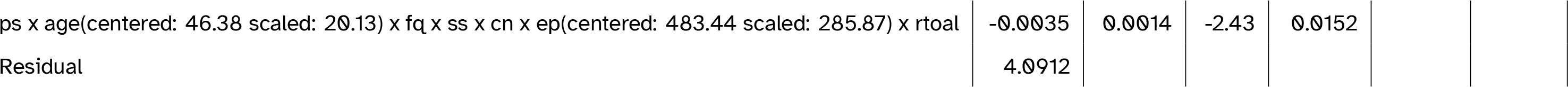
Full model summary for the best-fitting model including TOAL-4 score. Abbreviations: cn = canonicity; ep = epoch; fq = word frequency; ps = prestimulus amplitude; ps2 = quadratic prestimulus amplitude; rtoal = TOAL-4 score (residualised on age); ss = speaker-based surprisal; ss2 = quadratic speaker-based surprisal; ss3 = cubic speaker-based surprisal

**Table 12:**
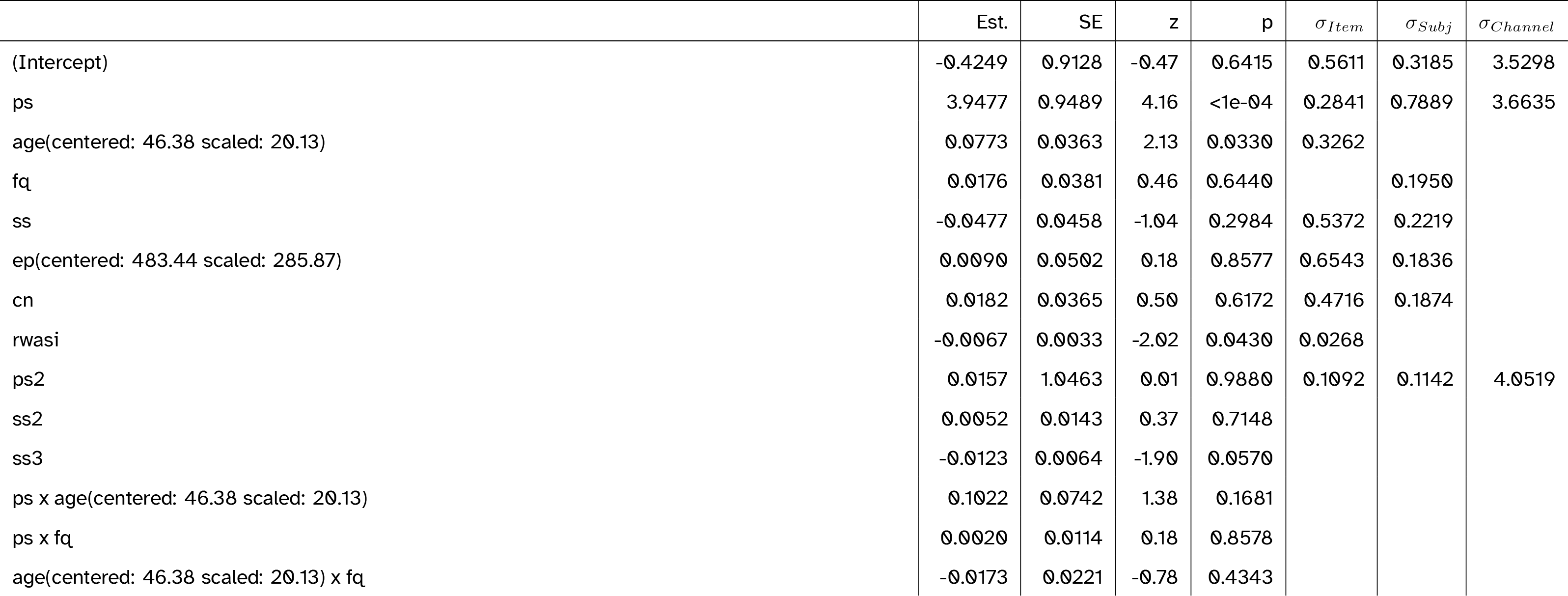
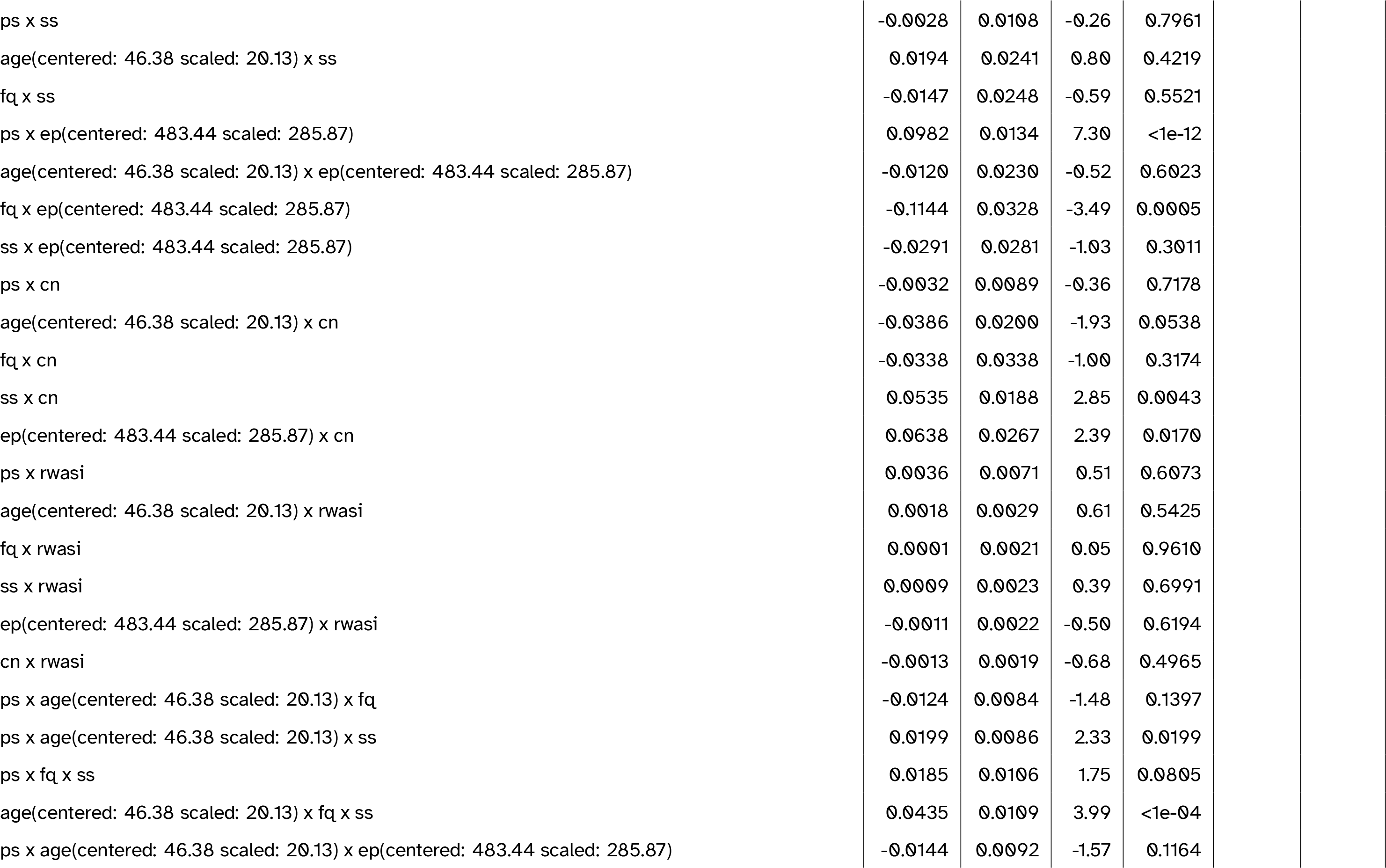
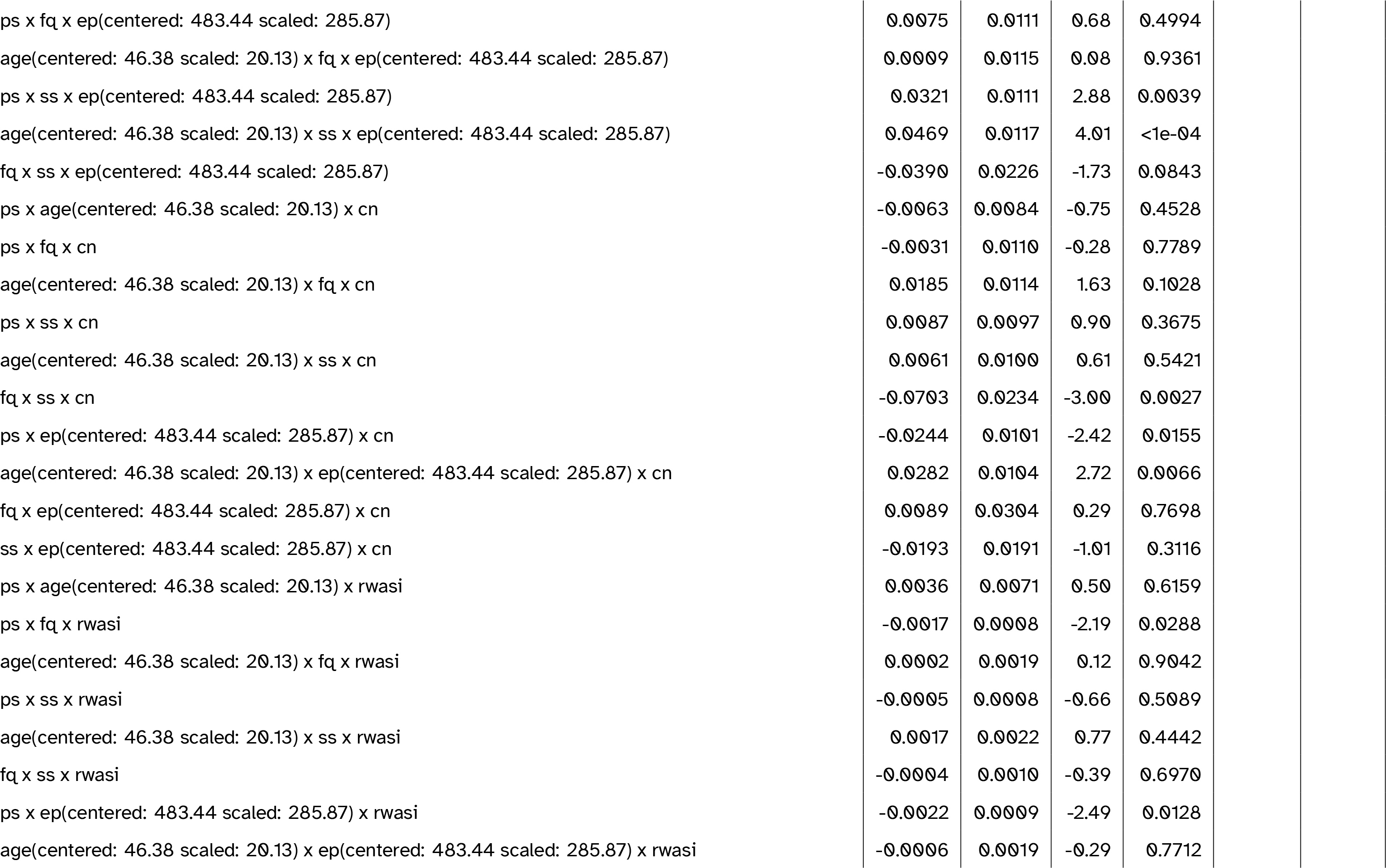
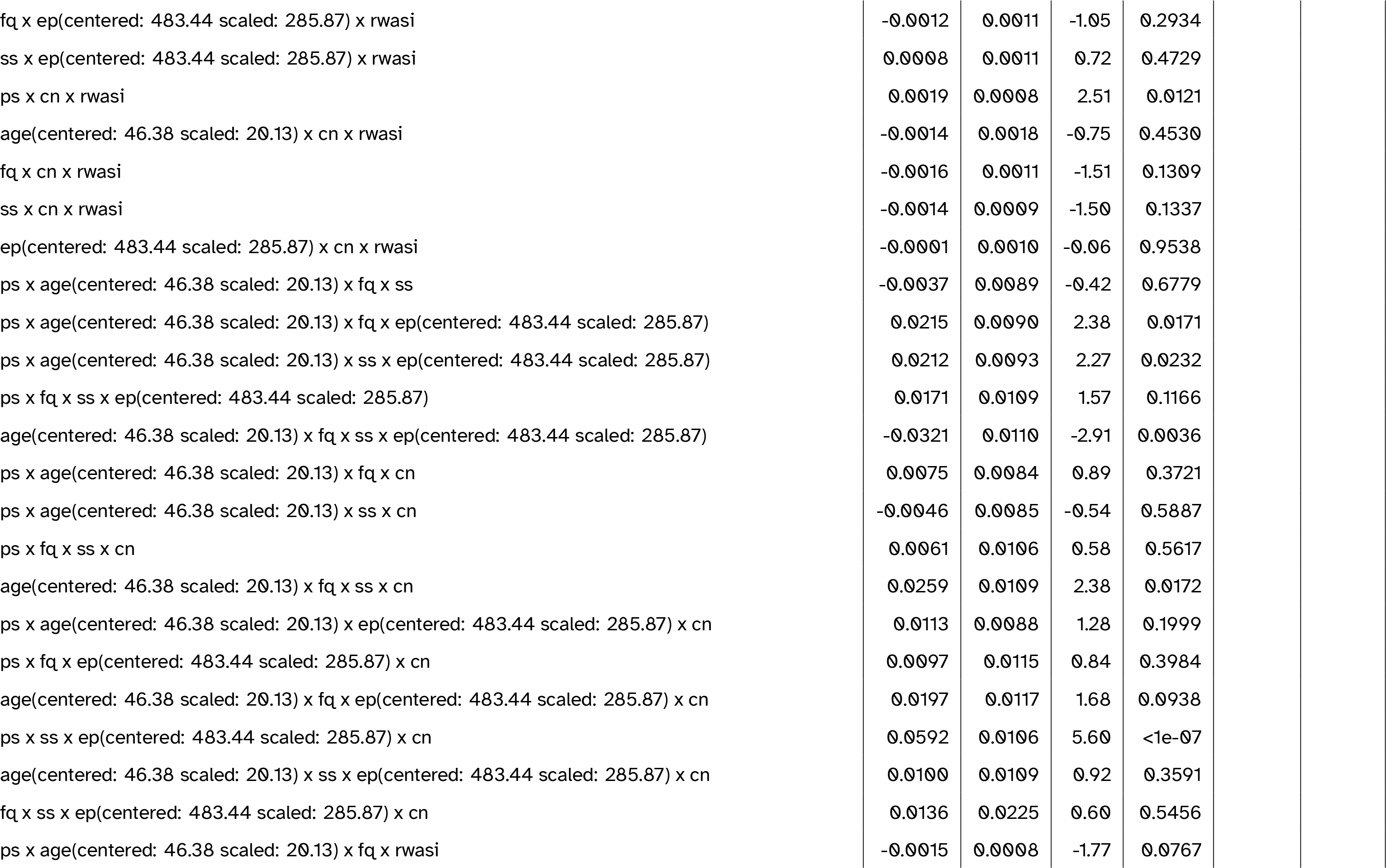
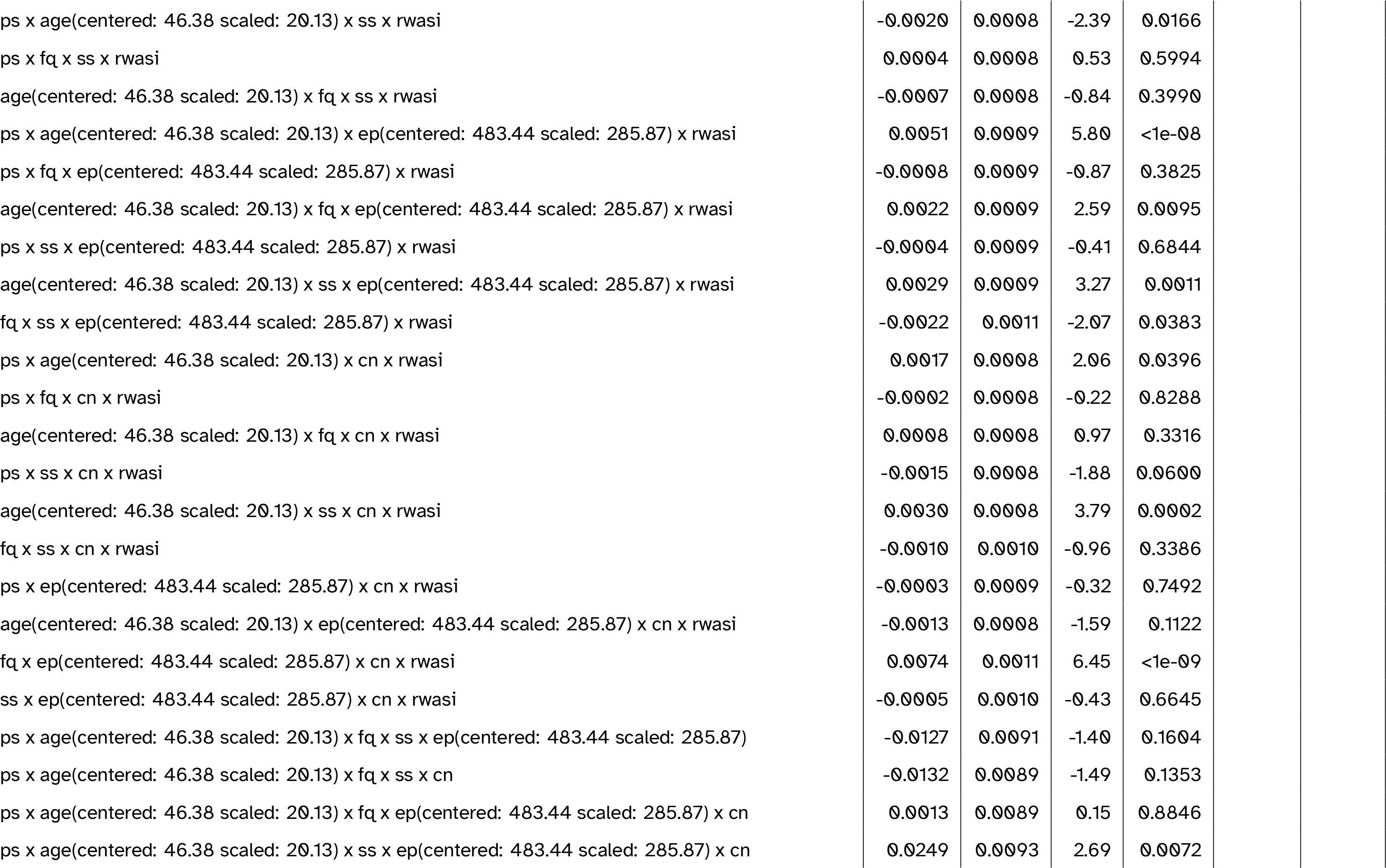
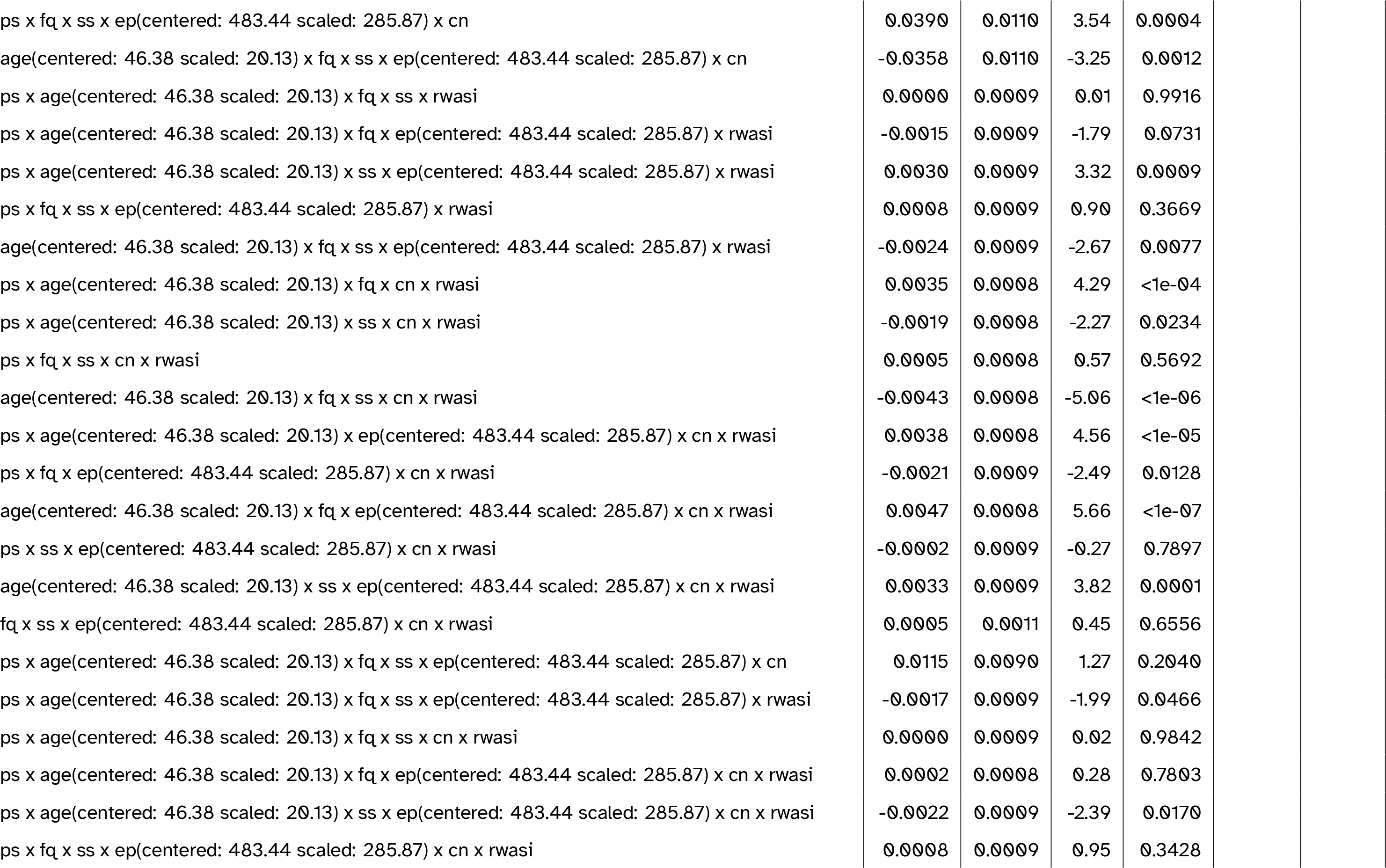
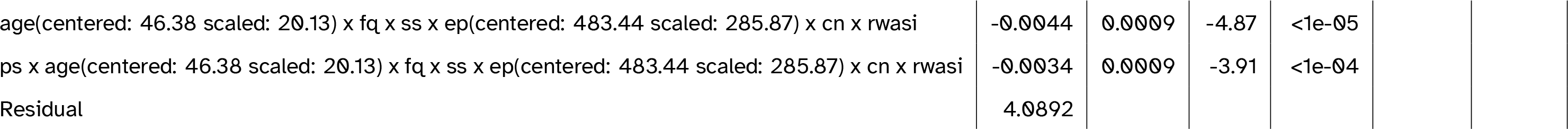
Full model summary for the best-fitting model including general cognitive ability (WASI FSIQ-4). Abbreviations: cn = canonicity; ep = epoch; fq = word frequency; ps = prestimulus amplitude; ps2 = quadratic prestimulus amplitude; rwasi = WASI score (residualised on age); ss = speaker-based surprisal; ss2 = quadratic speaker-based surprisal; ss3 = cubic speaker-based surprisal

## Footnotes

1 Note that, given the passive nature of the oddball task, oddball stimuli were not task-relevant. Their motivational significance was thereby merely a function of their lower probability of occurrence. Importantly, Nieuwenhuis et al. (2005) explicitly extend their noradrenergic theory of P3 generation to both the target-related P3b and the novelty P3a, as observed in passive oddball tasks such as the one employed here.

2 In contrast to the MMN effect for younger and middle-aged adults, older adults show a reduction of MMN amplitude over the course of the experiment. This is compatible with the findings reported by Moran et al. (2014).

